# Sugar Matters: How Protein Glycosylation Shapes Ligand Binding

**DOI:** 10.64898/2026.11.24.768426

**Authors:** Jing Huang, Silvia D’Andrea, Samm Arnold, Ling Han, Duong T. Bui, Elena N. Kitova, Ojas Singh, Lara K. Mahal, Elisa Fadda, John S. Klassen

**Author notes:** Corresponding Author: Department of Chemistry University of Alberta, Edmonton, AB CANADA T6G 2G2. Current address: Department of Chemistry and Department of Biology, University of Waterloo, Waterloo, Ontario, Canada (Duong T. Bui).

## Abstract

Altered protein glycosylation accompanies many diseases and can influence drug binding, yet the contributions of specific glycan features remain difficult to resolve within heterogeneous glycoprotein populations. Here, we combined sequential enzymatic glycan remodeling, composition-resolved variable-temperature native mass spectrometry (vT-nMS), complementary glycosylation analyses, and molecular modelling to investigate warfarin binding to the F1 and S variants of human α1-acid glycoprotein (AGP). Measured affinities spanned more than an order of magnitude across the glycan compositions and remodeled samples examined. Core α1–6 fucosylation favored binding by both variants and shifted the thermodynamics toward less favorable enthalpy and more favorable entropy. Antennary fucosylation generally weakened binding in F1 and had more variable effects in S. Weaker binding was associated with a less favorable entropic contribution despite more favorable enthalpy. Outer-arm trimming produced variant-dependent effects, including changes in enthalpic and entropic contributions that were partly obscured by comparatively small changes in affinity. Endo F3-mediated truncation also produced distinct responses in the two variants. Within the F1 truncation series, the three-core population exhibited the strongest binding and greatest enthalpic stabilization, whereas the corresponding S population exhibited a more favorable entropic contribution than F1 without a comparable affinity enhancement. No warfarin complexes were detected for compositions assigned as containing bisecting GlcNAc. Molecular simulations and reconstructed glycan ensembles suggested that changes in glycan orientation and glycan–protein interactions near the binding-funnel entrance may contribute to the observed binding effects. Together, these findings reveal how glycan structure and protein sequence jointly determine the affinity and thermodynamic balance of ligand recognition and demonstrate the value of composition-resolved vT-nMS for dissecting functional heterogeneity in glycoprotein populations.

TOC graphic

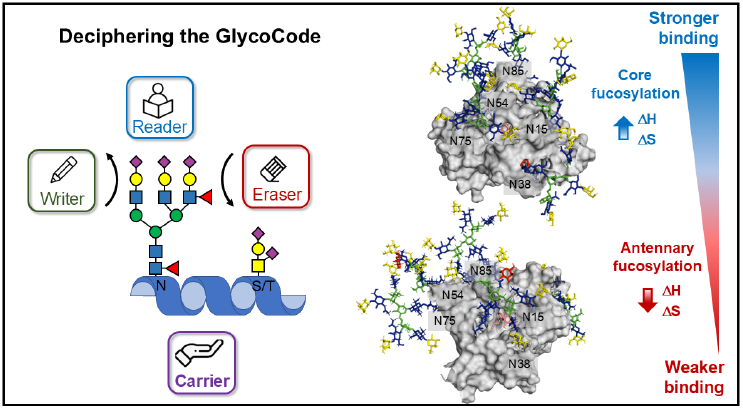

## INTRODUCTION

Glycosylation is one of the most widespread and structurally diverse post-translational modifications of proteins, with an estimated 50% to 70% of human proteins bearing covalently attached glycans (carbohydrates).^1^ Glycan processing is mediated by the coordinated actions of glycosyltransferases and glycoside hydrolases, resulting in a heterogeneous population of glycoforms.^2,3^ The resulting macroheterogeneity (distribution of occupied sites) and microheterogeneity (types of glycans at a given site) can strongly influence protein folding, stability, and molecular recognition.^4–8^

Despite the well-established importance of glycosylation in modulating molecular recognition, quantitative measurements of binding interactions at the level of individual glycoforms remain remarkably limited. This deficiency stems largely from the inherent heterogeneity of glycoproteins, which often exist as complex ensembles of closely related glycoforms that differ in glycan composition, linkage, and branching. Such diversity presents a significant analytical challenge,^8^ as most conventional biophysical techniques, including isothermal titration calorimetry (ITC), surface plasmon resonance (SPR) and nuclear magnetic resonance (NMR) spectroscopy, and analytical ultracentrifugation, rely on bulk measurements that report averaged binding parameters and are, therefore, not readily suited to deconvoluting contributions from individual species within mixtures.^9–16^ As a result, glycoform-specific effects on binding kinetics and thermodynamics are frequently obscured. Given the central role of glycosylation in processes ranging from immune recognition to pathogen–host interactions and therapeutic protein efficacy, there is a pressing need to understand how specific glycoforms modulate molecular recognition. Achieving this requires approaches capable of resolving and quantitatively characterizing interactions at the level of discrete glycoforms, ideally without disrupting their native, heterogeneous distributions.

Native mass spectrometry (nMS) has emerged as a powerful approach for probing protein interactions under near-native conditions, without the need for labeling or immobilization.^17,18^ In the context of glycoproteins, nMS is particularly well suited to resolving intact species that differ in overall glycan composition, enabling the direct observation of heterogeneous protein populations.^19–23^ It can also simultaneously quantify protein–ligand interactions across coexisting glycan-defined species within a single experiment, yielding composition-resolved binding stoichiometries and affinities.^18^ Although each measured mass may represent an ensemble of isomeric and site-specific glycoforms, this level of compositional resolution provides a practical and experimentally accessible means of dissecting glycosylation heterogeneity. To further refine these measurements, enzymatic glycan remodeling strategies can be employed to systematically simplify or redefine glycan compositions, thereby reducing heterogeneity and enabling clearer structure–function relationships to be established.^22,24–26^ In parallel, liquid chromatography– tandem MS (LC–MS/MS)-based glycopeptide analysis provides site-specific structural assignments of glycans, though often in a semi-quantitative manner.^27,28^ When integrated, these approaches offer a comprehensive view of glycoprotein binding behavior, linking detailed glycan structural information to functional, composition-resolved binding measurements obtained under native-like conditions.

Despite this potential, applications of nMS-based strategies to glycoprotein–ligand interactions remain scarce, with only a small number of systems in which glycan heterogeneity has been directly linked to differences in binding behaviour.^29,30^ The most detailed work to date is from Robinson and coworkers, who leveraged high-resolution nMS to quantitatively investigate interactions involving the asialo forms of two *N*-glycosylated human plasma glycoproteins: α1-acid glycoprotein (AGP), a monomeric acute-phase protein involved in immunomodulation and drug binding, and haptoglobin (Hp), a tetrameric (α₂β₂) complex composed of two α–β heterodimers linked by disulfide bonds that binds and clears hemoglobin (Hb) released during hemolysis. ^29^ In this work, the influence of binding of the anticoagulant warfarin to asialo-AGP and binding of Hb to Hp was examined in the context of complex type *N*-glycosylation. Notably, the nMS measurements showed that the interactions depend on *N*-glycan branching and fucosylation. Increased *N*-glycan branching was found to attenuate binding in both systems, possibly through steric hindrance, while fucosylation weakened warfarin binding but stabilized the Hp–Hb interactions. More recently, it was reported that *N*-glycosylation of the SARS-CoV-2 receptor-binding domain (RBD) regulates sialoglycan binding.^31^ Specifically, the loss or truncation of the RBD *N*-glycan at Asn343 resulted in the near complete loss of GM1 and GM2 ganglioside oligosaccharide binding due to loss of protein structural stability in the sialoglycan binding site.^31^

Together, these and related studies establish integrative nMS strategies as a powerful approach for uncovering glycosylation-dependent effects that are otherwise obscured in ensemble measurements. However, while informative, variation in *K*_d_ alone provides limited mechanistic insight, as it often reflects compensating enthalpic and entropic contributions. We therefore hypothesize that resolving the underlying thermodynamic signatures of binding, specifically the enthalpic (Δ*H*) and entropic (Δ*S*) contributions, will provide deeper insight into the molecular origins of glycosylation-dependent interactions. To further contextualize these effects at the molecular level, such measurements can be complemented by molecular dynamics (MD) simulations, assisting the interpretation of the observed trends with structural and dynamic insight at the atomistic scale.

Here, we test this hypothesis using the AGP–warfarin complex system to characterize the thermodynamic origin behind the observed glycosylation-dependence of its binding affinity. Previous nMS, ITC, and MD studies have shown that AGP glycosylation alters ligand affinity, with proposed contributions from steric effects, conformational dynamics, and changes in the physicochemical environment of the binding cavity.^29,32–34^ However, these relationships remain largely correlative, and the enthalpic and entropic contributions of specific glycan features have not been resolved. To address this gap, we used sequential enzymatic remodeling to generate and characterize AGP samples with progressively simplified *N*-glycans and then applied composition-resolved variable-temperature (vT)-nMS to quantify glycan-dependent changes in warfarin affinity and thermodynamics. Complementary MD simulations and reconstruction of heterogeneous glycoform ensembles provide an atomistic framework for interpreting the experimental findings in terms of glycan conformational sampling and accessibility of the binding cavity. Together, this integrated approach establishes quantitative structure–thermodynamics relationships between *N*-glycan features and ligand recognition, moving beyond affinity alone toward a mechanistic understanding of glycosylation-dependent molecular recognition.

## MATERIALS AND METHODS

### Materials

Human plasma α-1 acid glycoprotein (AGP), neuraminidase from *Clostridium perfringens* (NanI subtype, denoted as NeuC), PNGase F, and warfarin were purchased from Sigma-Aldrich Canada (Oakville, Canada). α1-3,4 fucosidase, β1-4 galactosidase S, β-N-acetylglucosaminidase S and Endo F3 were purchased from New England Biolabs Canada (Whitby, Canada). Protein stock solutions were dialyzed into aqueous 200 mM ammonium acetate (at the required pH) using Amicon 0.5 mL microconcentrators (EMD Millipore, Billerica, MA) with a MW cut-off of 10 kDa. Protein concentrations were estimated by UV absorption at 280 nm. All stock solutions were stored at −20 °C until used.

### Other reagents

Other solvents and reagents were purchased from common commercial sources and were used without further purification.

### Separation of AGP variants

The method used to separate the human AGP F1 and S variants from the A variant by immobilized metal ion-affinity chromatography (IMAC) was described previously by Hervé *et al*.^35^ Experimental details are given as Supporting Information.

### Isothermal Titration Calorimetry (ITC)

The ITC measurements of the AGP-warfarin and asialo-AGP-warfarin binding affinity were carried out using a Microcal PEAQ ITC (Malvern Panalytical, Worcestershire, United Kingdom). Experimental details are given as Supporting Information.

### Native MS

All nMS binding measurements were performed in positive ion mode using a Q Exactive Ultra-High Mass Range (UHMR) Orbitrap mass spectrometer (Thermo Fisher Scientific, Bremen, Germany) equipped with a home-built temperature controlled nanoESI source.^36^ Details of the experimental and instrumental parameters used for the vT-nMS binding measurements and the data analysis procedures are provided as Supporting Information.

### NanoLC-MS/MS analysis

Glycopeptide analysis was performed using a Waters NanoACQUITY UPLC system (Waters, Milford, MA) coupled with 15T solariX-XR Fourier-transform ion cyclotron resonance mass spectrometer (Bruker-Daltonics, Billerica, MA). Released *N*-glycans analysis was performed using a Thermo Scientific™ Vanquish™ UHPLC system coupled with a fluorescence detector (FLD) (Thermo Scientific, Waltham, MA, USA) and ESI-MS (Thermo Q Exactive Orbitrap, Thermo Fisher Scientific, Bremen, Germany). Experimental details and instrumental parameters used and data analysis procedures are provided as Supporting Information.

### Computational Methods

#### MD simulations

AGP glycoforms were generated using the ReGlyco tool in GlycoShape (https://glycoshape.org).^37^ The simulation box was solvated with TIP3P water^38^ and 150 mM of NaCl. The protein and counterions, and glycan atoms were represented with the Amberff14SB^39^ and GLYCAM06j-1,^40^ respectively. Gaussian accelerated molecular dynamics (GaMD) simulations^41^ were performed using the implementation in AMBER24^42^ in dual-boost mode, with independent boost potentials applied to the dihedral and total potential energies. Protein stability and local conformational dynamics were assessed using backbone root mean square deviation (RMSD) and Cα root mean square fluctuation (RMSF) values calculated with scripts run in the Visual Molecular Dynamics (VMD) Tcl console.^43^

#### Conformational ensemble analysis of AGP glycoforms

For this analysis we selected a group of AGP glycoforms (Table S1) that recapitulates the main features of the experimentally characterised ensembles, namely a reference glycoform, core-fucosylated, and bisected glycoforms, together with a systematic panel of antenna-fucosylated glycoforms differing in glycan branching, glycosylation site, and the specific antenna functionalized. The antenna-fucosylated systems are indicated as Models 1–12 with glycan compositions shown in Table S1. All glycoforms were rebuilt with the ReGlyco Ensemble tool in GlycoShape using 200 frames for each selected glycan at each site selected from the MD database. Glycan–protein interactions were quantified using the Interactions plugin in VMD.^43^

## RESULTS AND DISCUSSION

Human AGP is a 183 amino acid plasma lipocalin with a MW of approximately 22 kDa for the polypeptide backbone and ∼40 kDa in its fully glycosylated form. AGP adopts the canonical lipocalin fold, consisting of an eight-stranded antiparallel β-barrel enclosing a deep internal binding cavity.^32^ This cavity is divided into three interconnected lobes (I–III), with lobe I forming the largest and deepest hydrophobic chamber, whereas lobes II and III are smaller and enriched in negatively charged residues. Structurally diverse endogenous metabolites and basic or neutral drugs occupy partially overlapping regions of the cavity. Negatively charged warfarin binds predominantly within the hydrophobic lobe I (Figure 1).^34^ The β-barrel is flanked by several α-helices and surface loops, while two conserved disulfide bonds (Cys5–Cys147 and Cys72– Cys165) stabilize the overall fold.^44^ Human AGP exists as several genetic variants, including the ORM1-derived F1 and S variants, which differ by a single amino acid substitution (Q20R). For this study, the F1 and S variants were isolated as a mixture by metal-ion affinity chromatography from a commercial AGP preparation derived from pooled human plasma.^35^ Both variants contain five occupied *N*-glycosylation sites (Asn15, Asn38, Asn54, Asn75, and Asn85; mature-protein numbering) bearing branched, complex-type *N*-glycans with extensive sialylation and variable fucosylation. These sites are distributed across the solvent-exposed protein surface, with Asn75 positioned closest to the cavity entrance and Asn38 located on the major α-helix adjacent to the β-barrel (Figure 1).^34^ Thus, although the glycans are unlikely to contact a cavity-bound ligand directly, their distribution across the protein surface provides multiple routes through which glycosylation could modulate ligand recognition, including changes in protein conformation and dynamics, local hydration, and steric accessibility to the binding cavity.

**Figure 1.**
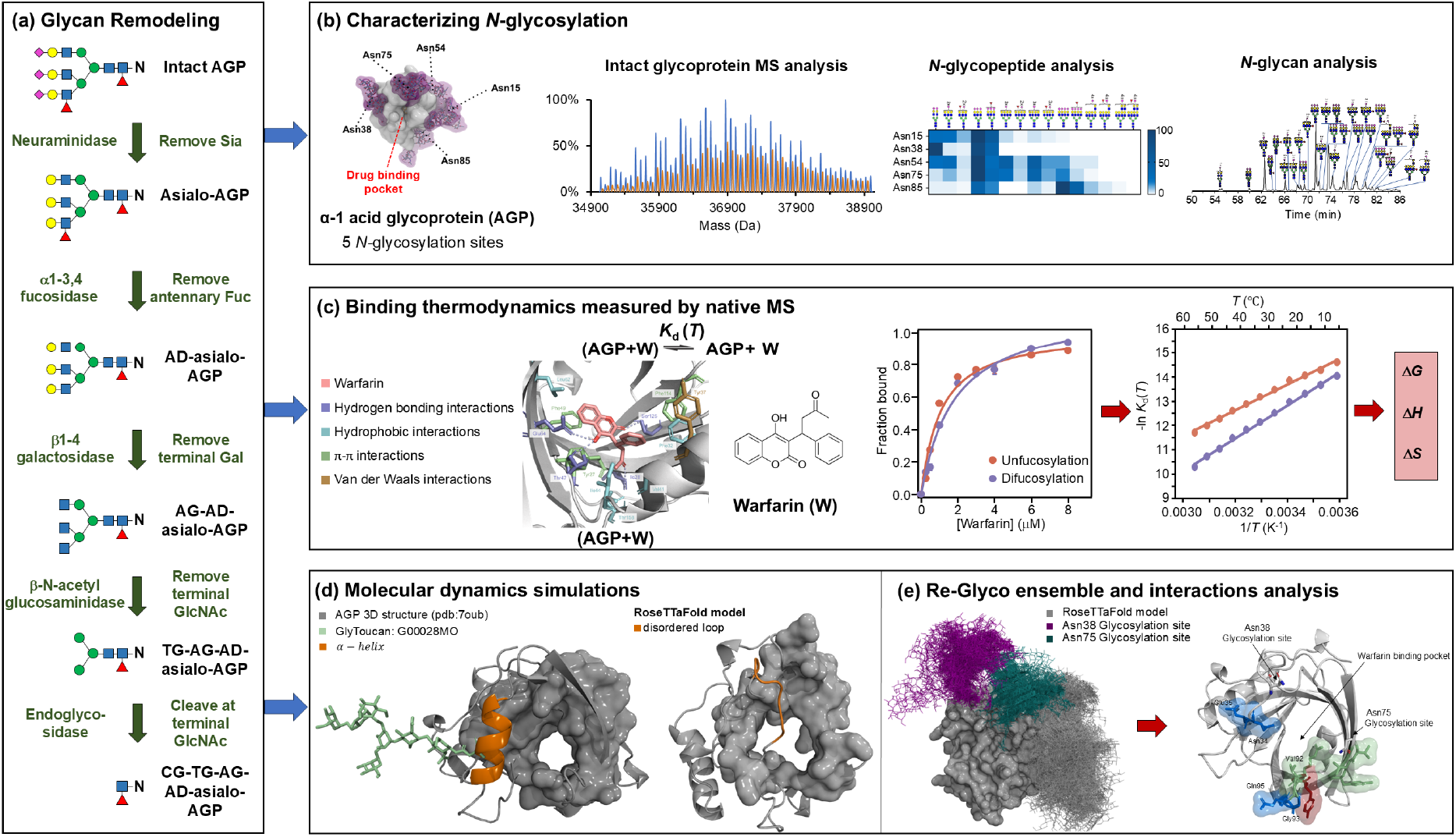
Integrated workflow for elucidating glycosylation-dependent α1-acid glycoprotein (AGP)–warfarin (W) interactions. (a) Sequential glycosidase treatments were used to systematically remodel the AGP *N*-glycans. (b) AGP microheterogeneity was characterized by intact glycoprotein MS, site-specific *N*-glycopeptide analysis, and released *N*-glycan analysis. The *N*-glycosylation sites are indicated on the AGP-F1 structure, together with the predicted binding pose of warfarin in the binding pocket. (c) Native MS binding measurements performed over a range of temperatures were used to quantify glycan composition-dependent warfarin binding and determine the corresponding thermodynamic parameters. (d) MD simulations provided molecular-level insight into the experimentally observed effects of glycosylation on warfarin binding. (e) GlycoShape Re-Glyco Ensemble was used to generate conformational ensembles of representative AGP F1 glycoforms and characterize glycan–protein interactions around the binding-pocket entrance.

To define how *N*-glycosylation regulates ligand binding to AGP, we established an integrated workflow combining enzymatic glycan remodeling, structural characterization of glycosylation, nMS binding measurements, and MD simulations (Figure 1). A panel of AGP samples with progressively truncated *N*-glycans was generated through sequential exo- and endoglycosidase treatments. The resulting structures ranged from extensively sialylated complex-type *N*-glycans to single core GlcNAc residues at occupied sites, providing a controlled platform for evaluating how specific *N*-glycan features influence ligand binding. The glycosylation status of each sample was characterized by intact MS, released *N*-glycan analysis, and site-specific glycoproteomics. Direct nMS measurements quantified glycan-dependent differences in warfarin affinity, and vT-nMS resolved the corresponding enthalpic and entropic contributions. Molecular simulations and reconstructed glycan ensembles were used to explore possible structural explanations for the observed binding trends.

### Enzymatically treated AGP F1/S variant panel for binding studies

#### Untreated AGP

Analysis of untreated AGP is complicated by extensive glycan microheterogeneity and the nearly identical mass contributions of two Fuc (292.29 Da) and one Neu5Ac (291.26 Da), which preclude unambiguous compositional assignments from intact-protein masses alone (Figure S1, Table S2). Nevertheless, spectral deconvolution resolved the F1 and S variants and revealed broad compositional distributions for each. Within each variant, species sharing a common Hex/HexNAc composition were grouped into families P1–P9, with adjacent families separated by one Hex–HexNAc unit and increasing P number corresponding primarily to greater overall glycan branching. Each family contained multiple species differing in total Fuc and Neu5Ac content. Released *N*-glycan analysis revealed predominantly sialylated, variably fucosylated tri- and tetra-antennary complex-type structures, together with a minor population of elongated tetra-antennary glycans (Figure S2a). Site-specific analysis showed predominantly tri-antennary glycans at Asn15, bi- and tri-antennary glycans at Asn38, and tri- and tetra-antennary glycans at Asn54, Asn75, and Asn85 (Figure S2b).

#### Asialo-AGP

To reduce the compositional heterogeneity of untreated AGP, Neu5Ac was removed using the broad-specificity neuraminidase NeuC. Desialylation substantially simplified the intact mass spectrum, resolving peak series for the F1 and S variants, separated by 28 Da, and enabling assignment of more than 100 glycoprotein compositions (Figure S3a, Table S3). The compositions retained the P1–P9 family classification established for untreated AGP, with each family resolved into five fucosylation states (F0–F4) corresponding to zero to four Fuc residues. Notably, the two variants exhibited indistinguishable glycosylation profiles. In the more abundant F1 variant, additional species consistent with bisecting GlcNAc were distinguished from partially overlapping F4 compositions by accurate mass and represented approximately 25–30% of the corresponding families. Released *N*-glycan and site-specific glycopeptide analyses confirmed complete desialylation (Figures S3b and S4a). Glycan branching varied among the five sites, whereas fucosylation showed no significant site dependence. Glycans containing bisecting GlcNAc were detected at low abundance in the released *N*-glycan profile but not by glycopeptide analysis.

#### Antennary defucosylated (AD)-asialo-AGP

Asialo-AGP was treated with an α1–3,4 fucosidase to selectively remove antennary Fuc residues. Intact MS confirmed efficient defucosylation, with the relative abundance of fucosylated glycoforms decreasing from approximately 55% to 12% following treatment (Figure S3c, Table S4). Glycopeptide analysis localized the remaining fucosylated glycans predominantly to Asn38 and Asn75 (Figure S3d). Although glycopeptide analysis alone could not distinguish core from antennary fucosylation, the linkage specificity of the α1–3,4 fucosidase, together with released *N*-glycan analysis (Figure S4b), indicated that the residual Fuc was core α1–6-linked. This residual core fucosylation was retained during subsequent glycan-remodeling steps.

#### Agalactosylated (AG)-AD-asialo-AGP

Treatment of AD-asialo-AGP with exo-β-galactosidase extensively removed terminal Gal, reducing both molecular mass and compositional heterogeneity (Figure S3e, Table S5). The resulting compositions formed nine families (P′1–P′9), each containing the 15 Hex residues expected from five Man_3_GlcNAc_2_ cores but differing in total HexNAc content. The low-abundance P′7–P′9 families were excluded from subsequent binding and thermodynamic analyses. Site-specific glycopeptide and released *N*-glycan analyses confirmed extensive degalactosylation across all five sites and among bi-, tri-, and tetra-antennary glycans (Figures S3f and S4c). The small residual population of Gal-containing glycans likely contained extended LacNAc chains in which internal Gal residues were protected from exo-β-galactosidase cleavage by terminal GlcNAc.

#### Terminal GlcNAc-removed (TG)-AG-AD-asialo-AGP

To assess the contribution of terminal GlcNAc to ligand recognition, AG-AD-asialo-AGP was treated with β-N-acetylglucosaminidase. Intact MS confirmed extensive GlcNAc removal (Figure S3g, Table S6). The predominant family, P′′1, corresponded to a Man_3_GlcNAc_2_ core at each of the five *N*-glycosylation sites, apart from variable residual core fucosylation. The minor P′′2 and P′′3 families were consistent with retention of one and two enzyme-resistant bisecting GlcNAc residues, respectively. Released *N*-glycan analysis confirmed *N*-glycan truncation but did not permit localization of the bisecting GlcNAc residues (Figure S3h and S4d).

#### Core GlcNAc (CG)-TG-AG-AD-asialo-AGP

To examine the contribution of the residual *N*-glycan core to ligand recognition, TG-AG-AD-asialo-AGP was treated with Endo F3. Cleavage within the chitobiose core leaves a single GlcNAc attached to the protein or, at core-fucosylated sites, a GlcNAc–Fuc remnant. Intact MS revealed a series of compositions, designated P′′′1–P′′′11, reflecting AGP molecules with the Man_3_GlcNAc_2_ core retained at between zero and five sites (Figure S3i, Table S7). The endpoint populations, corresponding to removal of the Man_3_GlcNAc_2_ core from all five sites (P′′′1) or its retention at all five sites (P′′′11), were detected at low abundance and excluded from subsequent binding and thermodynamic analyses. Site-specific glycopeptide analysis confirmed that Endo F3 cleavage efficiency varied among the five sites (Figure S3j). Cleavage was greatest at Asn85, followed by Asn38 and Asn75, and was less extensive at Asn15 and Asn54. Released *N*-glycan analysis revealed two predominant products corresponding to the Endo F3-cleaved and uncleaved core structures (Figure S4e).

### Warfarin binding to AGP F1 and S variants

#### Effects of glycosylation on affinity

To determine how *N*-glycan structure influences warfarin recognition by the F1 and S variants, nMS titrations were performed using the panel of enzymatically remodeled AGP samples. Free AGP species and their corresponding 1:1 AGP– warfarin complexes were resolved (Figure 2a), enabling the bound fraction and *K*_d_ to be determined independently for each glycan composition (Figure 2b). Comparison across the compositional series, therefore, provided a quantitative measure of how specific glycan features influence warfarin affinity.

**Figure 2.**
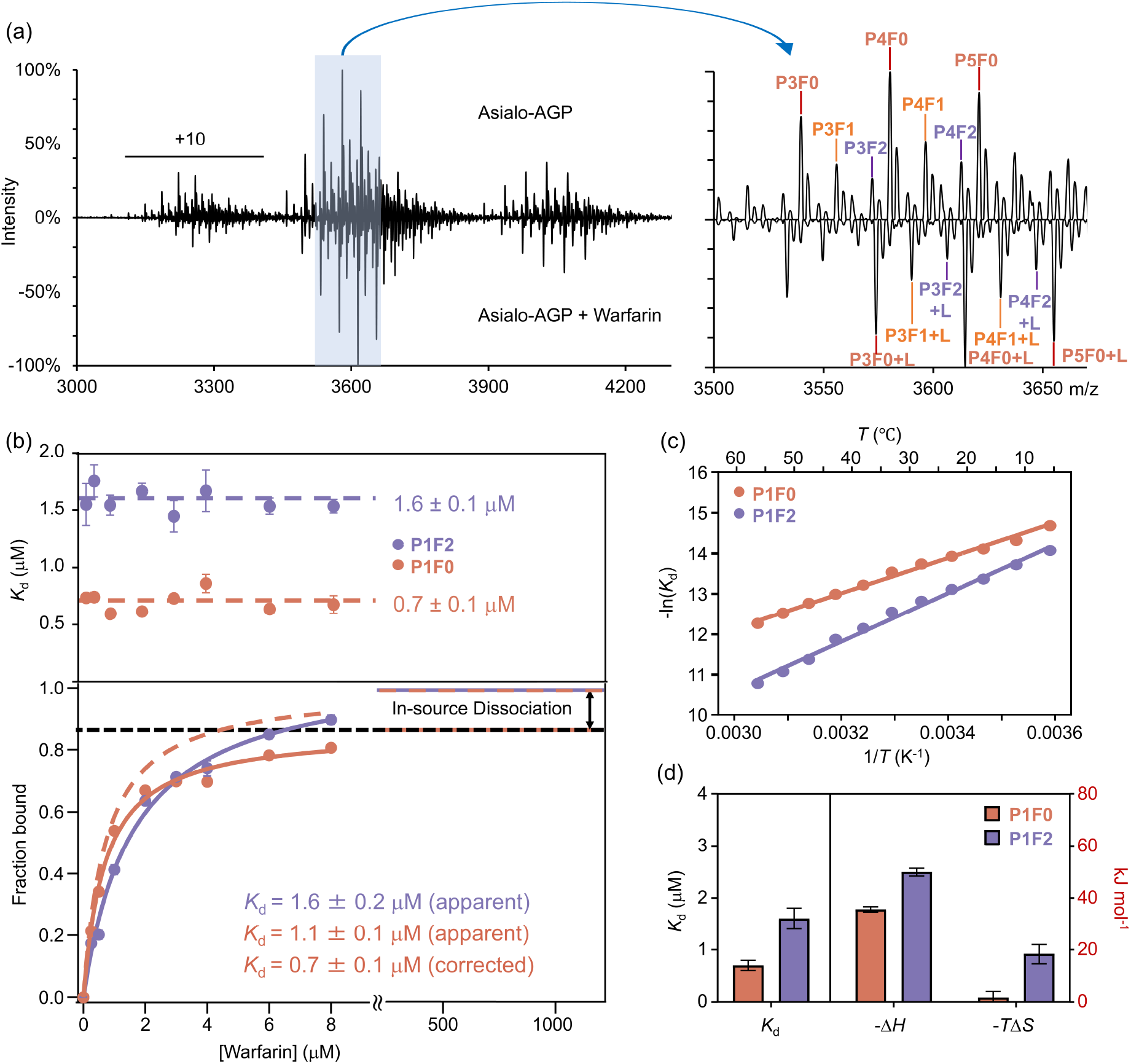
Representative nMS measurements of warfarin binding to asialo-AGP F1. (a) Native mass spectra acquired for asialo-AGP (3 μM) in the absence or presence of warfarin (6 μM) in 200 mM ammonium acetate (pH 7.6, 25 °C). The left panel shows the region containing signals for free and warfarin-bound asialo-AGP in the 8+ to 10+ charge states. The right panel shows an expanded view of the region highlighted in blue. (b) Warfarin binding to representative unfucosylated (P1F0) and difucosylated (P1F2) asialo-AGP F1 variant compositions. The lower panels show the fraction of bound AGP as a function of warfarin concentration. Solid curves represent fits to the apparent binding data, whereas red dashed curves represent fits after correction for in-source dissociation. Corrected *K*_d_ was determined using a one-site binding model. The upper panels show the corrected *K*_d_ values (eq S11) calculated at each warfarin concentration; horizontal dashed lines indicate the average *K*_d_ values. (c) van’t Hoff plots for warfarin binding to the unfucosylated and difucosylated asialo-AGP compositions, obtained by vT-nMS. (d) Comparison of the *K*_d_, −Δ*H*, and -*T*Δ*S* values; the *K*_d_ and -*T*Δ*S* values correspond to a temperature of 298.15 K.

A notable feature of the titration curves was that the apparent maximum bound fraction varied substantially among glycan compositions. Whereas some approached complete saturation, others plateaued at approximately 80%, even at warfarin concentrations expected to be saturating (Figures 2b and S6). Such sub-stoichiometric plateaus cannot be explained by differences in affinity alone and are consistent with composition-dependent dissociation of AGP–warfarin complexes during ESI and ion transfer into the mass spectrometer. If uncorrected, this in-source dissociation would systematically overestimate *K*_d_. Therefore, to obtain more reliable estimates of the solution-phase affinities, the titration data were analyzed using a binding model that explicitly accounts for in-source dissociation (eq S8).^45^ The model assumes that, for each glycan composition, a constant fraction of the complex dissociates during analysis, independent of ligand concentration. The corrected *K*_d_ values calculated at individual ligand concentrations were essentially concentration-independent and agreed closely with those obtained by global non-linear regression (Figure 2b), supporting the internal consistency of the correction.

#### Influence of fucosylation and glycan branching

Corrected *K*_d_ values (Figure 3) for individual asialo-AGP glycan compositions spanned approximately sevenfold for F1 (0.3–2.2 μM, Table S8) and fourfold for S (0.2–0.8 μM; Table S10). Among the lower-mass branching families (P1–P5) of the F1 variant, difucosylated species bound warfarin more weakly than their unfucosylated counterparts (Figure S6). This difference progressively diminished across P6–P9. Monofucosylated species generally exhibited affinities similar to those of their unfucosylated counterparts, although the monofucosylated P9 species bound warfarin more strongly. Trifucosylated compositions exhibited variable effects, with affinities that were comparable to, weaker than, or stronger than those of the corresponding unfucosylated species. In contrast, difucosylation of the S variant generally enhanced warfarin binding among the lower-mass families (P2–P4), although this effect diminished or became variable across P5–P9 (Figure 3b). Mono- and trifucosylated S variant compositions showed comparatively modest or variable differences from their unfucosylated counterparts.

**Figure 3.**
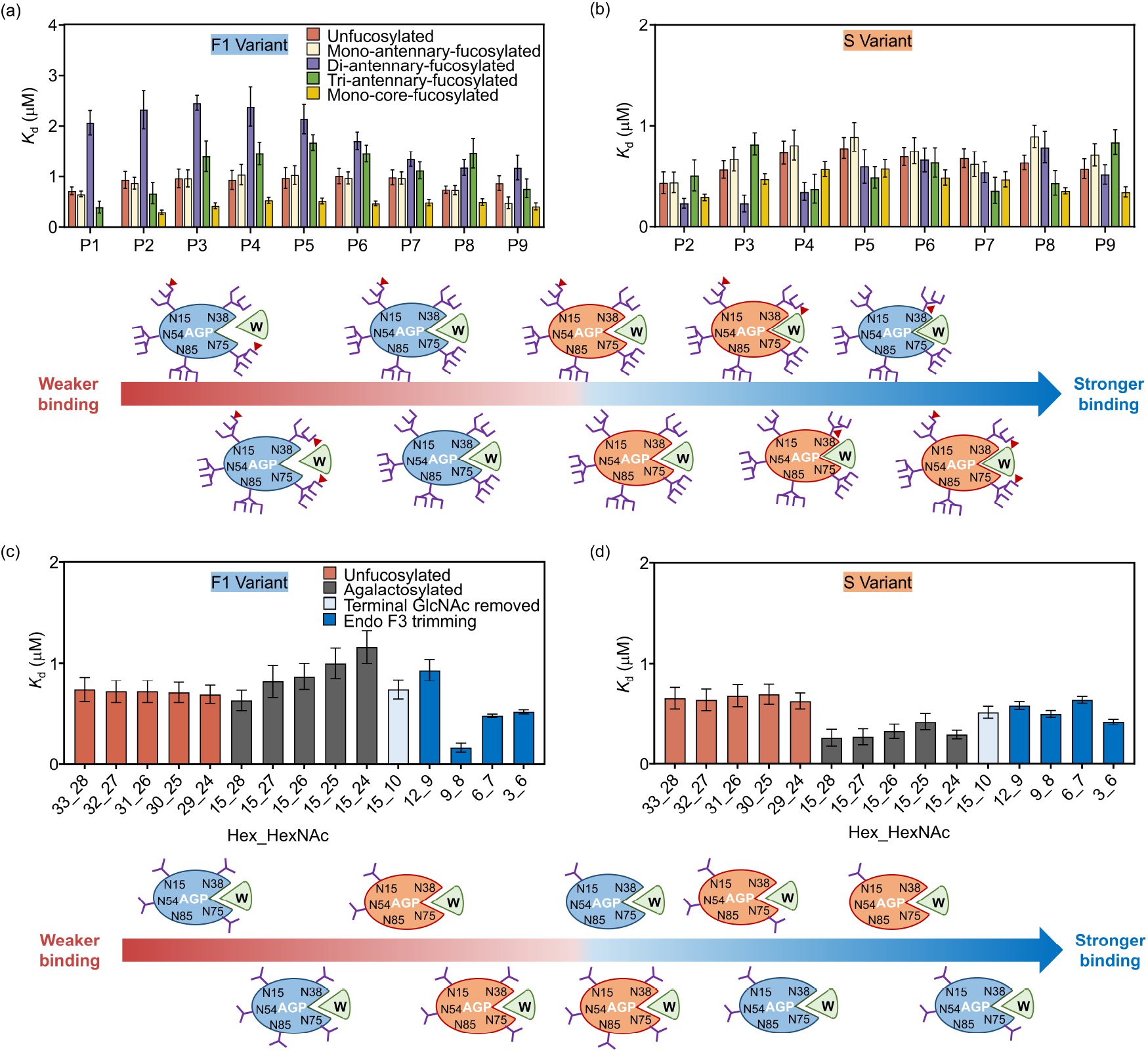
Effects of fucosylation and glycan trimming on warfarin binding to AGP. Comparison of *K*_d_ values across fucosylation states for asialo-AGP and AD-asialo-AGP compositions of the (a) F1 and (b) S variants. Effects of sequential glycan trimming on *K*_d_ for the (c) F1 and (d) S variants. Unfucosylated compositions are shown for AD-asialo-AGP (red, P1-P5, Hex_HexNAc: 29_24 to 33_28), AG-AD-asialo-AGP (grey, P’1-P’5, Hex_HexNAc: 15_24 to 15_28), TG-AG-AD-asialo-AGP (light blue, P’’1, Hex_HexNAc: 15_10), CG-TG-AG-AD-asialo-AGP (blue, P’’’2-P’’’9, Hex_HexNAc: 3_6 to 12_9). Glycan compositions are specified as Hex_HexNAc. Schematics summarize qualitative affinity trends across selected glycosylation states; depicted glycan locations are illustrative and do not represent uniquely resolved site-specific assignments. All *K*_d_ values correspond to 298.15 K.

#### Opposing effects of core and antennary fucosylation

Comparison of F1 and S compositions with equivalent branching revealed a consistent trend toward stronger warfarin binding by the residual core-fucosylated species in AD-asialo-AGP than by their unfucosylated counterparts (Figure S7, Tables S9 and S11). This trend was observed across the composition families examined, suggesting that the favorable effect of core fucosylation is largely independent of glycan size. In contrast, before antennary defucosylation, difucosylated F1 compositions generally bound more weakly than their unfucosylated counterparts, whereas difucosylated S compositions generally exhibited stronger binding among the lower-mass families. Together, these results suggest that antennary Fuc is primarily responsible for the reduced affinity associated with total fucosylation in the F1 variant, whereas core α1–6 Fuc tends to enhance warfarin binding by both variants.

To estimate the respective effects of the two fucosylation motifs, binding data obtained before and after α1–3,4 fucosidase treatment were analyzed together. The model incorporated the measured affinities and distributions of total fucosylation states determined by intact MS, with residual fucosylation assigned as core linked Fuc (Figure 3a and 3b). For the F1 variant, core fucosylation increased warfarin affinity by approximately two-fold, whereas antennary fucosylation decreased affinity by as much as two-fold. Because antennary Fuc was more abundant, its unfavorable contribution dominated the net effect, explaining why difucosylated compositions generally bound more weakly despite the favorable contribution of core fucosylation. Core fucosylation similarly favored binding by the S variant; however, antennary fucosylation did not exert the consistently unfavorable effect observed for F1. Thus, although core fucosylation enhanced warfarin binding by both variants, the influence of antennary fucosylation was variant dependent.

These opposing and variant-dependent effects may explain the difference from the findings of Robinson and co-workers, who reported that both increasing *N*-glycan branching and monofucosylation attenuate warfarin binding.^29^ Because the relative proportions of core- and antennary-fucosylated glycans can vary among AGP preparations, the apparent effect of total fucosylation will depend on the distribution of the two motifs. Core and antennary fucosylation should, therefore, be considered independently when evaluating glycosylation-dependent ligand recognition.

#### Apparent suppression of warfarin binding by bisecting GlcNAc

No warfarin-bound complexes were detected for compositions assigned as containing bisecting GlcNAc in either asialo-AGP or AD-asialo-AGP, despite clear complex formation by other compositions in the same spectra (Figure S10). This consistent absence suggests that bisecting GlcNAc may substantially weaken warfarin binding. However, preferential in-source dissociation of these complexes cannot be excluded, as the measurements cannot distinguish reduced solution-phase affinity from reduced gas-phase complex stability. If the effect originates in solution, bisecting GlcNAc may alter the conformational ensemble of the attached *N*-glycan and, through glycan– protein interactions, influence the structure, hydration, or accessibility of the warfarin-binding cavity.

#### Effects of sequential outer-arm trimming

Sequential removal of terminal Gal (AG-AD-asialo-AGP) and GlcNAc (TG-AG-AD-asialo-AGP) further simplified the *N*-glycan structures (Figures S11 and S13-S14) but produced little overall change in the corrected *K*_d_ values for the F1 variant (Figure 3c). Across both remodeling steps, core-monofucosylated compositions consistently tended to bind warfarin more strongly than their unfucosylated counterparts, indicating that the favorable contribution of core fucosylation persisted after truncation of the glycan antennae. The magnitude of this difference nevertheless decreased with progressive trimming, suggesting that the outer branches modulate, but are not solely responsible for, the effect of core fucosylation. The S variant exhibited a distinct response to outer-arm trimming: removal of terminal Gal generally decreased *K*_d_, whereas subsequent removal of terminal GlcNAc increased *K*_d_, partially reversing the affinity enhancement associated with Gal removal (Figure 3d). Core-monofucosylated S compositions continued to bind more strongly than their unfucosylated counterparts, although this difference diminished with progressive trimming.

#### Dependence on the number of retained *N*-glycan cores

Analysis of CG-TG-AG-AD-asialo-AGP allowed us to examine how the number of sites retaining an intact Man₃GlcNAc₂ core influences warfarin binding (Figure S3i). Endo F3 treatment cleaved the cores at varying numbers of sites, leaving a single protein-linked GlcNAc at each cleaved site, with core Fuc retained where present. Species retaining either no intact cores or all five cores were insufficiently abundant in the Endo F3-treated CG-TG-AG-AD-asialo-AGP for reliable quantification. Binding analysis of the Endo F3-treated sample was therefore restricted to species retaining one to four cores, with TG-AG-AD-asialo-AGP before Endo F3 treatment serving as the five-core reference.

Warfarin affinity varied non-monotonically with the number of retained cores (Figure 3c). F1 variant species retaining one or two cores exhibited similar *K*_d_ values of 0.53 ± 0.03 and 0.48 ± 0.02 μM, respectively. Affinity was greatest for the three-core population (0.16 ± 0.04 μM) but decreased markedly for the four-core population (0.91 ± 0.08 μM), approaching that of TG-AG-AD-asialo-AGP with all five cores retained (0.71 ± 0.10 μM). Thus, among the populations quantified, affinity was greatest at an intermediate degree of *N*-glycan truncation.

The site-specific susceptibility to Endo F3 provides a possible explanation for this non-monotonic behavior. Glycopeptide analysis suggests that the four-core population is enriched in species cleaved at Asn85, whereas the three-core population likely includes additional cleavage at Asn38. Notably, Asn38 is positioned near the entrance to the ligand-binding cavity, where its *N*-glycan may influence local structural organization and ligand accessibility. Cleavage of this glycan could therefore alter the environment of the cavity and contribute to the enhanced warfarin affinity observed for the three-core population (0.16 ± 0.04 μM). However, intact MS establishes the number of retained cores without distinguishing species with different site-specific patterns of cleavage. Consequently, the enhanced affinity cannot be attributed specifically to cleavage at Asn38, although the results suggest that the locations of the retained *N*-glycan cores may influence warfarin binding.

### Glycan-dependent thermodynamics of warfarin binding to AGP

Temperature-dependent measurements of the enzymatically remodeled AGP samples enabled the thermodynamic parameters of warfarin binding to be determined for individual glycan compositions. The resulting van’t Hoff plots were linear over the temperature range examined (Figures 2c and S8), indicating that changes in binding heat capacity were not resolvable and supporting use of the linear van’t Hoff relationship. The slopes and intercepts yielded Δ*H* and Δ*S*, respectively, after accounting for the corresponding gas-constant terms, providing a composition-resolved measure of how specific glycan features alter the enthalpic and entropic contributions to AGP–warfarin binding.

The vT-nMS-derived parameters were compared with measurements of warfarin binding to asialo-AGP by ITC. Population-weighted mean dissociation constants, enthalpies, and entropies were calculated from the composition-resolved nMS results using their relative abundances. These means agreed closely with the apparent parameters obtained from single-site ITC analysis (Figure S5), supporting consistency between the methods.

#### Core versus antennary fucosylation

Fucosylation produced composition- and variant-dependent changes in the thermodynamics of warfarin binding (Figure 4). For the F1 variant, difucosylated species in antennary-fucose-rich asialo-AGP generally exhibited more favorable binding enthalpies than their unfucosylated counterparts, particularly across P1–P5, but this stabilization was offset by a greater entropic penalty (Figures 4a and S8). These differences diminished across P6–P9. Monofucosylated species generally exhibited the opposite pattern, less favorable enthalpy accompanied by a more favorable entropic contribution, whereas trifucosylated species resembled their unfucosylated counterparts. Deconvolution of the core- and antennary-fucosylated populations associated core fucosylation with a consistent shift toward less favorable enthalpy and more favorable entropy. For the S variant, both fucosylation motifs produced this general thermodynamic signature, with the largest compensating changes observed for difucosylated species (Figure 4b).

**Figure 4.**
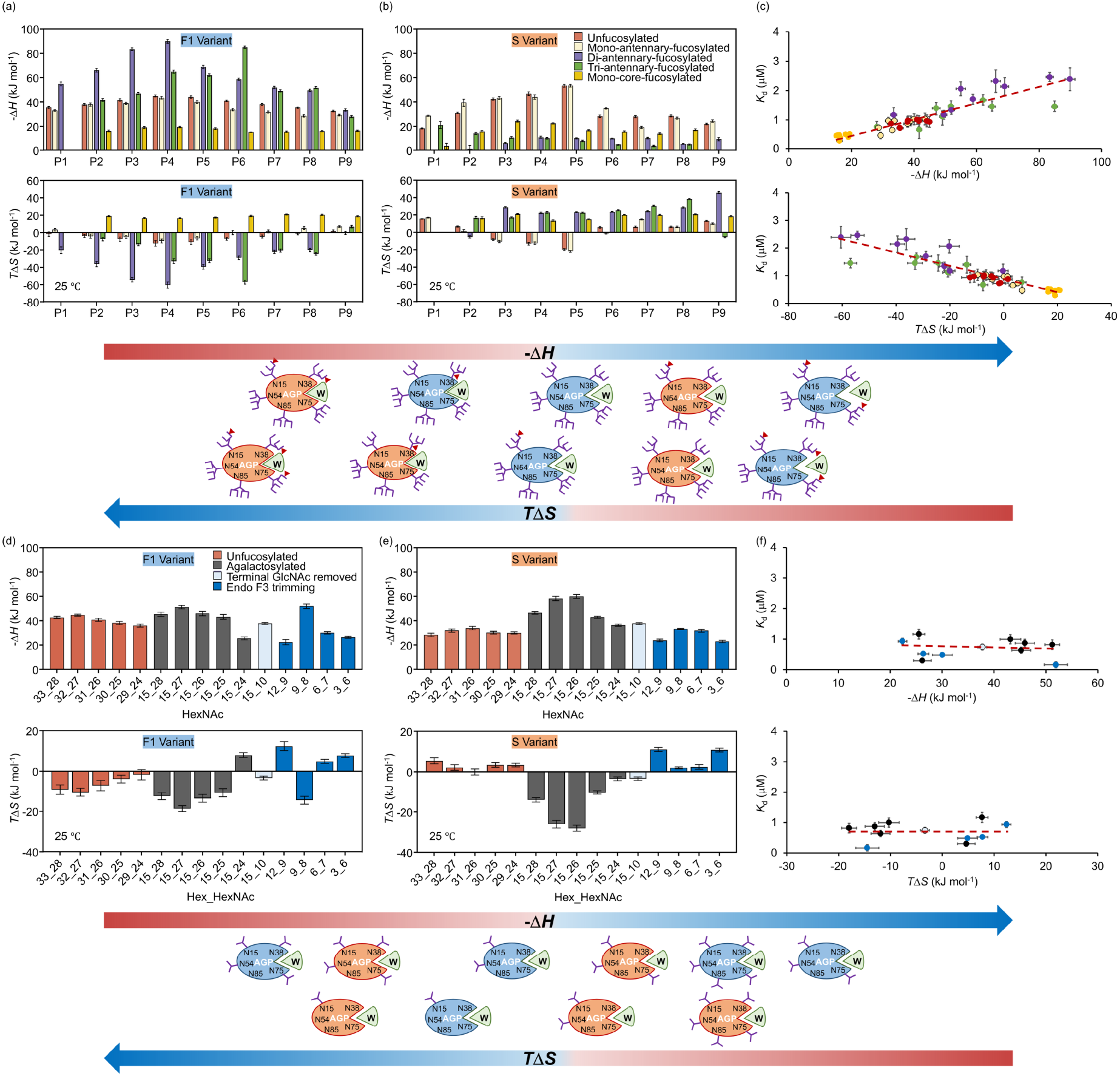
Effects of fucosylation and glycan trimming on the thermodynamics of warfarin binding to AGP. Enthalpic (−Δ*H*) and entropic (*T*Δ*S*) contributions to binding by asialo-AGP and AD-asialo-AGP compositions of the (a) F1 and (b) S variants, compared across fucosylation states. (c) Relationships between *K*_d_ and −Δ*H* (upper panel; R² = 0.82) or *T*Δ*S* (lower panel; R² = 0.80) for the F1 fucosylation series. Thermodynamic contributions following sequential glycan trimming for the (d) F1 and (e) S variants. Glycan compositions are specified as Hex_HexNAc. (f) Corresponding relationships between *K*_d_ and −Δ*H* (upper panel; R² = 0.02) or *T*Δ*S* (lower panel; R² = 0.01) for the F1 trimming series. Dashed red lines indicate linear fits. Schematics summarize the qualitative trends in the enthalpic and entropic contributions across selected glycosylation states. *T*Δ*S* values are reported at 298.15 K.

Thus, core and antennary fucosylation are not thermodynamically equivalent (Figure S9). Core α1–6 Fuc, which lies near the protein surface, may restrict or redistribute the conformational ensemble of the attached *N*-glycan, thereby reducing the conformational reorganization required upon binding. When located near the binding cavity, core Fuc may also influence ligand access. The MD simulations described below support this interpretation. In contrast, the more favorable enthalpy and greater entropic penalty associated with antennary fucosylation in the F1 variant suggest increased conformational or hydration reorganization upon binding. The reversal of this thermodynamic signature in the S variant indicates that the effect of antennary Fuc depends strongly on its structural context and coupling to the protein.

Across the fucosylated AGP compositions, increasingly favorable binding enthalpy was accompanied by an increasingly unfavorable entropic contribution, demonstrating pronounced enthalpy–entropy compensation. Consequently, fucosylation can substantially redistribute the enthalpic and entropic contributions to AGP–warfarin recognition while producing comparatively modest or even counterintuitive changes in net affinity.

#### Thermodynamic effects of sequential glycan trimming

Removal of terminal Gal and GlcNAc produced variant-dependent changes in the enthalpic and entropic contributions to warfarin binding, even where changes in *K*_d_ were comparatively small (Figures 4d, 4e and S12–S14). Thus, the limited changes in affinity observed for some compositions concealed changes in the underlying binding thermodynamics. Nevertheless, the thermodynamic signature associated with core fucosylation persisted after outer-arm trimming: core-monofucosylated compositions of both AG-AD-asialo-AGP and TG-AG-AD-asialo-AGP exhibited less favorable enthalpic contributions but more favorable entropic contributions than their unfucosylated counterparts.

Endo F3 treatment produced a distinct dependence on the number of sites retaining an uncleaved Man_3_GlcNAc_2_ core (Figure 4d). For the F1 variant, the four-core population exhibited a thermodynamic profile similar to that of AGP retaining cores at all five sites. By contrast, the three-core population showed substantially more favorable binding enthalpy, partially offset by a larger entropic penalty. Further truncation to two retained cores reduced the enthalpic stabilization and produced a more favorable entropic contribution. Glycopeptide analysis indicated that Endo F3 cleavage was most extensive at Asn85, followed by Asn38 and Asn75. The four-, three-, and two-core populations may, therefore, be progressively enriched in species cleaved at these sites in that order. However, because each population contains unresolved site-occupancy isomers, the thermodynamic effects cannot be assigned to cleavage at a particular site.

For the S variant, Endo F3-mediated glycan trimming produced thermodynamic trends similar to those observed for F1, with a notable divergence for the three-core population (Figure 4e). Compared with the corresponding F1 population, the S three-core population exhibited less favorable binding enthalpy and a favorable, rather than unfavorable, entropic contribution. The Q20R substitution in the S variant could alter structural coupling between the region surrounding residue 20 and the Asn38 glycosylation site, causing glycan removal to reorganize protein–glycan interactions, conformational dynamics or hydration differently from F1. Although this possibility requires validation, the distinct behavior of the three-core population points to potential coupling between protein sequence and site-specific glycosylation.

Across the fucosylation series, stronger warfarin binding was generally associated with a more favorable entropic contribution despite less favorable enthalpy. The F1 glycan-trimming series showed a different pattern, with little linear correlation between *K*_d_ and either −Δ*H* or *T*Δ*S* (Figure 4f). The distinct thermodynamic profiles of the F1 and S three-core populations further illustrate that the balance of enthalpic and entropic contributions depends on both the extent of trimming and the AGP variant.

### MD simulations and ReGlyco Ensemble analysis

To investigate the structural basis of the experimentally observed effects of fucosylation on warfarin binding, we focused on Asn38 and Asn75, the glycosylation sites closest to the entrance of the ligand-binding funnel, where attached glycans could modulate ligand accessibility. All experimentally determined structures of human AGP available in the PDB are from proteins expressed in *E. coli* and thus lacking glycosylation. Across these structures, encompassing the A, S, and F1 variants, Asn38 lies near the middle of an α-helical segment connecting two β-strands at the funnel entrance (Figure 5a). Given the reported potential of *N*-glycosylation to disrupt α-helical structure,^46,47^ we hypothesized that glycosylation at Asn38 could alter the extent or stability of this helix in fully glycosylated human AGP. We used two complementary strategies to account for this possibility in our structural analysis.

**Figure 5.**
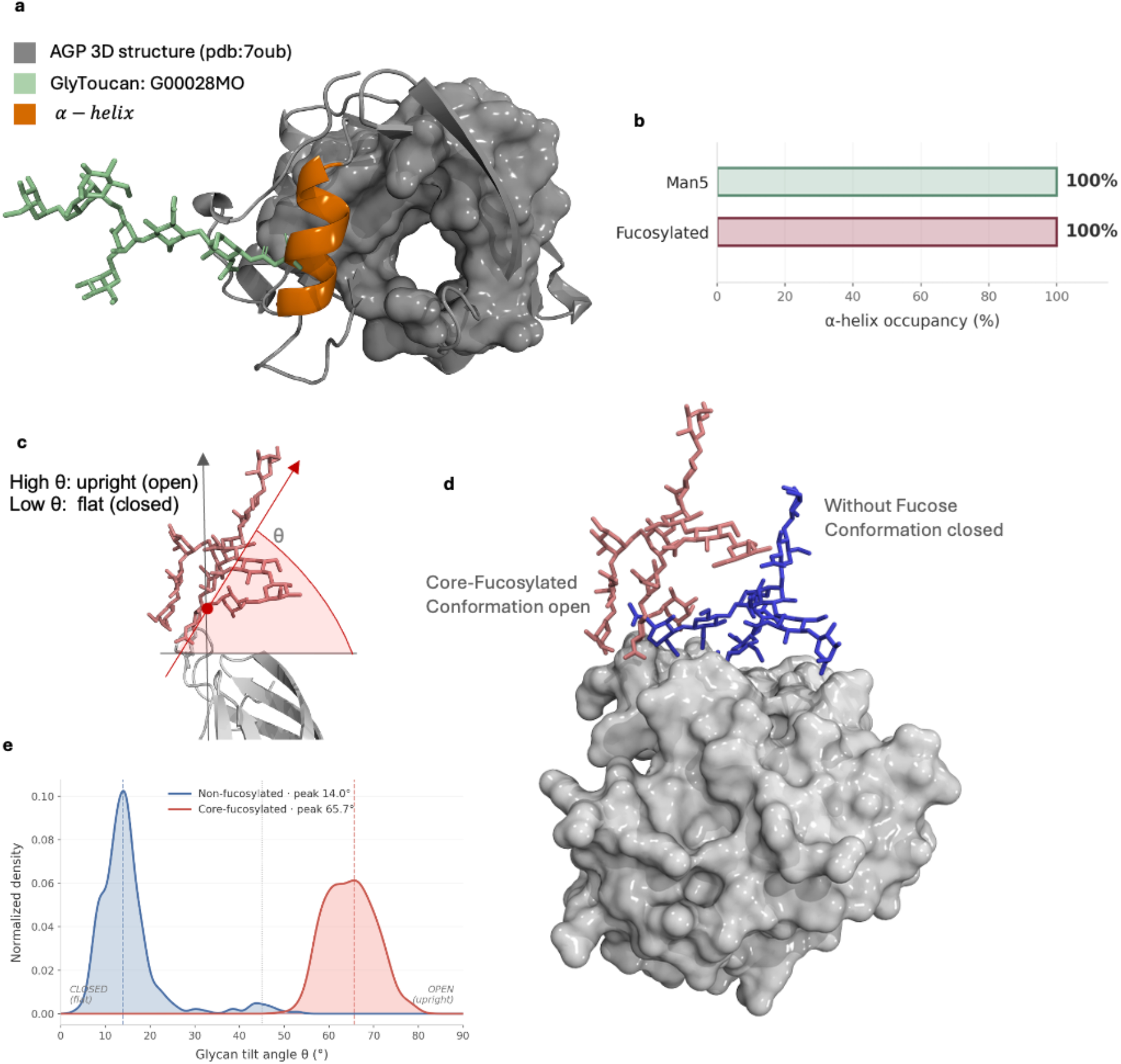
(a) AGP 3D structure (PDB 7OUB, grey surface) with the Asn38 α-helix highlighted (orange) and the Man5 *N*-glycan (GlyTouCan G00028MO, green sticks) modeled at this site. (b) α-helix occupancy measured with the *cpptraj* module of Amber14 based on DSSP values^49^ over the 1 µs GaMD trajectories of the fully glycosylated PDB 7OUB (1.82 Å), for the Man5 (GlyTouCan ID G00028MO) and core-fucosylated complex glycan (GlyTouCan ID G87921DA) compositions. (c) Definition of the glycan tilt angle (θ) used to quantify the orientation of the Asn38 glycan relative to the binding-pocket entrance plane in the Asn38 loop model of F1. (d) Representative structures of the non-fucosylated (blue) and core-fucosylated (salmon) Asn38 glycan relative to the AGP F1 variant surface (grey), illustrating the closed and open conformations, respectively. (e) Probability density distributions of the glycan tilt angle θ from enhanced-sampling MD simulations of the Asn38 loop model of F1, comparing non-fucosylated (blue, peak at 14.0°) and core-fucosylated (red, peak at 65.7°) Asn38 glycan compositions.

First, we introduced an *N*-glycan at Asn38 into an experimentally determined AGP structure and ran MD with an enhanced sampling scheme to assess the stability of the α-helical segment following glycosylation. We chose PDB 7OUB as a protein model, corresponding to variant A, with the highest-resolution experimental structure available. Using GlycoShape ReGlyco,^37^ we glycosylated Asn38 with either Man5 (GlyTouCan ID G00028MO) or a core-fucosylated complex *N*-glycan (GlyTouCan ID G87921DA). Each system was sampled for 1 μs with GaMD.^41^ The α-helical segment containing Asn38 remained stable throughout both trajectories (Figure 5b). With Asn38 retained in this α-helical conformation, the attached *N*-glycan projected away from the entrance of the ligand-binding funnel for both glycan structures examined.

As a second strategy, we generated a model of the AGP S variant using RFdiffusion3,^48^ constraining residues 37–42 to adopt a loop conformation in place of the corresponding α-helical segment. We then performed GaMD simulations to assess whether this region regained partial helicity and to characterize the associated conformational behaviour of the Asn38 *N*-glycan. The F1 variant, which differs from the S variant by a single substitution at residue 20, was built and simulated independently using the same approach and yielded a structure indistinguishable from that of the S variant.

In the F1 variant, GaMD did not recover any significant degree of helicity in the region but it produced a wide variety of orientations of the Asn38 glycan relative to the funnel entrance. These orientations were quantified using a glycan tilt angle (θ) relative to the funnel-entrance plane defined by the Cα atoms of residues 67–69, 71–73, and 91–92, with larger angles corresponding to more upright glycan orientations (Figure 5c). Man5 adopted an orientation that partially occluded the funnel entrance, with the distribution maximum value at θ = 14.0°. Meanwhile, the core-fucosylated glycan adopted a more upright orientation, with a distribution maximum value at θ = 65.7°, positioning the glycan away from the funnel entrance (Figure 5d and 5e).

Although these GaMD simulations do not establish the conformation of the Asn38 region in glycosylated human AGP, the backbone outside the Asn38 loop remained stable in the loop-model simulation (Figure S15). The results support changes in glycan orientation near the funnel entrance as a possible contributor to glycosylation-dependent warfarin recognition, without excluding contributions from local structural rearrangements, protein dynamics, or hydration.

To investigate this aspect across a range of glycoforms representative of the experimentally observed glycan microheterogeneity of AGP, we used GlycoShape Re-Glyco Ensemble.^37^ We first compared an unfucosylated reference glycoform (G0) with glycoforms carrying core fucose at Asn38 (G1, Figure 6c), Asn75 (G2, Figure 6e), or both sites (G3). For this analysis we focused on Asn75, the other *N*-glycosylation site at the funnel entrance, which, unlike Asn38, lies within a well-defined structured region of the protein (Figure 6a).

**Figure 6.**
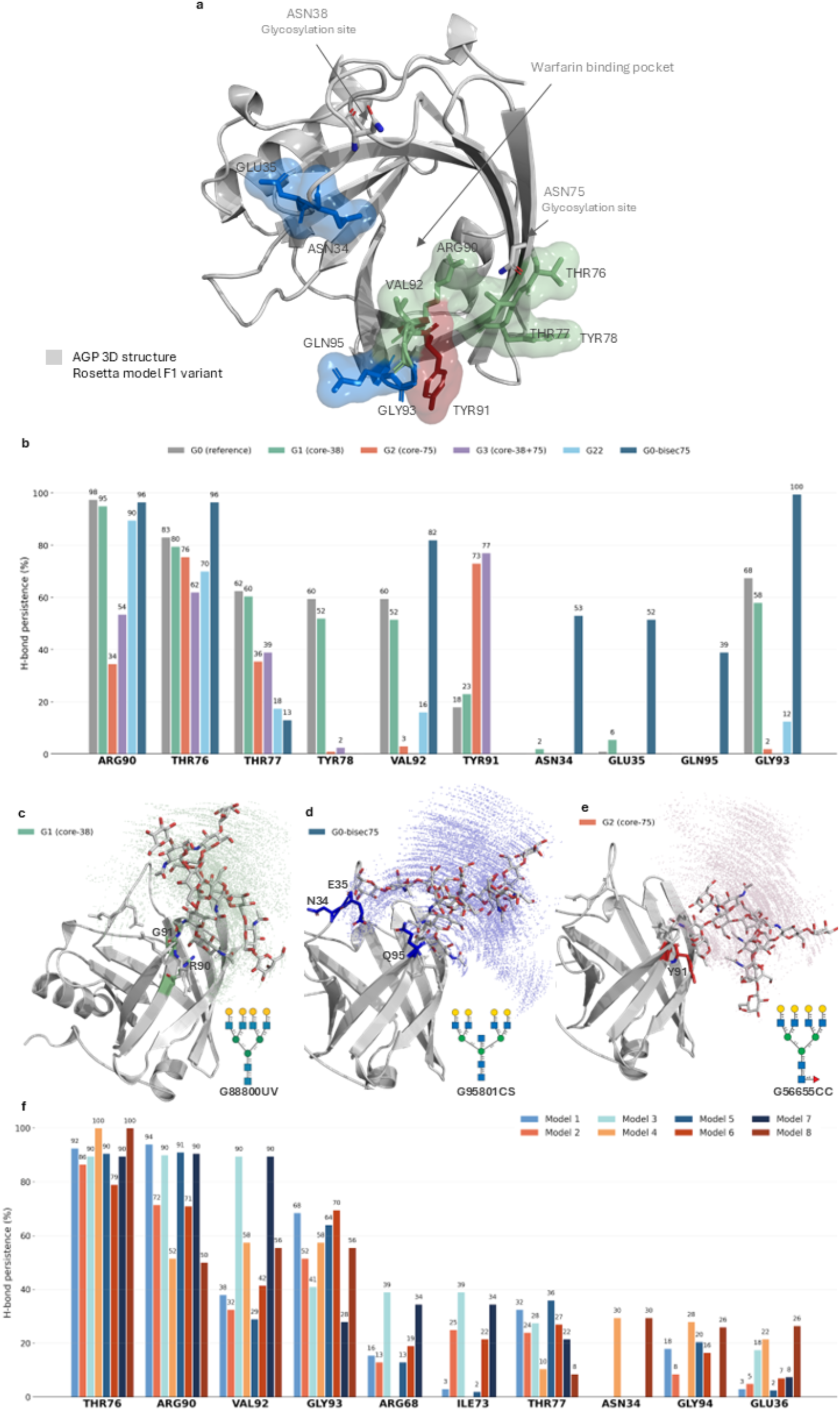
(a) RFdiffusion3 model of the AGP F1 variant (grey cartoon) showing the Asn38 and Asn75 *N*-glycosylation sites and the entrance to the warfarin-binding pocket. Residues included in the glycan–protein interaction analysis are shown as sticks. (b) Hydrogen-bond persistence between AGP residues and the Asn75-linked *N*-glycan in reconstructed ensembles of fully glycosylated S-variant glycoforms. G0 represents the reference glycoform in which none of the five *N*-glycans carries core fucose; G1 and G2 contain core α1–6 fucose at Asn38 and Asn75, respectively, while G3 contains core α1–6 fucose at both sites. Additional glycoforms include G22 and G0-bisec75 [definitions given in Table S1]. Hydrogen-bond persistence was calculated over the conformational ensemble of the Asn75-linked glycan. (c–e) Representative conformations of the Asn75-linked *N*-glycan and the corresponding conformational ensembles for G1 (c), G0-bisec75 (d), and G2 (e). The protein is shown as a grey cartoon. Selected glycan conformations are shown as sticks, while the coloured dots represent the positions of the glycan heavy atoms across 200 conformations generated using Re-Glyco, illustrating their spatial distribution around the pocket entrance. The selected conformations belong to GlycoShape Cluster 0 for G1 (44.18% of the glycan conformational ensemble) and G0-bisec75 (63.49%), and Cluster 1 for G2 (31.16%). The corresponding glycan structures are shown using Symbol Nomenclature for Glycans (SNFG). (f) Hydrogen-bond persistence between AGP residues and the Asn75-linked glycan across eight models containing antennary fucosylation. Models 1–8 differ in glycan branching at Asn38 and Asn75 and in the antenna carrying fucose. The complete glycan compositions and structural definitions of Models 1–8 are reported in Table S1.

Analysis of the Asn75 glycan ensembles revealed a clear effect of core fucosylation (Figure 6b). In G0, the Asn75 glycan interacted predominantly with residues surrounding the glycosylation site, including Arg90 and Thr76. In the S-variant ensembles, core fucosylation at Asn75 increased the percentage of conformations with an Asn75 glycan–Tyr91 hydrogen bond from 18% in G0 to 73% in G2 and 77% in G3 (Figure 6b). The F1 ensembles showed the same qualitative trend (Figure S17). This increase was not observed when core fucose was present only at Asn38 (G1, 23%), indicating that it arises primarily from core fucosylation of the glycan at Asn75. The increased hydrogen-bond interaction with Tyr91, which is positioned laterally to the funnel entrance, was associated with glycan conformations directed toward the protein surface and away from the central opening (Figures 6a and 6e).

Antennary fucosylation produced a substantially more heterogeneous pattern (Figure 6f). To investigate this variability, we generated eight models differing in glycan branching at Asn38 and Asn75 and in the antenna carrying fucose (Models 1–8; Table S1). No common glycan–protein interaction pattern emerged across these models (Figures 6f and S16). Instead, the effect on the binding-funnel environment depended on the specific glycan context, including the degree of branching and the antenna carrying fucose. Thus, unlike core fucosylation, antennary fucosylation could not be associated with a single structural mechanism affecting funnel accessibility, consistent with its variable effects on warfarin affinity observed experimentally.

The results also indicated that the presence of a bisected *N*-glycan at Asn75 substantially reorganized the local glycan–protein hydrogen-bonding network (Figures 6b and S17). Compared with G0, G0-bisec75 exhibited persistent hydrogen-bond interactions involving Asn34, Glu35, and Gln95, whereas hydrogen-bond persistence with Tyr91, Thr77, and Tyr78 was markedly reduced. These changes suggest that bisection redistributes glycan–protein interactions toward a distinct set of residues. This redistribution may favour glycan conformations that obstruct the funnel entrance and restrict access to the ligand-binding cavity, providing a possible structural explanation for the absence of detectable warfarin complexes.

## CONCLUSIONS

This study demonstrates that specific *N*-glycan features influence both the affinity and thermodynamics of warfarin binding to human AGP, with effects that depend on glycan composition and protein variant. By integrating sequential enzymatic remodeling with composition-resolved nMS, vT-nMS, complementary glycosylation analyses, and molecular modelling, we distinguished functional differences within the heterogeneous F1 and S populations. Thermodynamic analysis revealed that glycan modifications could substantially alter the enthalpic and entropic contributions to warfarin binding while producing comparatively small changes in *K*_d_. Core and antennary fucosylation exerted particularly distinct effects. Core fucosylation favored warfarin binding by both variants and shifted the thermodynamics toward less favorable enthalpy and more favorable entropy. The effects of antennary fucosylation on affinity depended on glycan composition and protein variant, generally weakening binding by F1 while producing more variable effects in S. Despite these differences, stronger binding across the fucosylation series was generally associated with more favorable entropic contributions. Thus, the two variants shared a common relationship between affinity and the underlying thermodynamic contributions, although their responses to antennary fucosylation differed.

Sequential glycan trimming further showed that ligand recognition does not depend simply on overall glycan size. Removal of terminal Gal and GlcNAc produced variant-dependent effects on affinity and altered the balance of enthalpic and entropic contributions, even where changes in *K*_d_ were comparatively small. Endo F3-mediated truncation produced a pronounced non-monotonic response in F1, with the three-core population exhibiting the strongest binding and greatest enthalpic stabilization within the truncation series. The corresponding S population exhibited a more favorable entropic contribution than F1 without a comparable affinity enhancement. These differences suggest that protein sequence influences the response to glycan truncation. Site-specific glycopeptide analysis provided possible explanations involving differential cleavage among glycosylation sites, but the unresolved distributions of retained cores prevent assignment of the binding effects to individual sites. Notably, no warfarin-bound complexes were detected for compositions assigned as containing bisecting GlcNAc. This observation suggests substantial suppression of binding, although preferential in-source dissociation remains an alternative explanation.

MD simulations and reconstructed glycan ensembles suggested site-specific mechanisms by which glycan modifications influence AGP funnel accessibility. In the Asn38 loop model, the core-fucosylated complex glycan adopted a more upright orientation than the unfucosylated glycan, reducing funnel occlusion, whereas core fucosylation at Asn75 redirected the glycan toward the protein surface. Antennary fucosylation showed no single structural mechanism, consistent with its variable effects on affinity. Bisecting GlcNAc substantially reorganized the Asn75 glycan–protein hydrogen-bond network, providing a potential structural explanation for the absence of detectable warfarin complexes.

Together, these findings highlight how glycan modifications may influence ligand recognition through changes in glycan orientation and local interactions around a binding pocket, without requiring direct glycan–ligand contacts. For AGP, this provides a framework for understanding how glycosylation changes may alter drug binding and why their effects depend on both glycan structure and protein variant. More broadly, the integration of composition-resolved binding measurements, thermodynamic analysis, and molecular simulations offers an approach to uncovering functional differences otherwise obscured within heterogeneous glycoprotein populations.

## SUPPORTING INFORMATION

Supporting Information accompanies this manuscript.

Experimental and computational methods; annotation of AGP glycan compositions; released *N*-glycan chromatograms and site-specific glycosylation profiles; comparison of warfarin-binding affinities and thermodynamic parameters determined by vT-nMS and ITC; warfarin-binding titration curves and van’t Hoff plots.

## ACKNOWLEDGEMENTS

The authors acknowledge funding from the Natural Sciences and Engineering Research Council of Canada (JSK), the Canada Foundation for Innovation (JSK), the Alberta Innovation and Advanced Education Research Capacity Program (JSK), and Canada Excellence Research Chairs Program (LKM).

