## Supplemental Data for "Sugar Matters: How Protein Glycosylation Shapes Ligand Binding"

* Corresponding Author:

https://orcid.org/0000-0002-3389-7112

### Table of contents

**Materials and methods……………………………………………………………………....S5**

Separation of AGP variants ………………………………..……..…………..……......S5

Preparation of enzymatically remodeled AGP samples. …..……………………...……......S5

Released *N*-glycan analysis by HILIC-UHPLC..……………………..……….……................S5

Glycopeptide analysis by nanoLC-MS/MS. ……………………..……..………………......S6

Isothermal titration calorimetry ……..………………………………………...……......S8

Native mass spectrometry ………………………...………………………………...……......S8

Temperature-controlled nanoESI device ………………………………………....……......S9

Deconvolution of ESI mass spectra …..…………………………………………......…......S9

Determination of composition-resolved binding affinities and correction for in-source dissociation……………………………………………………………………………………S9

Computational methods …..………………………………………………………………S14

**Tables**…………………………………………………………………………..............S18

**Table S1**. AGP glycoform nomenclature ………………………………...…………………S18

**Table S2**. Annotation of AGP glycan compositions ……………………...…………………S20

**Table S3**. Annotation of asialo-AGP glycan compositions.…………………...…...…………S25

**Table S4**. Annotation of AD-asialo-AGP glycan compositions.…………………………......S30

**Table S5**. Annotation of AG-AD-asialo-AGP glycan compositions.……………..….………S34

**Table S6**. Annotation of TG-AG-AD-asialo-AGP glycan compositions...………………..……S37

**Table S7**. Annotation of CG-TG-AG-AD-asialo-AGP glycan compositions …….………...…S38

**Table S8**. Affinities and thermodynamic parameters for asialo-AGP F1 variant compositions binding to warfarin…….………………………………………………………………..….S40

**Table S9**. Affinities and thermodynamic parameters for AD-asialo-AGP F1 variant compositions binding to warfarin……..………………………………………………………………….S42

**Table S10**. Affinities and thermodynamic parameters for asialo-AGP S variant compositions binding to warfarin…….……………………………...……………………………..….S43

**Table S11**. Affinities and thermodynamic parameters for AD-asialo-AGP S variant compositions binding to warfarin…….……………………………...……………………………..….S45

**Figures……………………..……………………………………………………..……..……...S46**

**Figure S1.** Representative ESI mass spectra acquired in the positive ion mode for ammonium acetate solutions of AGP………………..…………..………...………...………………..S46

**Figure S2.** Chromatogram of ProA labeled *N*-glycans derived from AGP and heatmap of glycopeptide ……………………………………………………………………………S47

**Figure S3.** Representative zero-charge deconvoluted native mass spectra and corresponding site-specific glycopeptide heatmaps for enzymatically remodeled AGP samples ……..……………..…S48

**Figure S4.** Chromatograms of ProA labeled *N*-glycans derived from asialo-AGP, AD-asialo-AGP, AG-AD-asialo-AGP, TG-AG-AD-asialo-AGP and CG-TG-AG-AD-asialo-AGP ……..……...S49

**Figure S5.** Comparison of the binding affinity and thermodynamic parameters between vT-nMS assay and ITC for AGP and asialo-AGP. ……………………………………………………S50

**Figure S6.** Titration curves for unfucosylated and di-fucosylated Asialo-AGP glycan compositions with warfarin.………………………………. ……………………………S51

**Figure S7.** Titration curves for unfucosylated and mono-core-fucosylated AD-asialo-AGP glycan compositions with warfarin.…………………………… …………………………………S52

**Figure S8.** van't Hoff plots for unfucosylated and di-fucosylated species in asialo-AGP F1 variant measured by vT-nMS ……………………………………………………………………..S53

**Figure S9.** van't Hoff plots for unfucosylated and mono-fucosylated species in AD-asialo-AGP F1 variant measured by vT-nMS……………………………………………………………S54

**Figure S10.** Representative native mass spectra of (a) asialo-AGP and (b) AD-asialo-AGP in the absence and presence of warfarin..…………… ………………………………………..………S55

**Figure S11.** Titration curves for unfucosylated and mono-core-fucosylated AG-AD-asialo-AGP F1 variant glycan compositions with warfarin.…………………………………………………S56

**Figure S12.** van't Hoff plots for unfucosylated and mono-core-fucosylated species in AG-AD-asialo-AGP F1 variant measured by vT-nMS……………………………………………….…S57

**Figure S13.** Binding and thermodynamic analysis of F1 variant glycosylation-dependent warfarin binding to AG-AD-asialo-AGP ……………………………………………………...………S58

**Figure S14.** Structural characterization and thermodynamic analysis of glycosylation-dependent warfarin binding to TG-AG-AD-asialo-AGP. ………………………………………..……………S59

**Figure S15.** Backbone RMSD along the GaMD trajectory of the RosettaFold-modeled AGP system.……………………………………………………………….………………………S60

**Figure S16.** Hydrogen-bond persistence between the Asn38 *N*-glycan and residues near the funnel entrance.……………………………………………………...……………………………S61

**Figure S17.** Comparison of hydrogen-bond persistence between the S and F1 AGP variants.……………………………………………………...…………..…………………S62

**References**………………………...……………………………..…………………...…...S63

**Materials and Methods**

**Separation of AGP variants.** Chromatography was performed at room temperature using HiTrap® IMAC Fast Flow (7 mm i.d., bed volume 1 mL) purchased from Sigma-Aldrich Canada. The column was loaded with 0.1 M copper sulfate solution and equilibrated with 200 mM ammonium acetate (AmAc, pH 7.6). Human AGP containing a mixture of the F1, S and A variants (2 mg/mL) was applied to the column at a constant flow rate of 1 mL/min. Fractions of 1 mL were collected. After elution of the unbound variants (predominantly the F1 and S variants) using 200 mM AmAc (pH 7.6), a second buffer (20 mM imidazole in 200 mM ammonium acetate, pH 7.6) was applied to remove the bound A variant. Protein stock solutions were buffer-exchanged using centrifugal microconcentrators into aqueous 200 mM AmAc (pH 7.6) using Amicon 0.5 mL microconcentrators (EMD Millipore, Billerica, MA) with a 10 kDa MW cut-off. Protein concentrations were estimated by UV absorption at 280 nm. All stock solutions were stored at -20 °C until used.

**Preparation of enzymatically remodeled AGP samples.** Asialo-AGP was prepared by incubating AGP (F1*S variants) with NeuC in 200 mM ammonium acetate (pH 6.8) at room temperature overnight. AD-asialo-AGP was prepared by incubating Asialo-AGP with α1-3,4 fucosidase in 10X Glycobuffer 1 and 10X recombinant albumin (rHSA) at 37 °C for 2 days. AG-AD-asialo-AGP was prepared by incubating AD-asialo-AGP with β1-4 galactosidase in 10X Glycobuffer 1 overnight at 37°C. TG-AG-AD-asialo-AGP was prepared by incubating AG-AD-asialo-AGP with β-N-Acetylglucosaminidase S in 10X Glycobuffer 1 overnight at 37 °C. CG-TG-AG-AD-asialo-AGP was prepared by incubating TG-AG-AD-asialo-AGP with Endo F3 in 200 mM ammonium acetate (pH 7.4) at room temperature for 2 days.

**Released *N*-glycan analysis by HILIC-UHPLC.** *N*-Glycans were released from the AGP samples by PNGase F digestion. Samples were denatured in buffer containing 0.5% SDS and 40 mM DTT at 100 °C for 10 min. GlycoBuffer 2, NP-40, and PNGase F were then added, and the samples were incubated at 37 °C for 24 h. Released glycans were labeled with procainamide (ProA) as described previously.[1]

HILIC analysis of the glycan libraries was performed using a Thermo Scientific™ Vanquish™ UHPLC system coupled to a fluorescence detector and ESI mass spectrometer (Thermo Q Exactive Orbitrap). The glycans were separated by HILIC using a Waters BEH Glycan column 150x2.1 mm i.d, with BEH particles size of 1.7 µm. The eluents were ammonium formate 100 mM (pH 4.5) (A) and acetonitrile (B). Separation was performed at a flow rate of 0.2 mL $min^{-1}$ with following gradients: 0-40 min, 75-60% B; 40-75 min, 60-55% B; 75-86 min, 55-50% B. The injection volume was 4 µL. The column compartment temperature was maintained at 60 °C. The fluorescence excitation and emission wavelengths were $\lambda_{ex}$ = 310 and $\lambda_{em}$ = 370 nm. The ESI-MS parameters were as follows: sheath gas flow rate, 40 arb, capillary temperature, 275 °C, probe heater temperature, 250 °C, aux gas flowrate,10 arb, spray voltage, 3.5 kV. The m/z range was 250-3000 in positive mode. The resolution for full MS analysis was at 70000. The maximum injection time was 100 ms and the automatic gain control (AGC) target was at $1x10^6$. The MS spectra were recorded using Xcalibur (Thermo, Version 4.1). The glycan composition corresponding to each peak identified by HILIC analysis was preliminarily assigned according to the MS, MS/MS data with the aid of the Glycoworkbench software and known biosynthetic pathways of *N*-glycan.[2] Structural assignments of separated isomers were further supported by glucose-unit (GU) retention indices.[3,4]

**Glycopeptide analysis by nanoLC-MS/MS.** For each analysis, 100 pmol of AGP in 200 mM ammonium acetate, pH 7.4, was reduced with 10 mM DTT at 60 °C for 1 h and subsequently alkylated with 20 mM iodoacetamide in the dark at room temperature for 45 min. Samples were digested with 2 μg thermolysin at 65 °C for 12 h. The digests were dried overnight in a SpeedVac and reconstituted in 50 μL of water containing 3% acetonitrile and 0.1% formic acid.

NanoLC-MS was performed on a Waters NanoACQUITY UPLC system (Waters, Milford, MA) coupled with 15T solariX-XR Fourier-transform ion cyclotron resonance (FT-ICR) mass spectrometer (Bruker-Daltonics, Billerica, MA). Mobile phases A and B were water and acetonitrile, respectively, each containing 0.1% (v/v) formic acid. For each injection, a 3 μL sample solution was first injected onto an ACQUITY UPLC M-Class Symmetry C18 trap column (180 μm i.d., 20 mm length, particle size 5 μm) with 1% B and a flow rate of 4 μL $min^{-1}$ for 4 min. After trapping, the solution was loaded onto a Waters ACQUITY UPLC M-Class Peptide BEH C18 nanoLC column (75 μm i.d., 150 mm length, with BEH particles size of 1.7 μm). Peptides were separated at 35 °C with a flow rate of 0.3 μL $min^{-1}$ and the gradient of 1–40% B in 110 min. The FT-ICR mass spectrometer was operated in data-dependent acquisition mode (positive ion mode) using FtmsControl software (version 2.2). The ions/droplets were transferred to the mass spectrometer using a nanoESI silica emitter of 10 μm tip i.d. (New Objective, Littleton, MA), with 2 kV voltage and a source drying gas ($N_2$, 1.2 L min-1 and 180 °C) to aid with desolvation. The ions were steered by a deflector (220 V) into the funnel (150 V)–skimmer (30 V) stage, and accumulated in a hexapole for 0.2 s (MS) or 1.0 s (MS/MS). For each MS scan, the top 6 most abundant ions were selected, sequentially isolated in the quadrupole, and subjected to MS/MS fragmentation either by CID in the hexapole collision cell (collision gas Ar at $\sim 7\times10^{-6}$ mbar and collision energies 30–50 V) or by ECD in the Para Cell (ECD pulse length 0.08 s, ECD Bias 1.0 V and ECD lens 15 V). Finally, the ions were detected in the ParaCell (sweep excitation power 30%, pressure $\sim 6\times10^{-10}$ mbar). Transients were acquired with 1M data points. The glycoproteomics data was first analyzed using BioTools version 3.2 (Bruker). The MS1 and MS2 mass tolerance were set to 5 ppm and 0.02 Da, respectively. The identification of (glyco)peptides (i.e., protease cleavage sites) was confirmed by MS/MS fragments followed by manual inspection. Ultimately, the relative distributions of the identified glycopeptides were determined through extracted ion chromatograms using DataAnalysis version 5.0 (Bruker) and a home-built script.

**Isothermal titration calorimetry.** Warfarin binding to untreated AGP and asialo-AGP was measured using a MicroCal PEAQ-ITC instrument. AGP or asialo-AGP (15 μM) in the sample cell was titrated with warfarin (300 μM) at 25 °C. Both solutions were prepared in 200 mM ammonium acetate, pH 7.6. The first injection was 0.4 μL over 0.8 s, followed by 29 injections of 2 μL over 4 s each, with 150 s between injections. The reference power was 5 μW.

**Native mass spectrometry.** Native MS measurements were performed using a Q Exactive UHMR Orbitrap mass spectrometer (Thermo Fisher Scientific, Bremen, Germany). Standard and submicron emitters were produced from borosilicate capillary glass with filament (1.0 mm outside diameter (o.d.), 0.78 mm inner diameter (i.d.), 10 cm length and 1.2 mm o.d., 0.69 mm i.d., 10 cm length, respectively, Sutter Instruments, CA) using a P-1000 micropipette puller and a P-97 micropipette puller (Sutter Instruments, CA), respectively. To perform nanoESI, a voltage of approximately +0.5 to +0.9 kV was applied to a platinum wire inserted inside the nanoESI tip and in contact with the solution. The solution temperature was 25 °C. A resolution (resolving power) of 12,500 was used for the binding measurement. Maximum injection time was 200 ms, the capillary temperature was 160 °C, the S-lens RF level was 100 and DC offset was 60. Relevant parameters were: low m/z detector optimization, low m/z ion transfer target, trapping gas pressure 3. Raw data were processed using the Thermo Xcalibur 4.2 software. Mass spectra were averaged over 1 min intervals and the sum of the charge state-normalized abundances of the reactant and the complex ions were calculated automatically using the SWARM software (https://github.com/pkitov/CUPRA-SWARM).[5]

**Temperature-controlled nanoESI device.** The variable-temperature (vT)-nMS measurements were performed using a custom-built device that precisely controlled the temperature of the solution in the nanoESI emitter.[6] Solution temperature inside the emitter was monitored with a T-type thermocouple and controlled within ±0.5 °C, over the range of 0–60 °C. Sub-ambient temperatures were achieved using cooled nitrogen gas (flow rate ~5 L $min^{-1}$), which was passed through copper tubing in contact with a dry ice/ethanol bath (-72 °C), whereas elevated temperatures were generated using embedded cartridge heaters. The nanoESI emitter was positioned ~5 mm from the MS inlet and operated with a source temperature of 60 °C to minimize heat transfer from the sampling capillary.

**Deconvolution of ESI mass spectra.** Deconvolution of the ESI mass spectra was performed using UniDec software (version 4.4.1)[7] for AGP samples and with warfarin. The following parameters were used: m/z range 2,400–5,000; bin every 0.5 (i.e., the raw MS data points were resampled at every 0.5 m/z to UniDec); charge range 6–14; mass range 22,000-50,000; sample mass every 5.0 Da; Peak FWHM (Th) 0.8; peak shape function Gaussian; and maximum number of iterations 100. A full description of the UniDec deconvolution parameters can be found from the User Manual at http://unidec.chem.ox.ac.uk/

**Determination of composition-resolved binding affinities and correction for in-source dissociation.** Warfarin binding to mass-resolved AGP compositions was quantified by direct nMS analysis. AGP samples were analyzed over a range of warfarin concentrations in 200 mM ammonium acetate containing 1% DMSO at pH 7.6. Only complexes containing one warfarin molecule per AGP molecule were detected; accordingly, the binding data were analyzed using a

1:1 binding model.[8] Each resolved AGP composition may contain multiple structural or site-specific glycoforms, and the resulting dissociation constant therefore describes binding by that composition without resolving contributions from its constituent glycoforms.

For a single protein species P binding one ligand L, the binding equilibrium is:

$$\mathrm{P} + \mathrm{L} \rightleftharpoons \mathrm{PL} \tag{S1}$$

and the dissociation constant $K_d$ is:

$$K_\mathrm{d} = \frac{[\mathrm{P}][\mathrm{L}]}{[\mathrm{PL}]} = \frac{[\mathrm{L}]_0}{R} - \frac{[\mathrm{P}]_0}{1+R} \tag{S2}$$

where $[P]_0$ and $[L]_0$ are the total protein and ligand concentrations, respectively and $R$ is the equilibrium concentration ratio of ligand-bound to free protein in solution:

$$R = \frac{[\mathrm{PL}]}{[\mathrm{P}]} \tag{S3a}$$

In the absence of non-specific ligand association and gas-phase dissociation of complex during analysis, and assuming equal detection responses for P and PL ions, the total gas-phase ion abundance (*Ab*) ratio ($R_{app}$) of PL to P ions measured by nMS provides an estimate of $R$:

$$R_\mathrm{app} = \frac{Ab(\mathrm{PL})}{Ab(\mathrm{P})} \approx \frac{[\mathrm{PL}]}{[\mathrm{P}]} = R \tag{S3b}$$

For an AGP sample containing multiple mass-resolved compositions, each composition ($P_i$) can bind warfarin, eq S4:[9]

$$\mathrm{P_i} + \mathrm{L} \rightleftharpoons \mathrm{P_iL} \tag{S4}$$

All compositions share the same equilibrium free-ligand concentration. Accounting for ligand bound to all compositions, the affinity of a given $P_i$ species ($K_{d,i}$) can be calculated from eq S5:

$$K_\mathrm{d,i} = \frac{[\mathrm{P_i}][\mathrm{L}]}{[\mathrm{P_iL}]} = \frac{[\mathrm{L}]_0}{R_\mathrm{i}} - \frac{1}{R_\mathrm{i}} \sum_\mathrm{j} \frac{R_\mathrm{j}[\mathrm{P_j}]_0}{1+R_\mathrm{j}} \tag{S5}$$

The solution concentration ratio and corresponding measured ion-abundance ratio are defined by eqs S6a and S6b, respectively:

$$R_\mathrm{i} = \frac{[\mathrm{P_iL}]}{[\mathrm{P_i}]} \tag{S6a}$$

$$R_\mathrm{i\_app} = \frac{Ab(\mathrm{P_iL})}{Ab(\mathrm{P_i})} \approx \frac{[\mathrm{P_iL}]}{[\mathrm{P_i}]} = R_\mathrm{i} \tag{S6b}$$

and $Ab(\mathrm{P_iL})$ and $Ab(\mathrm{P_i})$ are the total ion abundances of $\mathrm{P_iL}$ and $\mathrm{P_i}$, respectively.

We compared this competitive treatment with a simplified single-species binding model and found that the two approaches provided approximately equivalent affinity estimates under the experimental conditions examined. The simplified model was therefore used for the composition-resolved titration analysis described below.

Under the simplified single-species approximation, the solution-phase fraction of ligand-bound AGP for composition i ($F_\mathrm{i}$) is described by:

$$F_\mathrm{i} = \frac{R_\mathrm{i}}{R_\mathrm{i}+1} = \frac{[\mathrm{P_i}]_0 + [\mathrm{L}]_0 + K_\mathrm{d,i} - \sqrt{(K_\mathrm{d,i} - [\mathrm{L}]_0 + [\mathrm{P_i}]_0)^2 + 4K_\mathrm{d,i}[\mathrm{L}]_0}}{2[\mathrm{P_i}]_0} \tag{S7}$$

where $[\mathrm{P_i}]_0$ ($= f_\mathrm{i}[\mathrm{P}]_0$) is the total concentration of composition i.

Partial in-source dissociation of AGP–warfarin complexes during nMS analysis reduces the measured bound fraction relative to its solution-phase value. To account for this effect, the experimentally observed (by nMS) bound fraction, $F_\mathrm{i\_app}$ was modeled as:

$$F_\mathrm{i\_app} = \frac{R_\mathrm{i\_app}}{1 + R_\mathrm{i\_app}} = (1 - d_{f,\mathrm{i}})F_\mathrm{i} \tag{S8}$$

where $d_{f,\mathrm{i}}$ is the fraction of complexes of composition i that dissociate during analysis. The dissociation constant $K_\mathrm{d,i}$ and dissociation fraction $d_{f,\mathrm{i}}$ were determined by nonlinear regression of eq S8 against the measured bound fractions as a function of total warfarin concentration, with the fitted saturation limit being 1- $d_{f,\mathrm{i}}$. This model assumes that this fraction is independent of warfarin concentration under a given set of measurement conditions and that dissociation products contribute to the measured free-protein signal.

To assess whether a concentration-independent dissociation fraction adequately described the titration data, the fitted $d_{f,\mathrm{i}}$ value was used to calculate a corrected bound fraction at each warfarin concentration, $F_{\mathrm{i\_corr}}$:

$$F_{\mathrm{i\_corr}} = \frac{F_{\mathrm{i\_app}}}{1 - d_{f,\mathrm{i}}} \tag{S9}$$

The corresponding corrected bound-to-free ratio, $R_{\mathrm{i\_corr}}$, was calculated as:

$$R_{\mathrm{i\_corr}} = \frac{F_{\mathrm{i\_corr}}}{1 - F_{\mathrm{i\_corr}}} = \frac{R_{\mathrm{i\_app}}}{1 - d_{f,\mathrm{i}}(R_{\mathrm{i\_app}} + 1)} \tag{S10}$$

Under the same simplified single-species approximation, a dissociation constant was then calculated separately at each warfarin concentration:

$$K_{\mathrm{d,i}} = \frac{[\mathrm{L}]_0}{R_{\mathrm{i\_corr}}} - \frac{[\mathrm{P_i}]_0}{1 + R_{\mathrm{i\_corr}}} \tag{S11}$$

The resulting point-by-point $K_{\mathrm{d,i}}$ estimates were approximately independent of warfarin concentration and agreed with the value obtained by nonlinear regression of the complete titration (Figure 2b). This agreement supports the internal consistency of the correction and the use of a concentration-independent dissociation fraction over the concentration range examined.

The apparent Gibbs free energy of warfarin binding to composition i at temperature $T$ was calculated from the corrected dissociation constant:

$$\Delta G_{\mathrm{i}} = \mathrm{R}T\ln\left(K_{\mathrm{d,i}}/c^{o}\right) \tag{S12}$$

where R is the ideal gas constant and $c^{o}$ (1 mol $\mathrm{L}^{-1}$) is the standard concentration. The thermodynamic parameters reported here describe binding under the specified solution conditions and were determined without activity corrections.

Assuming that the binding enthalpy ($\Delta H_i$) and entropy ($\Delta S_i$) were approximately independent of temperature over the range examined, their values were determined using the linear van't Hoff relationship:

$$\ln\left(K_{d,i}/c^{o}\right) = \frac{\Delta H_i}{RT} - \frac{\Delta S_i}{R} \tag{S13}$$

Linear regression of $\ln(K_{d,i}/c^{o})$ against $1/T$ yielded $\Delta H_i/R$ from the slope and - $\Delta S_i/R$ from the intercept.

**Comparison with ITC.** To compare the composition-resolved nMS affinities with the ensemble affinity measured by ITC, a population-weighted mean $\overline{K}_{d,app}$ was calculated using eq S14:

$$\overline{K}_{d,app} = \sum_{i} f_i \, K_{d,i} \tag{S14}$$

Here, $f_i$ is the fractional abundance of composition i within the AGP population included in the calculation (where $\sum f_i = 1$) and $K_{d,i}$ is its corrected dissociation constant. This weighted mean summarizes the composition-resolved affinities and was compared with the dissociation constant obtained from the single-site ITC analysis.

Similarly, to compare the composition-resolved binding thermodynamics obtained by vT-nMS with the ensemble thermodynamic parameters measured by ITC, population-weighted mean enthalpy ($\overline{\Delta H}_{app}$) and entropy ($\overline{\Delta S}_{app}$) changes were calculated using eq S15a and S15b, respectively:

$$\overline{\Delta H}_{app} = \sum_{i} f_i \, \Delta H_i \tag{S15a}$$

$$\overline{\Delta S}_{app} = \sum_{i} f_i \, \Delta S_i \tag{S15b}$$

Here, $\Delta H_i$ and $\Delta S_i$ are the binding enthalpy and entropy changes for composition i, respectively, determined by van't Hoff analysis. These population-weighted means were compared with the apparent enthalpy and entropy changes obtained from single-site ITC analysis.

**Computational Methods**

**MD simulations.** AGP glycoforms were generated using the ReGlyco tool in GlycoShape (https://glycoshape.org)[11]. The PDB 7OUB structure of AGP (1.82 Å), corresponding to the A variant, was initially selected to investigate the conformational stability of the α-helix containing the glycosylation site at Asn38. To do this we generated two systems, one with AGP glycosylated with a Man5 (GlyTouCan G00028MO) at Asn38 and the other glycosylated at all sites and with a core-fucosylated N-glycan (GlyTouCan G87921DA) at Asn38. We used GlycoShape ReGlyco to reconstruct all glycoforms and GlycoShape GlySys to prepare the systems for MD simulations. Each glycoprotein was centered in a cubic simulation box with a minimum distance of 12 Å between the glycoprotein and the box boundary. The simulation box was solvated with TIP3P water,[12] and 150 mM of NaCl. The protein and counterions, and glycan atoms were represented with the Amberff14SB[13] and GLYCAM06j-1[14], respectively. The systems were relaxed through 500k steps of steepest-descent energy minimization with positional restraints applied to the heavy atoms of the protein and glycans. The systems were subsequently heated from 0 to 300 K over 1 ns in the NVT ensemble using Langevin dynamics, followed by 500 ps of NPT equilibration at 1 atm using a Berendsen barostat. All positional restraints were then removed before production simulations. Gaussian accelerated molecular dynamics (GaMD) simulations[15] were performed using the implementation in AMBER24[16] in dual-boost mode, with independent boost potentials applied to the dihedral and total potential energies. The reference energy was set to the lower bound (iE = 1), and the upper limit of the standard deviation of the boost potential was set to 6 kcal mol$^{-1}$ for both the dihedral ($\sigma_0^D$) and total potential ($\sigma_0^P$) boosts. An initial 1 ns conventional MD simulation was used to collect potential energy statistics, with the first 0.2 ns excluded as preprocessing. This was followed by 50 ns of GaMD equilibration, with the first 3.2 ns excluded to allow adaptation to the modified potential. Production GaMD simulations were then performed for 1 μs, continuously updating the boost parameters, with potential-energy averages and standard deviations recalculated every 0.5 ns. Bonds involving hydrogen atoms were constrained using SHAKE, allowing a 2 fs integration timestep, and long-range electrostatic interactions were treated using the particle mesh Ewald (PME) method with a 9 Å cutoff for non-bonded interactions. Secondary-structure evolution during the 1 μs production trajectories was monitored using DSSP.[17] Because the α-helix containing Asn38 did not unfold during GaMD simulations initiated from the crystallographic structure, an alternative structure was built by using RFdiffusion3.[18] In this structure the alpha helix where Asn38 is located was partially unfolded into a loop conformation. Local conformational sampling of the AlphaFold3 AGP structure was performed to generate glycan-compatible alternative conformations around Asn38. Residues between aa 32 and 44 were allowed to vary while the remainder of the protein and the amino-acid sequence remained fixed. Sampling used partial_t = 12 Å, step scale 0.95, noise scale 1.015, and was biased towards exposed, loop-like conformations. Candidates were retained when Asn38 adopted a non-helical conformation and no major geometric defects were detected. Glycan compatibility was then assessed with ReGlyco by jointly modelling glycans at Asn15, Asn38, Asn54, Asn75 and Asn85, retaining only structures for which all five glycans could be accommodated without steric clashes. Models were generated independently for the AGP S and F1 variants. Structures were glycosylated and solvated in 150 mM NaCl using ReGlyco and ran following the same GaMD protocol described above. For the S variant, GaMD simulations were performed for the non-core-fucosylated and core-fucosylated Asn38 glycoforms. The orientation of the Asn38 N-glycan relative to the AGP binding funnel was quantified using a glycan tilt angle (θ) defined relative to a plane constructed from the Cα atoms of residues 67–69, 71–73, and 91–92 and calculated from the resulting trajectories using a python script. To assess the dependence of the results on the AGP variant and choice of starting structure, the F1 variant was examined using two approaches. First, by introducing the R20Q mutation in S, while preserving the rest of the conformation. Second, by generating a new RoseTTAFold model corresponding to the F1 variant, which was used as a starting structure for the GaMD simulation ran following the same protocol. Protein stability and local conformational dynamics were assessed from backbone root mean square deviation (RMSD) values and Cα root mean square fluctuation (RMSF) values calculated with scripts run on the Visual Molecular Dynamics (VMD) tkl console.[19]

**Conformational ensemble analysis of AGP glycoforms.** To complement the GaMD simulations and characterize the conformational behaviour of the experimentally investigated *N*-glycans, we rebuilt conformational ensembles at each glycosylation site using ReGlyco Ensemble module in GlycoShape.[11,20] To rebuild each one of the 18 different glycoforms we tested, we sourced 200 glycan conformations for each type of glycan from the GlycoShape database and linked them at each occupied site. The analysis we performed encompasses a selected group of AGP glycoforms that recapitulates salient features of the experimentally characterised ensembles, namely a reference glycoform (Table S1), core-fucosylated, and bisected glycoforms, together with a systematic panel of antenna-fucosylated glycoforms differing in glycan branching, glycosylation site, and the specific antenna functionalized. The antenna-fucosylated systems are indicated as Models 1–12 with glycan compositions shown in Table S1. All glycoforms were rebuilt with the ReGlyco Ensemble tool in GlycoShape using 200 frames for each selected glycan at each site selected from the MD database. Glycan–protein interactions were quantified using the Interactions plugin in VMD.[19]

**Tables**

**Table S1.** AGP glycoform nomenclature.

*Reference, core-fucosylated, and bisecting glycoforms (variant S)*

| System | Asn38 glycan | Asn75 glycan | Description |
|---|---|---|---|
| **S-G0** | Tri-antennary, no fuc | Tetra-antennary, no fuc | Unfucosylated reference |
| **S-G1** | Tri-antennary, core-fuc | Tetra-antennary, no fuc | Core-fucosylation at Asn38 only |
| **S-G2** | Tri-antennary, no fuc | Tetra-antennary, core-fuc | Core-fucosylation at Asn75 only |
| **S-G3** | Tri-antennary, core-fuc | Tetra-antennary, core-fuc | Core-fucosylation at both sites |
| **S-G22** | Trimmed | Tetra-antennary, bisecting GlcNAc | Truncated Asn38 + bisecting at Asn75 |
| **S-G0-bisec75** | Tri-antennary, no fuc | Tetra-antennary, bisecting GlcNAc | Bisecting GlcNAc at Asn75 |

*Antennary-fucosylated glycoforms at Asn75 (variant F1 and S)*

| System | Asn38 glycan | Asn75 glycan | Description |
|---|---|---|---|
| **Model 1** | — | Tri-antennary, antennary-fuc | Antennary fucose at Asn75, α1,3 arm |
| **Model 2** | — | Tri-antennary, antennary-fuc | Antennary fucose at Asn75, α1,6 arm |
| **Model 3** | — | Tetra-antennary, antennary-fuc | Antennary fucose at Asn75, α1,3 arm |
| **Model 4** | — | Tetra-antennary, antennary-fuc | Antennary fucose at Asn75, α1,6 arm |
| **Model 5** | Bi-antennary, antennary-fuc | Tri-antennary, antennary-fuc | Antennary fucose at both sites, α1,3 arm |
| **Model 6** | Bi-antennary, antennary-fuc | Tri-antennary, antennary-fuc | Antennary fucose at both sites, α1,6 arm |
| **Model 7** | Tri-antennary, antennary-fuc | Tetra-antennary, antennary-fuc | Antennary fucose at both sites, α1,3 arm |

| | | | |
|---|---|---|---|
| **Model 8** | Tri-antennary, antennary-fuc | Tetra-antennary, antennary-fuc | Antennary fucose at both sites, α1,6 arm |

*Antennary-fucosylated glycoforms at Asn38 (variant F1 and S)*

| System | Asn38 glycan | Asn75 glycan | Description |
|---|---|---|---|
| **Model 9** | Tri-antennary, antennary-fuc | Tetra-antennary, no fuc | Antennary fucose at Asn38 only, α1,3 arm |
| **Model 10** | Tri-antennary, antennary-fuc | Tetra-antennary, no fuc | Antennary fucose at Asn38 only, α1,6 arm |
| **Model 11** | Bi-antennary, antennary-fuc | Tri-antennary, no fuc | Antennary fucose at Asn38 only, α1,3 arm |
| **Model 12** | Bi-antennary, antennary-fuc | Tri-antennary, no fuc | Antennary fucose at Asn38 only, α1,6 arm |

*Glycoforms modeled for the F1 variant*

| System | Asn38 glycan | Asn75 glycan | Description |
|---|---|---|---|
| **F1-G0** | Tri-antennary, no fuc | Tetra-antennary, no fuc | Unfucosylated reference |
| **F1-G1** | Tri-antennary, core-fuc | Tetra-antennary, no fuc | Core-fucosylation at Asn38 only |
| **F1-G2** | Tri-antennary, no fuc | Tetra-antennary, core-fuc | Core-fucosylation at Asn75 only |
| **F1-G3** | Tri-antennary, core-fuc | Tetra-antennary, core-fuc | Core-fucosylation at both sites |
| **F1-G22** | Trimmed | Tetra-antennary, bisecting GlcNAc | Truncated Asn38 + bisecting at Asn75 |
| **F1-G0-bisec75** | Tri-antennary, no fuc | Tetra-antennary, bisecting GlcNAc | Bisecting GlcNAc at Asn75 |

**Table S2.** Annotation of AGP glycan compositions.

| Num. | Experimental Mass (Da)[a] | Theoretical Mass (Da)[b] | Mass Difference (Da) | Relative Abundance (%) | Composition [c] | Genetic Variant | Annotation |
|---|---|---|---|---|---|---|---|
| 1 | 34685.78 | 34684.36 | 1.42 | 10.36% | 28_23_13.5 | F1 | P0 |
| 2 | 34903.91 | 34904.07 | 0.16 | 14.20% | 29_24_13 | F1 | P1 |
| 3 | 34932.16 | 34932.12 | 0.04 | 4.49% | 29_24_13 | S | P1 |
| 4 | 35050.44 | 35049.69 | 0.75 | 22.03% | 29_24_13.5 | F1 | P1 |
| 5 | 35075.01 | 35077.74 | 2.73 | 5.23% | 29_24_13.5 | S | P1 |
| 6 | 35118.41 | 35121.24 | 2.83 | 9.94% | 28_23_15 | F1 | P0 |
| 7 | 35149.72 | 35149.29 | 0.43 | 6.56% | 28_23_15 | S | P0 |
| 8 | 35196.08 | 35195.32 | 0.75 | 28.81% | 29_24_14 | F1 | P1 |
| 9 | 35220.10 | 35223.37 | 3.27 | 8.18% | 29_24_14 | S | P1 |
| 10 | 35268.86 | 35269.40 | 0.54 | 27.16% | 30_25_13 | F1 | P2 |
| 11 | 35298.40 | 35297.45 | 0.95 | 9.97% | 30_25_13 | S | P2 |
| 12 | 35342.03 | 35340.95 | 1.08 | 24.56% | 29_24_14.5 | F1 | P1 |
| 13 | 35367.71 | 35369.00 | 1.29 | 6.28% | 29_24_14.5 | S | P1 |
| 14 | 35416.36 | 35415.03 | 1.33 | 16.42% | 30_25_13.5 | F1 | P2 |
| 15 | 35441.91 | 35443.08 | 1.16 | 11.59% | 30_25_13.5 | S | P2 |
| 16 | 35488.41 | 35486.58 | 1.84 | 18.57% | 29_24_15 | F1 | P1 |
| 17 | 35513.35 | 35514.63 | 1.28 | 6.37% | 29_24_15 | S | P1 |
| 18 | 35561.10 | 35560.66 | 0.44 | 48.91% | 30_25_14 | F1 | P2 |
| 19 | 35586.92 | 35588.71 | 1.78 | 18.02% | 30_25_14 | S | P2 |
| 20 | 35634.85 | 35634.73 | 0.12 | 35.58% | 31_26_13 | F1 | P3 |
| 21 | 35659.74 | 35662.78 | 3.04 | 11.31% | 31_26_13 | S | P3 |
| 22 | 35707.52 | 35706.28 | 1.24 | 38.16% | 30_25_14.5 | F1 | P2 |
| 23 | 35733.55 | 35734.33 | 0.79 | 13.05% | 30_25_14.5 | S | P2 |

| | | | | | | | |
|---|---|---|---|---|---|---|---|
| 24 | 35781.92 | 35777.83 | 4.09 | 15.92% | 29_24_16 | F1 | P1 |
| 25 | 35809.50 | 35805.88 | 3.62 | 10.18% | 29_24_16 | S | P1 |
| 26 | 35852.46 | 35851.91 | 0.55 | 64.13% | 30_25_15 | F1 | P2 |
| 27 | 35878.34 | 35879.96 | 1.62 | 28.18% | 30_25_15 | S | P2 |
| 28 | 35926.31 | 35925.99 | 0.32 | 56.52% | 31_26_14 | F1 | P3 |
| 29 | 35952.26 | 35954.04 | 1.78 | 24.52% | 31_26_14 | S | P3 |
| 30 | 35999.09 | 36000.07 | 0.98 | 60.52% | 32_27_13 | F1 | P4 |
| 31 | 36024.62 | 36028.12 | 3.49 | 23.24% | 32_27_13 | S | P4 |
| 32 | 36073.28 | 36071.62 | 1.66 | 29.56% | 31_26_14.5 | F1 | P3 |
| 33 | 36100.15 | 36099.67 | 0.48 | 19.70% | 31_26_14.5 | S | P3 |
| 34 | 36145.52 | 36143.16 | 2.36 | 37.32% | 30_25_16 | F1 | P2 |
| 35 | 36173.23 | 36171.21 | 2.02 | 19.10% | 30_25_16 | S | P2 |
| 36 | 36217.54 | 36217.24 | 0.29 | 81.26% | 31_26_15 | F1 | P3 |
| 37 | 36243.72 | 36245.29 | 1.58 | 40.86% | 31_26_15 | S | P3 |
| 38 | 36291.54 | 36291.32 | 0.22 | 73.44% | 32_27_14 | F1 | P4 |
| 39 | 36317.23 | 36319.37 | 2.14 | 31.51% | 32_27_14 | S | P4 |
| 40 | 36364.29 | 36365.40 | 1.11 | 58.90% | 33_28_13 | F1 | P5 |
| 41 | 36390.59 | 36393.45 | 2.86 | 29.46% | 33_28_13 | S | P5 |
| 42 | 36438.06 | 36434.42 | 3.64 | 35.65% | 30_25_17 | F1 | P2 |
| 43 | 36465.11 | 36462.47 | 2.64 | 25.00% | 30_25_17 | S | P2 |
| 44 | 36509.21 | 36508.50 | 0.72 | 76.43% | 31_26_16 | F1 | P3 |
| 45 | 36535.87 | 36536.55 | 0.67 | 40.80% | 31_26_16 | S | P3 |
| 46 | 36582.84 | 36582.58 | 0.27 | 95.11% | 32_27_15 | F1 | P4 |
| 47 | 36609.25 | 36610.63 | 1.37 | 49.75% | 32_27_15 | S | P4 |
| 48 | 36655.79 | 36656.65 | 0.86 | 75.68% | 33_28_14 | F1 | P5 |

| | | | | | | | |
|---|---|---|---|---|---|---|---|
| 49 | 36682.33 | 36684.70 | 2.38 | 37.28% | 33_28_14 | S | P5 |
| 50 | 36729.57 | 36728.20 | 1.36 | 51.89% | 32_27_15.5 | F1 | P4 |
| 51 | 36756.67 | 36756.25 | 0.42 | 36.83% | 32_27_15.5 | S | P4 |
| 52 | 36802.20 | 36799.75 | 2.45 | 44.92% | 31_26_17 | F1 | P3 |
| 53 | 36830.24 | 36827.80 | 2.44 | 30.37% | 31_26_17 | S | P3 |
| 54 | 36874.08 | 36873.83 | 0.25 | 100.00% | 32_27_16 | F1 | P4 |
| 55 | 36900.88 | 36901.88 | 1.00 | 54.53% | 32_27_16 | S | P4 |
| 56 | 36947.87 | 36947.91 | 0.04 | 79.45% | 33_28_15 | F1 | P5 |
| 57 | 36974.46 | 36975.96 | 1.50 | 45.05% | 33_28_15 | S | P5 |
| 58 | 37020.66 | 37021.99 | 1.33 | 70.35% | 34_29_14 | F1 | P6 |
| 59 | 37047.89 | 37050.04 | 2.15 | 39.01% | 34_29_14 | S | P6 |
| 60 | 37094.34 | 37091.01 | 3.33 | 41.97% | 31_26_18 | F1 | P3 |
| 61 | 37122.18 | 37119.06 | 3.12 | 35.23% | 31_26_18 | S | P3 |
| 62 | 37166.09 | 37165.09 | 1.01 | 75.37% | 32_27_17 | F1 | P4 |
| 63 | 37193.38 | 37193.14 | 0.25 | 41.65% | 32_27_17 | S | P4 |
| 64 | 37239.58 | 37239.16 | 0.41 | 85.83% | 33_28_16 | F1 | P5 |
| 65 | 37266.32 | 37267.21 | 0.90 | 51.32% | 33_28_16 | S | P5 |
| 66 | 37312.41 | 37313.24 | 0.84 | 66.49% | 34_29_15 | F1 | P6 |
| 67 | 37339.49 | 37341.29 | 1.80 | 36.46% | 34_29_15 | S | P6 |
| 68 | 37385.88 | 37384.79 | 1.09 | 52.69% | 33_28_16.5 | F1 | P5 |
| 69 | 37413.42 | 37412.84 | 0.58 | 38.56% | 33_28_16.5 | S | P5 |
| 70 | 37458.43 | 37456.34 | 2.09 | 41.91% | 32_27_18 | F1 | P4 |
| 71 | 37486.96 | 37484.39 | 2.57 | 28.44% | 32_27_18 | S | P4 |
| 72 | 37531.12 | 37530.42 | 0.70 | 76.54% | 33_28_17 | F1 | P5 |
| 73 | 37558.47 | 37558.47 | 0.00 | 44.60% | 33_28_17 | S | P5 |

| | | | | | | | |
|---|---|---|---|---|---|---|---|
| 74 | 37604.41 | 37604.50 | 0.09 | 51.00% | 34_29_16 | F1 | P6 |
| 75 | 37632.27 | 37632.55 | 0.27 | 31.08% | 34_29_16 | S | P6 |
| 76 | 37677.34 | 37678.58 | 1.23 | 59.40% | 35_30_15 | F1 | P7 |
| 77 | 37705.00 | 37706.63 | 1.63 | 35.79% | 35_30_15 | S | P7 |
| 78 | 37750.99 | 37747.59 | 3.40 | 29.43% | 32_27_19 | F1 | P4 |
| 79 | 37779.04 | 37775.64 | 3.40 | 24.50% | 32_27_19 | S | P4 |
| 80 | 37822.94 | 37821.67 | 1.27 | 54.52% | 33_28_18 | F1 | P5 |
| 81 | 37850.74 | 37849.72 | 1.01 | 31.82% | 33_28_18 | S | P5 |
| 82 | 37896.10 | 37895.75 | 0.34 | 37.85% | 34_29_17 | F1 | P6 |
| 83 | 37923.91 | 37923.80 | 0.10 | 25.32% | 34_29_17 | S | P6 |
| 84 | 37969.16 | 37969.83 | 0.67 | 52.49% | 35_30_16 | F1 | P7 |
| 85 | 37996.84 | 37997.88 | 1.04 | 28.76% | 35_30_16 | S | P7 |
| 86 | 38042.84 | 38043.91 | 1.07 | 25.74% | 36_31_15 | F1 | P8 |
| 87 | 38070.99 | 38071.96 | 0.97 | 20.96% | 36_31_15 | S | P8 |
| 88 | 38115.45 | 38112.93 | 2.53 | 36.26% | 33_28_19 | F1 | P5 |
| 89 | 38143.29 | 38140.98 | 2.31 | 21.49% | 33_28_19 | S | P5 |
| 90 | 38188.28 | 38187.01 | 1.27 | 23.64% | 34_29_18 | F1 | P6 |
| 91 | 38216.82 | 38215.06 | 1.77 | 18.97% | 34_29_18 | S | P6 |
| 92 | 38261.11 | 38261.08 | 0.03 | 38.69% | 35_30_17 | F1 | P7 |
| 93 | 38288.95 | 38289.13 | 0.18 | 21.55% | 35_30_17 | S | P7 |
| 94 | 38334.63 | 38335.16 | 0.53 | 24.60% | 36_31_16 | F1 | P8 |
| 95 | 38362.66 | 38363.21 | 0.56 | 16.67% | 36_31_16 | S | P8 |
| 96 | 38407.35 | 38404.18 | 3.16 | 27.09% | 33_28_20 | F1 | P5 |
| 97 | 38435.78 | 38432.23 | 3.54 | 16.36% | 33_28_20 | S | P5 |
| 98 | 38480.52 | 38478.26 | 2.26 | 17.13% | 34_29_19 | F1 | P6 |

| | | | | | | | |
|---|---|---|---|---|---|---|---|
| 99 | 38509.14 | 38506.31 | 2.83 | 12.93% | 34_29_19 | S | P6 |
| 100 | 38552.96 | 38552.34 | 0.62 | 23.37% | 35_30_18 | F1 | P7 |
| 101 | 38581.41 | 38580.39 | 1.02 | 14.31% | 35_30_18 | S | P7 |
| 102 | 38626.35 | 38626.42 | 0.07 | 22.31% | 36_31_17 | F1 | P8 |
| 103 | 38654.66 | 38654.47 | 0.19 | 14.21% | 36_31_17 | S | P8 |
| 104 | 38699.66 | 38700.50 | 0.83 | 21.80% | 37_32_16 | F1 | P9 |
| 105 | 38727.76 | 38728.55 | 0.79 | 12.49% | 37_32_16 | S | P9 |
| 106 | 38844.96 | 38843.59 | 1.37 | 13.85% | 35_30_19 | F1 | P7 |
| 107 | 38872.28 | 38871.64 | 0.64 | 10.03% | 35_30_19 | S | P7 |
| 108 | 38918.28 | 38917.67 | 0.60 | 15.34% | 36_31_18 | F1 | P8 |

a. The experimental masses were calculated based on the average MW of glycan compositions at three charge states (+8 to +10).
b. The theoretical masses were calculated from the mass of peptide backbone of AGP (F1 variant: 21539 Da, S variant: 21567 Da) and the average masses of monosaccharides (Hex: 162.14 Da, HexNAc: 203.19 Da, Fuc: 146.14 Da, Sia: 291.26 Da).
c. Glycan compositions are expressed as the number of Hexose (Hex), N-acetylhexosamine (HexNAc), Fucose/2+Sialic acid (Fuc/2+Sia).

**Table S3.** Annotation of asialo-AGP glycan compositions.

| Num. | Experimental Mass (Da)[a] | Theoretical Mass (Da)[b] | Mass Difference (Da) | Relative Abundance (%) | Composition[c] | Genetic Variant | Annotation[d] |
|---|---|---|---|---|---|---|---|
| 1 | 30753.32 | 30752.42 | 0.83 | 1.57% | 28_23_0 | F1 | P0 |
| 2 | 31116.81 | 31117.76 | 1.01 | 19.57% | 29_24_0 | F1 | P1 |
| 3 | 31141.59 | 31145.81 | 4.28 | 2.38% | 29_24_0 | S | P1 |
| 4 | 31262.99 | 31263.90 | 0.98 | 10.21% | 29_24_1 | F1 | P1F1 |
| 5 | 31293.68 | 31291.95 | 1.67 | 3.56% | 29_24_1 | S | P1F1 |
| 6 | 31325.76 | 31320.95 | 4.75 | 3.85% | 29_25_0 | F1 | ★ |
| 7 | 31409.43 | 31410.04 | 0.68 | 8.69% | 29_24_2 | F1 | P1F2 |
| 8 | 31438.29 | 31438.09 | 0.14 | 1.06% | 29_24_2 | S | P1F2 |
| 9 | 31482.12 | 31483.09 | 1.03 | 45.48% | 30_25_0 | F1 | P2 |
| 10 | 31508.60 | 31511.14 | 2.60 | 13.98% | 30_25_0 | S | P2 |
| 11 | 31556.79 | 31556.18 | 0.55 | 6.57% | 29_24_3 | F1 | P1F3 |
| 12 | 31582.39 | 31584.23 | 1.90 | 2.09% | 29_24_3 | S | P1F3 |
| 13 | 31628.57 | 31629.23 | 0.73 | 25.02% | 30_25_1 | F1 | P2F1 |
| 14 | 31655.41 | 31657.28 | 1.94 | 8.08% | 30_25_1 | S | P2F1 |
| 15 | 31690.50 | 31686.28 | 4.21 | 8.51% | 30_26_0 | F1 | ★ |
| 16 | 31701.04 | 31702.32 | 1.28 | 2.72% | 29_24_4 | F1 | P1F4 |
| 17 | 31710.02 | 31714.33 | 4.37 | 0.97% | 30_26_0 | S | ★ |
| 18 | 31775.10 | 31775.37 | 0.33 | 21.17% | 30_25_2 | F1 | P2F2 |
| 19 | 31802.39 | 31803.42 | 1.10 | 5.72% | 30_25_2 | S | P2F2 |
| 20 | 31847.80 | 31848.42 | 0.69 | 72.37% | 31_26_0 | F1 | P3 |
| 21 | 31874.10 | 31876.47 | 2.44 | 27.49% | 31_26_0 | S | P3 |
| 22 | 31921.61 | 31921.51 | 0.04 | 12.51% | 30_25_3 | F1 | P2F3 |
| 23 | 31949.34 | 31949.56 | 0.29 | 5.56% | 30_25_3 | S | P2F3 |

| | | | | | | | |
|---|---|---|---|---|---|---|---|
| 24 | 31994.12 | 31994.57 | 0.50 | 38.54% | 31_26_1 | F1 | P3F1 |
| 25 | 32021.16 | 32022.62 | 1.52 | 16.97% | 31_26_1 | S | P3F1 |
| 26 | 32056.43 | 32051.62 | 4.81 | 6.17% | 31_27_0 | F1 | ★ |
| 27 | 32067.36 | 32067.66 | 0.30 | 2.98% | 30_25_4 | F1 | P2F4 |
| 28 | 32096.04 | 32095.71 | 0.33 | 5.72% | 30_25_4 | S | P2F4 |
| 29 | 32140.78 | 32140.71 | 0.07 | 32.80% | 31_26_2 | F1 | P3F2 |
| 30 | 32167.91 | 32168.76 | 0.85 | 12.20% | 31_26_2 | S | P3F2 |
| 31 | 32213.27 | 32213.76 | 0.48 | 100.00% | 32_27_0 | F1 | P4 |
| 32 | 32239.34 | 32241.81 | 2.47 | 42.57% | 32_27_0 | S | P4 |
| 33 | 32287.12 | 32286.85 | 0.28 | 20.15% | 31_26_3 | F1 | P3F3 |
| 34 | 32314.59 | 32314.90 | 0.30 | 9.69% | 31_26_3 | S | P3F3 |
| 35 | 32359.60 | 32359.90 | 0.30 | 53.28% | 32_27_1 | F1 | P4F1 |
| 36 | 32386.62 | 32387.95 | 1.33 | 26.68% | 32_27_1 | S | P4F1 |
| 37 | 32421.17 | 32416.95 | 4.22 | 8.12% | 32_28_0 | F1 | ★ |
| 38 | 32432.14 | 32432.99 | 0.85 | 4.56% | 31_26_4 | F1 | P3F4 |
| 39 | 32461.27 | 32461.04 | 0.23 | 10.63% | 31_26_4 | S | P3F4 |
| 40 | 32506.21 | 32506.04 | 0.17 | 45.14% | 32_27_2 | F1 | P4F2 |
| 41 | 32533.35 | 32534.09 | 0.74 | 18.88% | 32_27_2 | S | P4F2 |
| 42 | 32578.91 | 32579.09 | 0.18 | 85.33% | 33_28_0 | F1 | P5 |
| 43 | 32605.09 | 32607.14 | 2.05 | 43.02% | 33_28_0 | S | P5 |
| 44 | 32652.36 | 32652.18 | 0.18 | 28.19% | 32_27_3 | F1 | P4F3 |
| 45 | 32679.92 | 32680.23 | 0.31 | 14.535 | 32_27_3 | S | P4F3 |
| 46 | 32725.24 | 32725.23 | 0.01 | 46.84% | 33_28_1 | F1 | P5F1 |
| 47 | 32752.22 | 32753.28 | 1.06 | 28.01% | 33_28_1 | S | P5F1 |
| 48 | 32787.19 | 32782.28 | 4.91 | 8.16% | 33_29_0 | F1 | ★ |

| | | | | | | | |
|---|---|---|---|---|---|---|---|
| 49 | 32797.32 | 32798.32 | 1.01 | 4.94% | 32_27_4 | F1 | P4F4 |
| 50 | 32826.28 | 32826.37 | 0.09 | 13.87% | 32_27_4 | S | P4F4 |
| 51 | 32871.62 | 32871.37 | 0.24 | 39.15% | 33_28_2 | F1 | P5F2 |
| 52 | 32898.77 | 32899.42 | 0.66 | 20.33% | 33_28_2 | S | P5F2 |
| 53 | 32944.77 | 32944.42 | 0.34 | 42.66% | 34_29_0 | F1 | P6 |
| 54 | 32971.17 | 32972.47 | 1.31 | 24.12% | 34_29_0 | S | P6 |
| 55 | 33017.76 | 33017.51 | 0.25 | 25.82% | 33_28_3 | F1 | P5F3 |
| 56 | 33045.07 | 33045.56 | 0.50 | 14.93% | 33_28_3 | S | P5F3 |
| 57 | 33090.84 | 33090.56 | 0.28 | 24.13% | 34_29_1 | F1 | P6F1 |
| 58 | 33118.01 | 33118.61 | 0.60 | 16.18% | 34_29_1 | S | P6F1 |
| 59 | 33150.41 | 33147.62 | 2.79 | 3.58% | 34_30_0 | F1 | ★ |
| 60 | 33164.10 | 33163.66 | 0.44 | 13.54% | 33_28_4 | F1 | P5F4 |
| 61 | 33191.57 | 33191.71 | 0.13 | 12.01% | 33_28_4 | S | P5F4 |
| 62 | 33237.16 | 33236.71 | 0.45 | 17.81% | 34_29_2 | F1 | P6F2 |
| 63 | 33265.07 | 33264.76 | 0.32 | 11.55% | 34_29_2 | S | P6F2 |
| 64 | 33309.81 | 33309.76 | 0.05 | 33.82% | 35_30_0 | F1 | P7 |
| 65 | 33336.84 | 33337.81 | 0.97 | 18.16% | 35_30_0 | S | P7 |
| 66 | 33384.28 | 33382.85 | 1.44 | 10.45% | 34_29_3 | F1 | P6F3 |
| 67 | 33411.49 | 33410.90 | 0.59 | 8.35% | 34_29_3 | S | P6F3 |
| 68 | 33455.79 | 33455.90 | 0.10 | 21.13% | 35_30_1 | F1 | P7F1 |
| 69 | 33483.22 | 33483.95 | 0.73 | 12.24% | 35_30_1 | S | P7F1 |
| 70 | 33517.33 | 33512.95 | 4.38 | 3.96% | 35_31_0 | F1 | ★ |
| 71 | 33527.47 | 33528.99 | 1.52 | 3.71% | 34_29_4 | F1 | P6F4 |
| 72 | 33558.33 | 33557.04 | 1.29 | 6.39% | 34_29_4 | S | P6F4 |
| 73 | 33602.17 | 33602.04 | 0.13 | 16.18% | 35_30_2 | F1 | P7F2 |

| | | | | | | | |
|---|---|---|---|---|---|---|---|
| 74 | 33630.21 | 33630.09 | 0.12 | 8.50% | 35_30_2 | S | P7F2 |
| 75 | 33675.22 | 33675.09 | 0.13 | 18.01% | 36_31_0 | F1 | P8 |
| 76 | 33702.32 | 33703.14 | 0.82 | 10.61% | 36_31_0 | S | P8 |
| 77 | 33748.15 | 33748.18 | 0.03 | 10.13% | 35_30_3 | F1 | P7F3 |
| 78 | 33776.36 | 33776.23 | 0.13 | 5.97% | 35_30_3 | S | P7F3 |
| 79 | 33821.33 | 33821.23 | 0.10 | 10.84% | 36_31_1 | F1 | P8F1 |
| 80 | 33849.31 | 33849.28 | 0.03 | 7.35% | 36_31_1 | S | P8F1 |
| 81 | 33882.90 | 33878.28 | 4.62 | 2.16% | 36_32_0 | F1 | ★ |
| 82 | 33893.71 | 33894.32 | 0.61 | 3.81% | 35_30_4 | F1 | P7F4 |
| 83 | 33923.18 | 33922.37 | 0.81 | 4.46% | 35_30_4 | S | P7F4 |
| 84 | 33967.83 | 33967.37 | 0.46 | 8.90% | 36_31_2 | F1 | P8F2 |
| 85 | 33995.79 | 33995.42 | 0.36 | 5.35% | 36_31_2 | S | P8F2 |
| 86 | 34040.35 | 34040.42 | 0.08 | 12.07% | 37_32_0 | F1 | P9 |
| 87 | 34067.84 | 34068.47 | 0.63 | 6.91% | 37_32_0 | S | P9 |
| 88 | 34113.53 | 34113.51 | 0.01 | 5.97% | 36_31_3 | F1 | P8F3 |
| 89 | 34141.91 | 34141.56 | 0.34 | 3.63% | 36_31_3 | S | P8F3 |
| 90 | 34186.97 | 34186.56 | 0.41 | 6.96% | 37_32_1 | F1 | P9F1 |
| 91 | 34214.16 | 34214.61 | 0.45 | 4.75% | 37_32_1 | S | P9F1 |
| 92 | 34247.32 | 34243.62 | 3.70 | 1.06% | 37_33_0 | F1 | ★ |
| 93 | 34260.08 | 34259.65 | 0.42 | 2.50% | 36_31_4 | F1 | P8F4 |
| 94 | 34286.25 | 34287.70 | 1.46 | 3.09% | 36_31_4 | S | P8F4 |
| 95 | 34333.36 | 34332.71 | 0.66 | 6.60% | 37_32_2 | F1 | P9F2 |
| 96 | 34358.35 | 34360.76 | 2.41 | 3.89% | 37_32_2 | S | P9F2 |
| 97 | 34402.09 | 34405.76 | 3.67 | 7.20% | 38_33_0 | F1 | P10 |
| 98 | 34432.66 | 34433.81 | 1.15 | 5.49% | 38_33_0 | S | P10 |

| | | | | | | | |
|---|---|---|---|---|---|---|---|
| 99 | 34474.50 | 34478.85 | 4.34 | 3.62% | 37_32_3 | F1 | P9F3 |
| 100 | 34505.84 | 34506.90 | 1.05 | 7.38% | 37_32_3 | S | P9F3 |
| 101 | 34547.74 | 34551.90 | 4.16 | 4.61% | 38_33_1 | F1 | P10F1 |
| 102 | 34577.75 | 34579.95 | 2.20 | 12.68% | 38_33_1 | S | P10F1 |
| 103 | 34613.42 | 34608.95 | 4.47 | 0.52% | 38_34_0 | F1 | ★ |
| 104 | 34626.23 | 34624.99 | 1.24 | 1.79% | 37_32_4 | F1 | P9F4 |
| 105 | 34651.67 | 34653.04 | 1.37 | 2.24% | 37_32_4 | S | P9F4 |

a. The experimental masses were calculated based on the average MW of glycan compositions at three charge states (+8 to +10).
b. The theoretical masses were calculated from the mass of peptide backbone of AGP (F1 variant: 21539 Da, S variant: 21567 Da) and the average masses of monosaccharides (Hex: 162.14 Da, HexNAc: 203.19 Da, Fuc: 146.14 Da).
c. Glycan compositions are expressed as the number of Hex, HexNAc, and Fuc residues.
d. Red stars identify compositions consistent with the presence of terminal GlcNAc residues.

**Table S4.** Annotation of Antennary AD-asialo-AGP glycan compositions.

| Num. | Experimental Mass (Da)[a] | Theoretical Mass (Da)[b] | Mass Difference (Da) | Relative Abundance (%) | Composition [c] | Genetic Variant | Annotation[d] |
|---|---|---|---|---|---|---|---|
| 1 | 31116.81 | 31117.76 | 0.95 | 18.0% | 29_24_0 | F1 | P1 |
| 2 | 31141.59 | 31145.81 | 4.22 | 5.4% | 29_24_0 | S | P1 |
| 3 | 31262.99 | 31263.90 | 0.91 | 0.8% | 29_24_1 | F1 | P1F1 |
| 4 | 31293.68 | 31291.95 | 1.73 | 0.9% | 29_24_1 | S | P1F1 |
| 5 | 31325.76 | 31320.95 | 4.81 | 3.2% | 29_25_0 | F1 | ★ |
| 6 | 31349.33 | 31349.00 | 0.33 | 1.2% | 29_25_0 | S | ★ |
| 7 | 31409.43 | 31410.04 | 0.61 | 0.6% | 29_24_2 | F1 | P1F2 |
| 8 | 31438.29 | 31438.09 | 0.20 | 0.2% | 29_24_2 | S | P1F2 |
| 9 | 31482.12 | 31483.09 | 0.97 | 44.2% | 30_25_0 | F1 | P2 |
| 10 | 31508.60 | 31511.14 | 2.54 | 18.2% | 30_25_0 | S | P2 |
| 11 | 31628.57 | 31629.23 | 0.66 | 2.4% | 30_25_1 | F1 | P2F1 |
| 12 | 31655.41 | 31657.28 | 1.88 | 1.3% | 30_25_1 | S | P2F1 |
| 13 | 31691.50 | 31686.28 | 5.21 | 11.2% | 30_26_0 | F1 | ★ |
| 14 | 31710.02 | 31714.33 | 4.31 | 3.6% | 30_26_0 | S | ★ |
| 15 | 31775.10 | 31775.37 | 0.27 | 1.7% | 30_25_2 | F1 | P2F2 |
| 16 | 31802.39 | 31803.42 | 1.03 | 0.7% | 30_25_2 | S | P2F2 |
| 17 | 31847.80 | 31848.42 | 0.63 | 72.2% | 31_26_0 | F1 | P3 |
| 18 | 31874.10 | 31876.47 | 2.38 | 33.1% | 31_26_0 | S | P3 |
| 19 | 31994.12 | 31994.57 | 0.44 | 4.7% | 31_26_1 | F1 | P3F1 |
| 20 | 32021.16 | 32022.62 | 1.46 | 3.5% | 31_26_1 | S | P3F1 |
| 21 | 32057.06 | 32051.62 | 5.45 | 19.6% | 31_27_0 | F1 | ★ |
| 22 | 32078.90 | 32079.67 | 0.77 | 6.0% | 31_27_0 | S | ★ |

| | | | | | | | |
|---|---|---|---|---|---|---|---|
| 23 | 32140.78 | 32140.71 | 0.07 | 2.8% | 31_26_2 | F1 | P3F2 |
| 24 | 32167.91 | 32168.76 | 0.85 | 1.5% | 31_26_2 | S | P3F2 |
| 25 | 32213.27 | 32213.76 | 0.48 | 100.0% | 32_27_0 | F1 | P4 |
| 26 | 32239.34 | 32241.81 | 2.47 | 48.4% | 32_27_0 | S | P4 |
| 27 | 32359.60 | 32359.90 | 0.30 | 7.9% | 32_27_1 | F1 | P4F1 |
| 28 | 32386.62 | 32387.95 | 1.33 | 6.9% | 32_27_1 | S | P4F1 |
| 29 | 32422.65 | 32416.95 | 5.70 | 28.1% | 32_28_0 | F1 | ★ |
| 30 | 32444.74 | 32445.00 | 0.26 | 8.5% | 32_28_0 | S | ★ |
| 31 | 32506.21 | 32506.04 | 0.17 | 4.3% | 32_27_2 | F1 | P4F2 |
| 32 | 32533.35 | 32534.09 | 0.74 | 2.4% | 32_27_2 | S | P4F2 |
| 33 | 32578.91 | 32579.09 | 0.18 | 83.8% | 33_28_0 | F1 | P5 |
| 34 | 32605.09 | 32607.14 | 2.05 | 44.5% | 33_28_0 | S | P5 |
| 35 | 32725.24 | 32725.23 | 0.01 | 9.6% | 33_28_1 | F1 | P5F1 |
| 36 | 32752.22 | 32753.28 | 1.06 | 7.2% | 33_28_1 | S | P5F1 |
| 37 | 32788.38 | 32782.28 | 6.10 | 25.0% | 33_29_0 | F1 | ★ |
| 38 | 32810.18 | 32810.33 | 0.15 | 6.6% | 33_29_0 | S | ★ |
| 39 | 32871.62 | 32871.37 | 0.24 | 4.3% | 33_28_2 | F1 | P5F2 |
| 40 | 32898.77 | 32899.42 | 0.66 | 2.3% | 33_28_2 | S | P5F2 |
| 41 | 32944.77 | 32944.42 | 0.34 | 29.8% | 34_29_0 | F1 | P6 |
| 42 | 32971.17 | 32972.47 | 1.31 | 16.9% | 34_29_0 | S | P6 |
| 43 | 33090.84 | 33090.56 | 0.28 | 6.7% | 34_29_1 | F1 | P6F1 |
| 44 | 33118.01 | 33118.61 | 0.60 | 3.8% | 34_29_1 | S | P6F1 |
| 45 | 33153.82 | 33147.62 | 6.20 | 8.1% | 34_30_0 | F1 | ★ |
| 46 | 33175.75 | 33175.67 | 0.08 | 2.0% | 34_30_0 | S | ★ |
| 47 | 33237.16 | 33236.71 | 0.45 | 2.3% | 34_29_2 | F1 | P6F2 |

| | | | | | | | |
|---|---|---|---|---|---|---|---|
| 48 | 33265.07 | 33264.76 | 0.32 | 1.3% | 34_29_2 | S | P6F2 |
| 49 | 33309.81 | 33309.76 | 0.05 | 17.5% | 35_30_0 | F1 | P7 |
| 50 | 33336.84 | 33337.81 | 0.97 | 10.9% | 35_30_0 | S | P7 |
| 51 | 33455.79 | 33455.90 | 0.10 | 4.9% | 35_30_1 | F1 | P7F1 |
| 52 | 33483.22 | 33483.95 | 0.73 | 2.8% | 35_30_1 | S | P7F1 |
| 53 | 33519.26 | 33512.95 | 6.31 | 4.7% | 35_31_0 | F1 | ★ |
| 54 | 33541.08 | 33541.00 | 0.08 | 1.3% | 35_31_0 | S | ★ |
| 55 | 33602.17 | 33602.04 | 0.13 | 2.0% | 35_30_2 | F1 | P7F2 |
| 56 | 33630.21 | 33630.09 | 0.12 | 1.1% | 35_30_2 | S | P7F2 |
| 57 | 33675.22 | 33675.09 | 0.13 | 11.3% | 36_31_0 | F1 | P8 |
| 58 | 33702.32 | 33703.14 | 0.82 | 6.9% | 36_31_0 | S | P8 |
| 59 | 33821.33 | 33821.23 | 0.10 | 3.7% | 36_31_1 | F1 | P8F1 |
| 60 | 33849.31 | 33849.28 | 0.03 | 2.2% | 36_31_1 | S | P8F1 |
| 61 | 33884.18 | 33878.28 | 5.90 | 2.8% | 36_32_0 | F1 | ★ |
| 62 | 33906.14 | 33906.33 | 0.19 | 1% | 36_32_0 | S | ★ |
| 63 | 33967.83 | 33967.37 | 0.46 | 2% | 36_31_2 | F1 | P8F2 |
| 64 | 33995.79 | 33995.42 | 0.36 | 2% | 36_31_2 | S | P8F2 |
| 65 | 34040.35 | 34040.42 | 0.08 | 7% | 37_32_0 | F1 | P9 |
| 66 | 34067.84 | 34068.47 | 0.63 | 4% | 37_32_0 | S | P9 |
| 67 | 34186.97 | 34186.56 | 0.41 | 2% | 37_32_1 | F1 | P9F1 |
| 68 | 34214.16 | 34214.61 | 0.45 | 2% | 37_32_1 | S | P9F1 |
| 69 | 34250.01 | 34243.62 | 6.39 | 2% | 37_33_0 | F1 | ★ |
| 70 | 34271.31 | 34271.67 | 0.36 | 0% | 37_33_0 | S | ★ |
| 71 | 34333.36 | 34332.71 | 0.66 | 1% | 37_32_2 | F1 | P9F2 |
| 72 | 34358.35 | 34360.76 | 2.41 | 1% | 37_32_2 | S | P9F2 |

| | | | | | | | |
|---|---|---|---|---|---|---|---|
| 73 | 34402.09 | 34405.76 | 3.67 | 4% | 38_33_0 | F1 | P10 |
| 74 | 34432.66 | 34433.81 | 1.15 | 3% | 38_33_0 | S | P10 |
| 75 | 34547.74 | 34551.90 | 4.16 | 1% | 38_33_1 | F1 | P10F1 |
| 76 | 34577.75 | 34579.95 | 2.20 | 9% | 38_33_1 | S | P10F1 |

a. The experimental masses were calculated based on the average MW of glycan compositions at three charge states (+8 to +10).
b. The theoretical masses were calculated from the mass of peptide backbone of AGP (F1 variant: 21539 Da, S variant: 21567 Da) and the average masses of monosaccharides (Hex: 162.14 Da, HexNAc: 203.19 Da, Fuc: 146.14 Da).
c. Glycan compositions are expressed as the number of Hex, HexNAc, and Fuc residues.
d. Red stars identify compositions consistent with the presence of terminal GlcNAc residues.

**Table S5.** Annotation of AG-AD-asialo-AGP glycan compositions.

| Num. | Experimental Mass (Da)[a] | Theoretical Mass (Da)[b] | Mass Difference (Da) | Relative Abundance (%) | Composition [c] | Genetic Variant | Annotation |
|---|---|---|---|---|---|---|---|
| 1 | 28644.88 | 28644.60 | 0.29 | 3.4% | 15_23_0 | F1 | P’0 |
| 2 | 28667.14 | 28672.65 | 5.51 | 2.2% | 15_23_0 | S | P’0 |
| 3 | 28846.98 | 28847.79 | 0.81 | 22.5% | 15_24_0 | F1 | P’1 |
| 4 | 28869.36 | 28875.84 | 6.48 | 6.8% | 15_24_0 | S | P’1 |
| 5 | 28942.69 | 28936.88 | 5.81 | 2.0% | 15_23_2 | F1 | P’0F2 |
| 6 | 28967.13 | 28964.93 | 2.20 | 1.7% | 15_23_2 | S | P’0F2 |
| 7 | 28988.97 | 28993.93 | 4.96 | 1.7% | 15_24_1 | F1 | P’1F1 |
| 8 | 29049.55 | 29050.98 | 1.43 | 52.3% | 15_25_0 | F1 | P’2 |
| 9 | 29073.29 | 29079.03 | 5.74 | 20.4% | 15_25_0 | S | P’2 |
| 10 | 29113.07 | 29115.02 | 1.95 | 4.1% | 17_23_1 | F1 | |
| 11 | 29144.77 | 29140.07 | 4.69 | 3.6% | 15_24_2 | F1 | P’1F2 |
| 12 | 29168.18 | 29168.12 | 0.05 | 2.4% | 15_24_2 | S | P’1F2 |
| 13 | 29194.62 | 29197.12 | 2.50 | 2.6% | 15_25_1 | F1 | P’2F1 |
| 14 | 29252.36 | 29254.17 | 1.81 | 79.3% | 15_26_0 | F1 | P’3 |
| 15 | 29277.20 | 29282.22 | 5.02 | 35.5% | 15_26_0 | S | P’3 |
| 16 | 29317.34 | 29318.21 | 0.87 | 6.8% | 17_24_1 | F1 | |
| 17 | 29350.47 | 29346.26 | 4.21 | 5.5% | 17_24_1 | S | |
| 18 | 29374.74 | 29371.31 | 3.42 | 3.8% | 15_25_2 | S | P’2F2 |
| 19 | 29397.16 | 29400.32 | 3.15 | 3.4% | 15_26_1 | F1 | P’3F1 |
| 20 | 29415.42 | 29416.32 | 0.90 | 9.8% | 16_26_0 | F1 | |
| 21 | 29455.88 | 29457.37 | 1.49 | 100.0% | 15_27_0 | F1 | P’4 |
| 22 | 29480.90 | 29485.42 | 4.52 | 49.4% | 15_27_0 | S | P’4 |
| 23 | 29520.76 | 29521.40 | 0.65 | 9.7% | 17_25_1 | F1 | |

| | | | | | | | |
|---|---|---|---|---|---|---|---|
| 24 | 29550.12 | 29546.46 | 3.66 | 6.5% | 15_26_2 | F1 | P’3F2 |
| 25 | 29577.72 | 29574.51 | 3.21 | 5.1% | 15_26_2 | S | P’3F2 |
| 26 | 29598.90 | 29603.51 | 4.61 | 3.8% | 15_27_1 | F1 | P’4F1 |
| 27 | 29618.96 | 29619.51 | 0.55 | 17.5% | 16_27_0 | F1 | |
| 28 | 29639.61 | 29647.56 | 7.95 | 21.7% | 16_27_0 | S | |
| 29 | 29659.17 | 29660.56 | 1.39 | 67.5% | 15_28_0 | F1 | P’5 |
| 30 | 29685.20 | 29688.61 | 3.41 | 41.5% | 15_28_0 | S | P’5 |
| 31 | 29707.17 | 29708.60 | 1.43 | 5.7% | 16_26_2 | F1 | |
| 32 | 29724.05 | 29724.60 | 0.55 | 10.0% | 17_26_1 | F1 | |
| 33 | 29753.70 | 29752.65 | 1.05 | 6.1% | 17_26_1 | S | |
| 34 | 29760.95 | 29765.65 | 4.70 | 5.0% | 16_27_1 | F1 | |
| 35 | 29781.55 | 29781.65 | 0.10 | 5.1% | 17_27_0 | F1 | |
| 36 | 29797.96 | 29793.70 | 4.26 | 5.5% | 16_27_1 | S | |
| 37 | 29821.55 | 29822.70 | 1.15 | 24.1% | 16_28_0 | F1 | |
| 38 | 29845.69 | 29850.75 | 5.06 | 10.2% | 16_28_0 | S | |
| 39 | 29869.54 | 29863.75 | 5.79 | 24.1% | 15_29_0 | F1 | P’6 |
| 40 | 29890.11 | 29891.80 | 1.69 | 10.9% | 15_29_0 | S | P’6 |
| 41 | 29908.77 | 29911.79 | 3.02 | 4.9% | 16_27_2 | F1 | |
| 42 | 29927.78 | 29927.79 | 0.01 | 6.3% | 17_27_1 | F1 | |
| 43 | 29951.54 | 29952.84 | 1.31 | 4.0% | 15_28_2 | F1 | P’5F2 |
| 44 | 29967.23 | 29968.84 | 1.61 | 5.6% | 16_28_1 | F1 | |
| 45 | 29985.01 | 29984.84 | 0.17 | 4.7% | 17_28_0 | F1 | |
| 46 | 30000.57 | 29996.89 | 3.68 | 5.5% | 16_28_1 | S | |
| 47 | 30024.96 | 30025.89 | 0.93 | 20.4% | 16_29_0 | F1 | |
| 48 | 30050.36 | 30053.94 | 3.58 | 11.4% | 16_29_0 | S | |

| | | | | | | | |
|---|---|---|---|---|---|---|---|
| 49 | 30072.48 | 30066.94 | 5.53 | 4.6% | 15_30_0 | F1 | P’7 |
| 50 | 30090.40 | 30094.99 | 4.59 | 3.8% | 15_30_0 | S | P’7 |
| 51 | 30130.53 | 30130.98 | 0.45 | 4.3% | 17_28_1 | F1 | |
| 52 | 30187.66 | 30188.03 | 0.37 | 7.9% | 17_29_0 | F1 | |
| 53 | 30209.83 | 30213.09 | 3.26 | 4.5% | 15_30_1 | F1 | P’7F1 |
| 54 | 30230.21 | 30229.09 | 1.12 | 8.3% | 16_30_0 | F1 | |
| 55 | 30254.68 | 30257.14 | 2.46 | 6.0% | 16_30_0 | S | |
| 56 | 30275.39 | 30270.14 | 5.25 | 2.7% | 15_31_0 | F1 | P’8 |
| 57 | 30355.13 | 30359.23 | 4.10 | 3.3% | 15_30_2 | F1 | P’7F2 |
| 58 | 30390.15 | 30391.23 | 1.08 | 10.2% | 17_30_0 | F1 | |
| 59 | 30415.56 | 30419.28 | 3.71 | 6% | 17_30_0 | S | |
| 60 | 30478.11 | 30473.33 | 4.78 | 3% | 15_32_0 | F1 | P’9 |
| 61 | 30593.96 | 30594.42 | 0.45 | 6% | 17_31_0 | F1 | |
| 62 | 30619.51 | 30619.47 | 0.04 | 4% | 15_32_1 | F1 | P’9F1 |
| 63 | 30761.16 | 30765.61 | 4.45 | 5% | 15_32_2 | F1 | P’9F2 |

a. The experimental masses were calculated based on the average MW of glycan compositions at three charge states (+8 to +10).
b. The theoretical masses were calculated from the mass of peptide backbone of AGP (F1 variant: 21539 Da, S variant: 21567 Da) and the average masses of monosaccharides (Hex: 162.14 Da, HexNAc: 203.19 Da, Fuc: 146.14 Da).
c. Glycan compositions are expressed as the number of Hex, HexNAc, and Fuc residues.

**Table S6.** Annotation of TG-AG-AD-asialo-AGP glycan compositions.

| Num. | Experimental Mass (Da)[a] | Theoretical Mass (Da)[b] | Mass Difference (Da) | Relative Abundance (%) | Composition [c] | Genetic Variant | Annotation[d] |
|---|---|---|---|---|---|---|---|
| 1 | 26002.49 | 26003.09 | 0.60 | 100% | 15_10_0 | F1 | P’’1 |
| 2 | 26027.30 | 26031.14 | 3.85 | 43% | 15_10_0 | S | P’’1 |
| 3 | 26148.43 | 26149.24 | 0.81 | 6% | 15_10_1 | F1 | P’’1F1 |
| 4 | 26212.43 | 26206.29 | 6.14 | 27% | 15_11_0 | F1 | P’’2★ |
| 5 | 26234.45 | 26234.34 | 0.11 | 10% | 15_11_0 | S | P’’2★ |
| 6 | 26297.12 | 26295.38 | 1.74 | 4% | 15_10_2 | F1 | P’’1F2 |
| 7 | 26327.41 | 26323.43 | 3.98 | 3% | 15_10_2 | S | P’’1F2 |
| 8 | 26405.34 | 26409.48 | 4.14 | 3% | 15_12_0 | F1 | P’’3★ |

a. The experimental masses were calculated based on the average MW of glycan compositions at three charge states (+7 to +9).
b. The theoretical masses were calculated from the mass of peptide backbone of AGP (F1 variant: 21539 Da, S variant: 21567 Da) and the average masses of monosaccharides (Hex: 162.14 Da, HexNAc: 203.19 Da, Fuc: 146.14 Da).
c. Glycan compositions are expressed as the number of Hex, HexNAc, and Fuc residues.
d. Red stars identify compositions consistent with the presence of terminal GlcNAc residues.

**Table S7.** Annotation of CG-TG-AG-AD-asialo-AGP glycan compositions.

| Num. | Experimental Mass (Da)[a] | Theoretical Mass (Da)[b] | Mass Difference (Da) | Relative Abundance (%) | Composition[c] | Genetic Variant | Annotation[d] |
|---|---|---|---|---|---|---|---|
| 1 | 22556.31 | 22555.06 | 1.24 | 1.3% | 0_5_0 | F1 | P'''1 |
| 2 | 22578.36 | 22583.06 | 4.70 | 0.4% | 0_5_0 | S | P'''1 |
| 3 | 23243.19 | 23244.68 | 1.49 | 7.2% | 3_6_0 | F1 | P'''2 |
| 4 | 23271.35 | 23272.68 | 1.33 | 2.3% | 3_6_0 | S | P'''2 |
| 5 | 23453.29 | 23447.87 | 5.42 | 4.7% | 3_7_0 | F1 | P'''3★ |
| 6 | 23473.62 | 23475.87 | 2.25 | 0.7% | 3_7_0 | S | P'''3★ |
| 7 | 23932.70 | 23934.29 | 1.59 | 34.8% | 6_7_0 | F1 | P'''4 |
| 8 | 23960.90 | 23962.29 | 1.39 | 11.5% | 6_7_0 | S | P'''4 |
| 9 | 24142.91 | 24137.48 | 5.43 | 14.4% | 6_8_0 | F1 | P'''5★ |
| 10 | 24163.65 | 24165.48 | 1.83 | 1.9% | 6_8_0 | S | P'''5★ |
| 11 | 24622.51 | 24623.91 | 1.40 | 100.0% | 9_8_0 | F1 | P'''6 |
| 12 | 24650.41 | 24651.91 | 1.50 | 36.1% | 9_8_0 | S | P'''6 |
| 13 | 24832.61 | 24827.10 | 5.52 | 11.3% | 9_9_0 | F1 | P'''7★ |
| 14 | 24854.39 | 24855.10 | 0.71 | 1.5% | 9_9_0 | S | P'''7★ |
| 15 | 25163.93 | 25176.43 | 12.50 | 5.0% | 9_10_1 | F1 | P'''8★ |
| 16 | 25191.78 | 25204.43 | 12.65 | 2.1% | 9_10_1 | S | P'''8★ |
| 17 | 25312.03 | 25313.52 | 1.49 | 12.4% | 12_9_0 | F1 | P'''9 |
| 18 | 25339.55 | 25341.52 | 1.97 | 4.3% | 12_9_0 | S | P'''9 |
| 19 | 25607.83 | 25605.80 | 2.02 | 5.8% | 12_9_2 | F1 | P'''9F2 |
| 20 | 25635.83 | 25633.80 | 2.03 | 2.0% | 12_9_2 | S | P'''9F2 |
| 21 | 26001.91 | 26003.13 | 1.23 | 1.6% | 15_10_0 | F1 | P'''11 |
| 22 | 26031.51 | 26031.13 | 0.38 | 0.8% | 15_10_0 | S | P'''11 |

a. The experimental masses were calculated based on the average MW of glycan compositions at three charge states (+7 to +9).
b. The theoretical masses were calculated from the mass of peptide backbone of AGP (F1 variant: 21539 Da, S variant: 21567 Da) and the average masses of monosaccharides (Hex: 162.14 Da, HexNAc: 203.19 Da, Fuc: 146.14 Da).
c. Glycan compositions are expressed as the number of Hex, HexNAc, and Fuc residues.
d. Red stars identify compositions consistent with the presence of terminal GlcNAc residues.

**Table S8.** Affinities ($K_d$, μM) and thermodynamic parameters (-Δ$H$ and $T$Δ$S$, kJ mol$^{-1}$) for asialo-AGP (3 μM) F1 variant compositions binding to warfarin (0.25 μM – 8 μM). Measurements were performed on ammonium acetate solutions (200 mM, pH 7.6) in 1% DMSO. [a,b]

| Annotation | Variant | Composition[c] | $K_d$ (μM) | 1/$K_d$ (μM$^{-1}$) | -Δ$H$ (kJ mol$^{-1}$) | $T$Δ$S$ (kJ mol$^{-1}$)[d] |
|---|---|---|---|---|---|---|
| P1F0 | F1 | 29_24_0 | 0.7 ± 0.1 | 1.4 ± 0.1 | 35.5 ± 1.0 | -1.6 ± 2.4 |
| P2F0 | F1 | 30_25_0 | 1.0 ± 0.2 | 1.1 ± 0.2 | 37.9 ± 0.8 | -3.9 ± 1.9 |
| P3F0 | F1 | 31_26_0 | 1.0 ± 0.2 | 1.0 ± 0.2 | 41.6 ± 1.0 | -7.5 ± 2.5 |
| P4F0 | F1 | 32_27_0 | 1.0 ± 0.2 | 1.1 ± 0.2 | 45.5 ± 1.3 | -12.5 ± 3.1 |
| P5F0 | F1 | 33_28_0 | 1.0 ± 0.2 | 1.0 ± 0.2 | 44.3 ± 1.1 | -11.0 ± 2.7 |
| P6F0 | F1 | 34_29_0 | 1.0 ± 0.2 | 1.0 ± 0.1 | 41.0 ± 0.6 | -7.2 ± 1.5 |
| P7F0 | F1 | 35_30_0 | 1.0 ± 0.2 | 1.0 ± 0.2 | 38.1 ± 0.7 | -4.5 ± 1.8 |
| P8F0 | F1 | 36_31_0 | 0.7 ± 0.1 | 1.3 ± 0.1 | 35.5 ± 0.6 | -1.2 ± 1.4 |
| P9F0 | F1 | 37_32_0 | 0.9 ± 0.1 | 1.1 ± 0.2 | 32.7 ± 0.7 | 1.6 ± 1.6 |

| Annotation | Variant | Composition[c] | $K_d$ (μM) | 1/$K_d$ (μM$^{-1}$) | -Δ$H$ (kJ mol$^{-1}$) | $T$Δ$S$ (kJ mol$^{-1}$)[d] |
|---|---|---|---|---|---|---|
| P1F1 | F1 | 29_24_1 | 0.6 ± 0.1 | 1.7 ± 0.2 | 30.5 ± 0.5 | 3.1 ± 1.4 |
| P2F1 | F1 | 30_25_1 | 0.8 ± 0.1 | 1.2 ± 0.2 | 35.9 ± 0.9 | -1.9 ± 2.3 |
| P3F1 | F1 | 31_26_1 | 0.9 ± 0.2 | 1.1 ± 0.2 | 36.6 ± 0.7 | -2.2 ± 1.7 |
| P4F1 | F1 | 32_27_1 | 1.0 ± 0.2 | 1.0 ± 0.2 | 40.0 ± 0.8 | -5.8 ± 2.1 |
| P5F1 | F1 | 33_28_1 | 0.9 ± 0.2 | 1.1 ± 0.2 | 35.6 ± 0.7 | -1.6 ± 1.8 |
| P6F1 | F1 | 34_29_1 | 0.8 ± 0.1 | 1.2 ± 0.1 | 28.6 ± 0.9 | 5.5 ± 2.3 |
| P7F1 | F1 | 35_30_1 | 0.8 ± 0.1 | 1.2 ± 0.1 | 28.1 ± 0.6 | 5.9 ± 1.6 |
| P8F1 | F1 | 36_31_1 | 0.6 ± 0.1 | 1.5 ± 0.1 | 24.4 ± 0.6 | 10.4 ± 1.6 |
| P9F1 | F1 | 37_32_1 | 0.5 ± 0.1 | 2.2 ± 0.5 | 24.9 ± 0.3 | 11.0 ± 0.7 |

| **Annotation** | **Variant** | **Composition**[c] | $K_d$ (μM) | $1/K_d$ ($\mu M^{-1}$) | $-\Delta H$ ($kJ\ mol^{-1}$) | $T\Delta S$ ($kJ\ mol^{-1}$)[d] |
|---|---|---|---|---|---|---|
| P1F2 | F1 | 29_24_2 | 1.9± 0.2 | 0.5 ± 0.1 | 49.9 ± 1.5 | -18.3 ± 3.7 |
| P2F2 | F1 | 30_25_2 | 2.0 ± 0.2 | 0.5 ± 0.1 | 60.6 ± 1.2 | -30.0 ± 3.1 |
| P3F2 | F1 | 31_26_2 | 2.2 ± 0.1 | 0.5 ± 0.1 | 74.3 ± 1.0 | -44.3 ± 2.4 |
| P4F2 | F1 | 32_27_2 | 2.1 ± 0.2 | 0.5 ± 0.1 | 77.6 ± 1.4 | -47.0 ± 3.6 |
| P5F2 | F1 | 33_28_2 | 1.7 ± 0.2 | 0.6 ± 0.1 | 56.6 ± 1.0 | -25.6 ± 2.5 |
| P6F2 | F1 | 34_29_2 | 1.2± 0.2 | 0.8 ± 0.1 | 42.4 ± 0.8 | -10.7 ± 1.9 |
| P7F2 | F1 | 35_30_2 | 1.1 ± 0.1 | 0.9 ± 0.1 | 41.0 ± 0.8 | -9.0 ± 1.9 |
| P8F2 | F1 | 36_31_2 | 0.9 ± 0.1 | 1.1 ± 0.2 | 35.8 ± 0.7 | -3.3 ± 1.8 |
| P9F2 | F1 | 37_32_2 | 0.9 ± 0.2 | 1.1 ± 0.3 | 27.4 ± 0.7 | 6.7 ± 1.8 |

| **Annotation** | **Variant** | **Composition**[c] | $K_d$ (μM) | $1/K_d$ ($\mu M^{-1}$) | $-\Delta H$ ($kJ\ mol^{-1}$) | $T\Delta S$ ($kJ\ mol^{-1}$)[d] |
|---|---|---|---|---|---|---|
| P1F3 | F1 | 29_24_3 | 0.3 ± 0.1 | 2.9 ± 1.0 | ND | ND |
| P2F3 | F1 | 30_25_3 | 0.6 ± 0.2 | 1.7 ± 0.6 | 36.8 ± 0.6 | -2.6 ± 1.6 |
| P3F3 | F1 | 31_26_3 | 1.2 ± 0.2 | 0.8 ± 0.2 | 40.6 ± 0.5 | -6.7 ± 1.3 |
| P4F3 | F1 | 32_27_3 | 1.2 ± 0.2 | 0.8 ± 0.2 | 52.2 ± 1.1 | -19.1 ± 2.7 |
| P5F3 | F1 | 33_28_3 | 1.3 ± 0.1 | 0.8 ± 0.1 | 45.8 ± 0.9 | -13.9 ± 2.1 |
| P6F3 | F1 | 34_29_3 | 0.8 ± 0.2 | 1.2 ± 0.2 | 40.2 ± 0.9 | -8.0 ± 2.1 |
| P7F3 | F1 | 35_30_3 | 0.8 ± 0.2 | 1.2 ± 0.2 | 32.9 ± 0.7 | -0.6 ± 1.8 |
| P8F3 | F1 | 36_31_3 | 0.9 ± 0.2 | 1.2 ± 0.3 | 29.9 ± 0.6 | 3.2 ± 1.4 |
| P9F3 | F1 | 37_32_3 | 0.5 ± 0.2 | 1.9 ± 0.6 | 20.5 ± 0.6 | 14.7 ± 1.4 |

a. Errors correspond to one standard deviation.
b. ND = Not determined.
c. The compositions of glycan compositions corresponded to the number of Hexose (Hex), N-acetylhexosamine (HexNAc), Fucose (Fuc).
d. Temperature ($T$) is 298.15 K.

**Table S9.** Affinities ($K_d$, μM) and thermodynamic parameters (-Δ$H$ and $T$Δ$S$, kJ $mol^{-1}$) for AD-asialo-AGP (3 μM) F1 variant compositions binding to warfarin (0.25 μM – 8 μM). Measurements were performed on ammonium acetate solutions (200 mM, pH 7.6) in 1% DMSO. [a,b]

| Annotation | Variant | Composition[c] | $K_d$ (μM) | $1/K_d$ ($\mu M^{-1}$) | -Δ$H$ (kJ $mol^{-1}$) | $T$Δ$S$ (kJ $mol^{-1}$)[d] |
|---|---|---|---|---|---|---|
| P1F0 | F1 | 29_24_0 | 0.7 ± 0.1 | 1.5 ± 0.2 | 36.0 ± 1.2 | -2.0 ± 1.2 |
| P2F0 | F1 | 30_25_0 | 0.7 ± 0.1 | 1.4 ± 0.2 | 38.2 ± 1.3 | -4.0 ± 1.3 |
| P3F0 | F1 | 31_26_0 | 0.7 ± 0.1 | 1.4 ± 0.2 | 40.8 ± 1.3 | -6.2 ± 1.2 |
| P4F0 | F1 | 32_27_0 | 0.7 ± 0.1 | 1.4 ± 0.2 | 44.6 ± 0.8 | -9.8 ± 0.8 |
| P5F0 | F1 | 33_28_0 | 0.7 ± 0.1 | 1.4 ± 0.2 | 42.7 ± 1.0 | -7.6 ± 1.0 |
| P6F0 | F1 | 34_29_0 | 0.7 ± 0.2 | 1.4 ± 0.3 | 38.7 ± 1.0 | -4.4 ± 1.0 |
| P7F0 | F1 | 35_30_0 | 0.7 ± 0.1 | 1.5 ± 0.3 | 38.1 ± 1.5 | -3.5 ± 1.5 |
| P8F0 | F1 | 36_31_0 | 0.6 ± 0.1 | 1.7 ± 0.3 | 35.0 ± 1.8 | -0.5 ± 1.7 |
| P9F0 | F1 | 37_32_0 | 0.6 ± 0.1 | 1.7 ± 0.3 | 31.9 ± 1.2 | 2.0 ± 1.2 |

| Annotation | Variant | Composition[c] | $K_d$ (μM) | $1/K_d$ ($\mu M^{-1}$) | -Δ$H$ (kJ $mol^{-1}$) | $T$Δ$S$ (kJ $mol^{-1}$)[d] |
|---|---|---|---|---|---|---|
| P1F1 | F1 | 29_24_1 | ND | ND | ND | ND |
| P2F1 | F1 | 30_25_1 | 0.3 ± 0.1 | 3.4 ± 0.5 | 16.2 ± 0.9 | 19.1 ± 0.9 |
| P3F1 | F1 | 31_26_1 | 0.4 ± 0.1 | 2.4 ± 0.3 | 19.1 ± 0.6 | 16.6 ± 0.6 |
| P4F1 | F1 | 32_27_1 | 0.5 ± 0.1 | 1.9 ± 0.2 | 19.5 ± 0.6 | 16.6 ± 0.6 |
| P5F1 | F1 | 33_28_1 | 0.5 ± 0.1 | 2.0 ± 0.2 | 18.2 ± 0.7 | 17.5 ± 0.7 |
| P6F1 | F1 | 34_29_1 | 0.5 ± 0.1 | 2.2 ± 0.2 | 15.9 ± 0.8 | 19.3 ± 0.8 |
| P7F1 | F1 | 35_30_1 | 0.5 ± 0.1 | 2.1 ± 0.3 | 15.6 ± 0.6 | 20.9 ± 0.6 |
| P8F1 | F1 | 36_31_1 | 0.5 ± 0.1 | 2.1 ± 0.3 | 16.1 ± 0.7 | 20.5 ± 0.6 |
| P9F1 | F1 | 37_32_1 | 0.4 ± 0.1 | 2.5 ± 0.4 | 16.4 ± 0.6 | 18.8± 0.6 |

a. Errors correspond to one standard deviation.
b. ND = Not determined.
c. The compositions of glycan compositions corresponded to the number of Hexose (Hex), N-acetylhexosamine (HexNAc), Fucose (Fuc).
d. Temperature ($T$) is 298.15 K.

**Table S10.** Affinities ($K_d$, μM) and thermodynamic parameters ($-\Delta H$ and $T\Delta S$, kJ $mol^{-1}$) for Asialo-AGP (3 μM) S variant compositions binding to warfarin (0.25 μM – 8 μM). Measurements were performed on ammonium acetate solutions (200 mM, pH 7.6) in 1% DMSO. [a,b]

| Annotation | Variant | Composition[c] | $K_d$ (μM) | $1/K_d$ ($μM^{-1}$) | $-\Delta H$ (kJ $mol^{-1}$) | $T\Delta S$ (kJ $mol^{-1}$)[d] |
|---|---|---|---|---|---|---|
| P2F0 | S | 30_25_0 | 0.4 ± 0.1 | 2.3 ± 0.6 | 31.1 ± 0.6 | 6.7 ± 0.6 |
| P3F0 | S | 31_26_0 | 0.6 ± 0.1 | 1.8 ± 0.3 | 42.5 ± 0.9 | -8.3 ± 0.9 |
| P4F0 | S | 32_27_0 | 0.7 ± 0.1 | 1.4 ± 0.2 | 46.8 ± 1.6 | -13.1 ± 1.6 |
| P5F0 | S | 33_28_0 | 0.8 ± 0.1 | 1.3 ± 0.2 | 53.4 ± 1.1 | -19.8 ± 1.1 |
| P6F0 | S | 34_29_0 | 0.7 ± 0.1 | 1.4 ± 0.2 | 28.4 ± 0.9 | 5.9 ± 0.9 |
| P7F0 | S | 35_30_0 | 0.7 ± 0.1 | 1.5 ± 0.2 | 28.1 ± 0.8 | 6.3 ± 0.8 |
| P8F0 | S | 36_31_0 | 0.6 ± 0.1 | 1.6 ± 0.2 | 28.6 ± 0.8 | 6.4 ± 0.8 |
| P9F0 | S | 37_32_0 | 0.6 ± 0.1 | 1.7 ± 0.3 | 22.0 ± 0.6 | 13.2 ± 0.6 |

| Annotation | Variant | Composition[c] | $K_d$ (μM) | $1/K_d$ ($μM^{-1}$) | $-\Delta H$ (kJ $mol^{-1}$) | $T\Delta S$ (kJ $mol^{-1}$)[d] |
|---|---|---|---|---|---|---|
| P2F1 | S | 30_25_1 | 0.4 ± 0.1 | 2.4 ± 0.5 | 33.6 ± 0.7 | 4.1 ± 1.0 |
| P3F1 | S | 31_26_1 | 0.6 ± 0.1 | 1.6 ± 0.3 | 37.7 ± 1.0 | -3.9 ± 1.0 |
| P4F1 | S | 32_27_1 | 0.8 ± 0.1 | 1.3 ± 0.2 | 39.0 ± 1.5 | -5.8 ± 1.5 |
| P5F1 | S | 33_28_1 | 0.8 ± 0.1 | 1.2 ± 0.2 | 45.6 ± 1.0 | -12.4 ± 0.9 |
| P6F1 | S | 34_29_1 | 0.7 ± 0.1 | 1.4 ± 0.2 | 30.7 ± 0.5 | 3.7 ± 0.5 |
| P7F1 | S | 35_30_1 | 0.6 ± 0.1 | 1.7 ± 0.3 | 18.4 ± 0.5 | 16.2 ± 0.5 |
| P8F1 | S | 36_31_1 | 0.7 ± 0.1 | 1.4 ± 0.2 | 23.0 ± 0.6 | 10.6 ± 0.6 |
| P9F1 | S | 37_32_1 | 0.6 ± 0.1 | 1.7 ± 0.3 | 21.9 ± 0.5 | 13.1 ± 0.5 |

| Annotation | Variant | Composition[c] | $K_d$ (μM) | $1/K_d$ ($\mu M^{-1}$) | $-\Delta H$ ($kJ\ mol^{-1}$) | $T\Delta S$ ($kJ\ mol^{-1}$)[d] |
|---|---|---|---|---|---|---|
| P2F2 | S | 30_25_2 | 0.2 ± 0.1 | 4.1 ± 0.5 | ND | ND |
| P3F2 | S | 31_26_2 | 0.3 ± 0.1 | 3.3 ± 0.8 | 9.1 ± 0.3 | 26.6 ± 0.3 |
| P4F2 | S | 32_27_2 | 0.3 ± 0.1 | 3.4 ± 0.6 | 15.9 ± 0.5 | 19.3 ± 0.5 |
| P5F2 | S | 33_28_2 | 0.6 ± 0.1 | 1.7 ± 0.4 | 14.6 ± 0.3 | 20.4 ± 0.3 |
| P6F2 | S | 34_29_2 | 0.6 ± 0.1 | 1.7 ± 0.3 | 12.3 ± 0.2 | 22.7 ± 0.2 |
| P7F2 | S | 35_30_2 | 0.5 ± 0.1 | 1.9 ± 0.3 | 12.2 ± 0.4 | 23.1 ± 0.4 |
| P8F2 | S | 36_31_2 | 0.6 ± 0.1 | 1.6 ± 0.4 | 8.9 ± 0.4 | 25.5 ± 0.4 |
| P9F2 | S | 37_32_2 | 0.4 ± 0.1 | 2.3 ± 0.4 | 1.8 ± 1.1 | 34.3 ± 1.1 |

| Annotation | Variant | Composition[c] | $K_d$ (μM) | $1/K_d$ ($\mu M^{-1}$) | $-\Delta H$ ($kJ\ mol^{-1}$) | $T\Delta S$ ($kJ\ mol^{-1}$)[d] |
|---|---|---|---|---|---|---|
| P2F3 | S | 30_25_3 | 0.7 ± 0.1 | 1.4 ± 0.2 | 16.9 ± 0.5 | 17.0 ± 0.5 |
| P3F3 | S | 31_26_3 | 0.4 ± 0.1 | 2.4 ± 0.8 | 14.9 ± 0.3 | 18.7 ± 0.3 |
| P4F3 | S | 32_27_3 | 0.5 ± 0.1 | 1.9 ± 0.3 | 17.3 ± 0.5 | 18.5 ± 0.5 |
| P5F3 | S | 33_28_3 | 0.6 ± 0.1 | 1.6 ± 0.3 | 16.2 ± 0.5 | 19.1 ± 0.6 |
| P6F3 | S | 34_29_3 | 0.4 ± 0.1 | 2.4 ± 0.6 | 11.9 ± 0.5 | 23.1 ± 0.5 |
| P7F3 | S | 35_30_3 | 0.4 ± 0.1 | 2.2 ± 0.5 | 9.9 ± 0.3 | 25.7 ± 0.3 |
| P8F3 | S | 36_31_3 | 0.5 ± 0.1 | 1.8 ± 0.4 | 7.2 ± 0.2 | 27.8 ± 0.2 |
| P9F3 | S | 37_32_3 | ND | ND | ND | ND |

a. Errors correspond to one standard deviation.
b. ND = Not determined.
c. The compositions of glycan compositions corresponded to the number of Hexose (Hex), N-acetylhexosamine (HexNAc), Fucose (Fuc).
d. Temperature ($T$) is 298.15 K.

**Table S11.** Affinities ($K_d$, μM) and thermodynamic parameters ($-\Delta H$ and $T\Delta S$, kJ $mol^{-1}$) for AD-asialo-AGP (3 μM) S variant compositions binding to warfarin (0.25 μM – 8 μM). Measurements were performed on ammonium acetate solutions (200 mM, pH 7.6) in 1% DMSO. [a,b]

| Annotation | Variant | Composition[c] | $K_d$ (μM) | $1/K_d$ ($\mu M^{-1}$) | $-\Delta H$ (kJ $mol^{-1}$) | $T\Delta S$ (kJ $mol^{-1}$)[d] |
|---|---|---|---|---|---|---|
| P1F0 | S | 29_24_0 | 0.6 ± 0.1 | 1.6 ± 0.2 | 30.0 ± 0.9 | 3.3 ± 0.9 |
| P2F0 | S | 30_25_0 | 0.7 ± 0.1 | 1.5 ± 0.2 | 30.2 ± 1.1 | 3.3 ± 1.1 |
| P3F0 | S | 31_26_0 | 0.7 ± 0.1 | 1.5 ± 0.2 | 34.0 ± 1.4 | -0.1 ± 1.4 |
| P4F0 | S | 32_27_0 | 0.6 ± 0.1 | 1.6 ± 0.3 | 31.9 ± 1.3 | 2.1 ± 1.3 |
| P5F0 | S | 33_28_0 | 0.7 ± 0.1 | 1.5 ± 0.3 | 28.5 ± 1.4 | 5.3 ± 1.4 |
| P6F0 | S | 34_29_0 | 0.8 ± 0.1 | 1.3 ± 0.2 | 16.4 ± 1.2 | 16.6 ± 1.2 |
| P7F0 | S | 35_30_0 | 0.8 ± 0.1 | 1.2 ± 0.2 | 19.8 ± 1.0 | 13.4 ± 1.0 |
| P8F0 | S | 36_31_0 | 0.7 ± 0.1 | 1.4 ± 0.2 | 20.2 ± 0.8 | 12.7 ± 0.9 |
| P9F0 | S | 37_32_0 | 0.7 ± 0.1 | 1.4 ± 0.2 | 15.5 ± 0.9 | 17.6 ± 0.9 |

| Annotation | Variant | Composition[c] | $K_d$ (μM) | $1/K_d$ ($\mu M^{-1}$) | $-\Delta H$ (kJ $mol^{-1}$) | $T\Delta S$ (kJ $mol^{-1}$)[d] |
|---|---|---|---|---|---|---|
| P1F1 | S | 29_24_1 | ND | ND | ND | ND |
| P2F1 | S | 30_25_1 | 0.3 ± 0.1 | 3.4 ± 0.4 | 18.7 ± 1.4 | 16.8 ± 1.4 |
| P3F1 | S | 31_26_1 | 0.5 ± 0.1 | 2.1 ± 0.2 | 16.0 ± 0.7 | 21.2 ± 0.7 |
| P4F1 | S | 32_27_1 | 0.6 ± 0.1 | 1.8 ± 0.2 | 24.3 ± 0.8 | 13.6 ± 0.8 |
| P5F1 | S | 33_28_1 | 0.6 ± 0.1 | 1.7 ± 0.3 | 22.5 ± 0.5 | 15.1 ± 0.5 |
| P6F1 | S | 34_29_1 | 0.5 ± 0.1 | 2.1 ± 0.3 | 16.8 ± 0.4 | 20.2 ± 0.4 |
| P7F1 | S | 35_30_1 | 0.5 ± 0.1 | 2.1 ± 0.3 | 15.7 ± 0.6 | 20.1 ± 0.6 |
| P8F1 | S | 36_31_1 | 0.4 ± 0.1 | 2.8 ± 0.2 | 14.1 ± 0.6 | 20.9 ± 0.6 |
| P9F1 | S | 37_32_1 | 0.3 ± 0.1 | 2.9 ± 0.4 | 17.2 ± 0.8 | 18.7 ± 0.8 |

a. Errors correspond to one standard deviation.
b. ND = Not determined.
c. The compositions of glycan compositions corresponded to the number of Hexose (Hex), N-acetylhexosamine (HexNAc), Fucose (Fuc).
d. Temperature ($T$) is 298.15 K.

### Figures

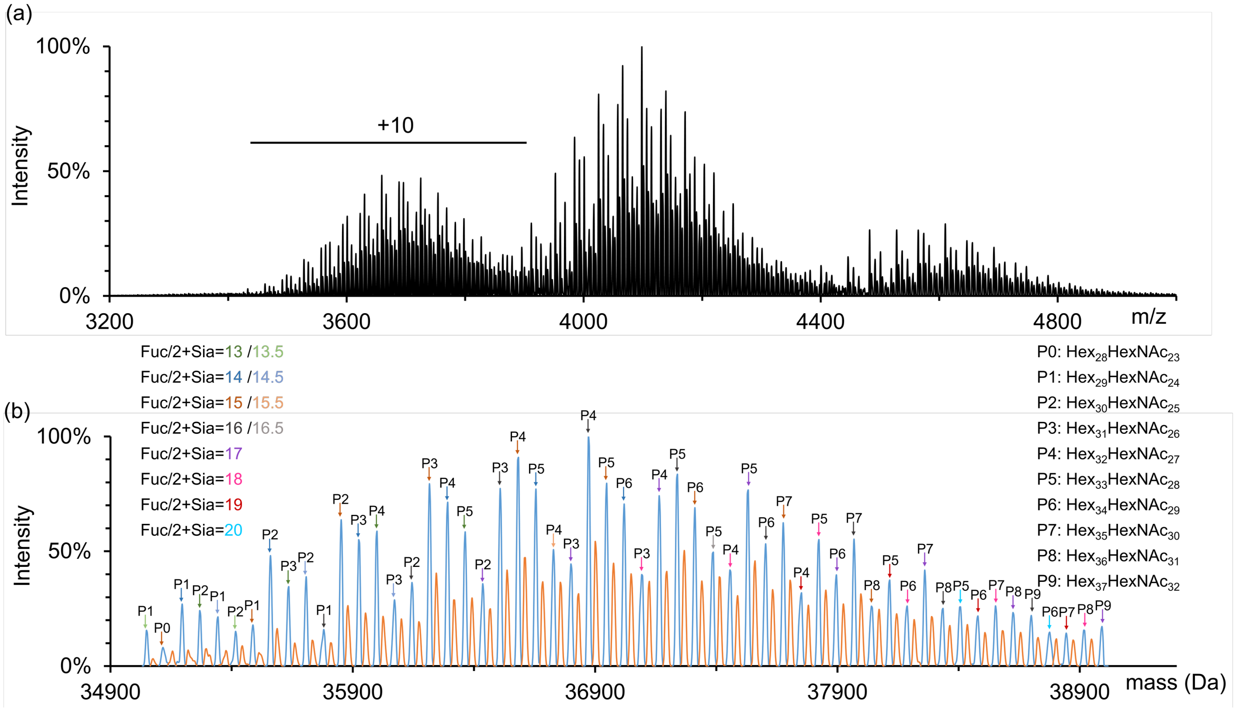


**Figure S1.** Representative (a) native mass spectrum and (b) corresponding zero-charge deconvoluted mass spectrum of AGP (3 μM) in 200 mM ammonium acetate (pH 7.6) containing 1% DMSO, acquired in positive-ion mode. Peak families sharing the same numbers of hexose (Hex) and *N*-acetylhexosamine (HexNAc) residues are labelled P0–P9. Colored arrows indicate the numbers of *N*-acetylneuraminic acid (Neu5Ac) and fucose (Fuc) residues associated with each peak.

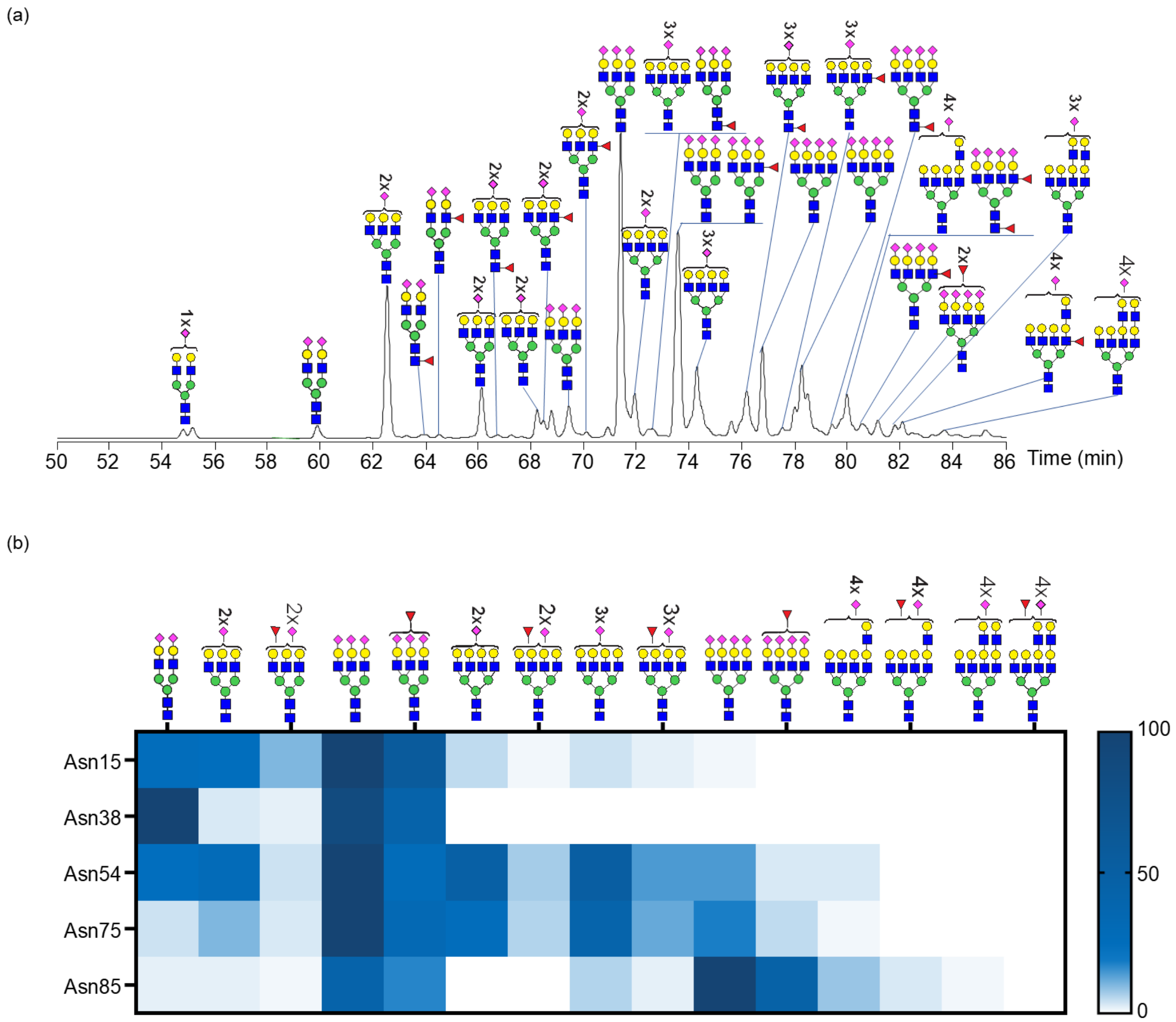


**Figure S2.** (a) HILIC-UHPLC chromatogram of ProA-labelled *N*-glycans released from AGP, obtained using fluorescence detection. (b) Heatmap showing the relative abundances of glycans at the five *N*-glycosylation sites of AGP, calculated from three replicate LC-MS/MS experiments.

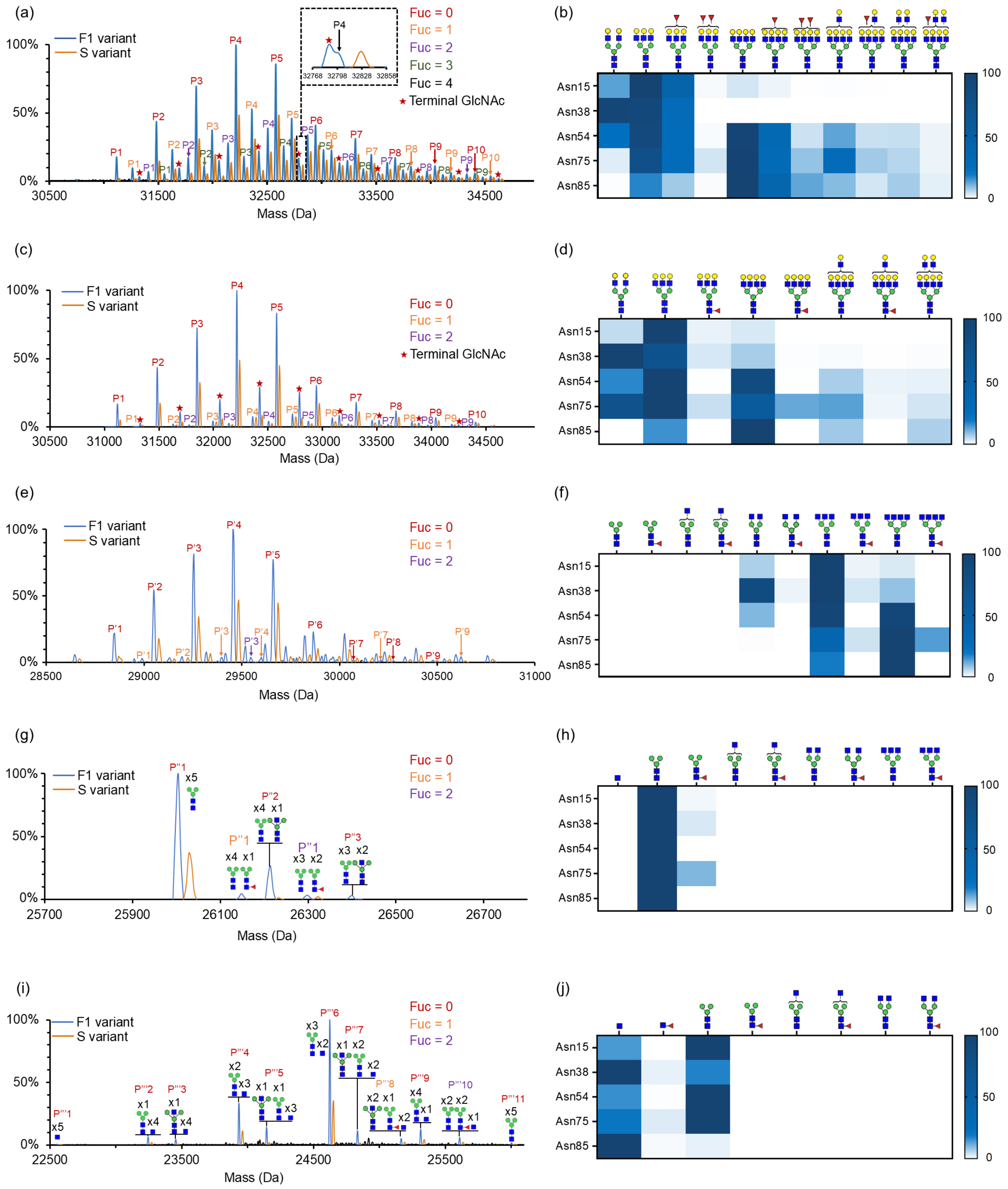

**Figure S3.** Characterization of enzymatically remodeled AGP samples. Representative zero-charge deconvoluted native mass spectra and corresponding heatmaps of site-specific glycan relative abundances are shown for (a, b) asialo-AGP, (c, d) AD-asialo-AGP, (e, f) AG-AD-asialo-AGP, (g, h) TG-AG-AD-asialo-AGP, and (i, j) CG-TG-AG-AD-asialo-AGP. Mass spectral peaks were assigned to the glycoprotein compositions listed in Tables S3–S7, respectively.

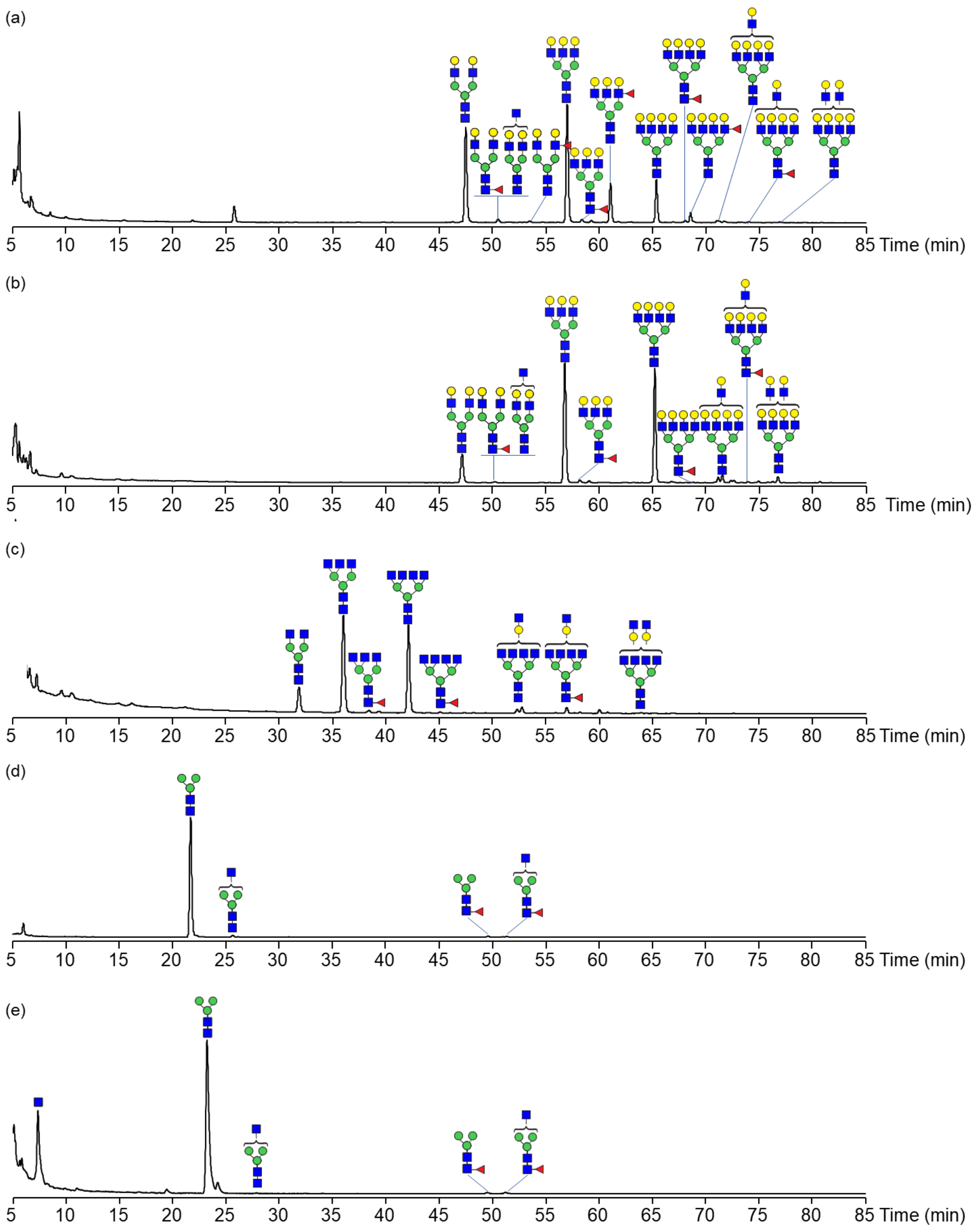


**Figure S4.** HILIC-UHPLC chromatograms of ProA-labelled *N*-glycans released from (a) asialo-AGP, (b) AD-asialo-AGP, (c) AG-AD-asialo-AGP, (d) TG-AG-AD-asialo-AGP, and (e) CG-TG-AG-AD-asialo-AGP, obtained using fluorescence detection.

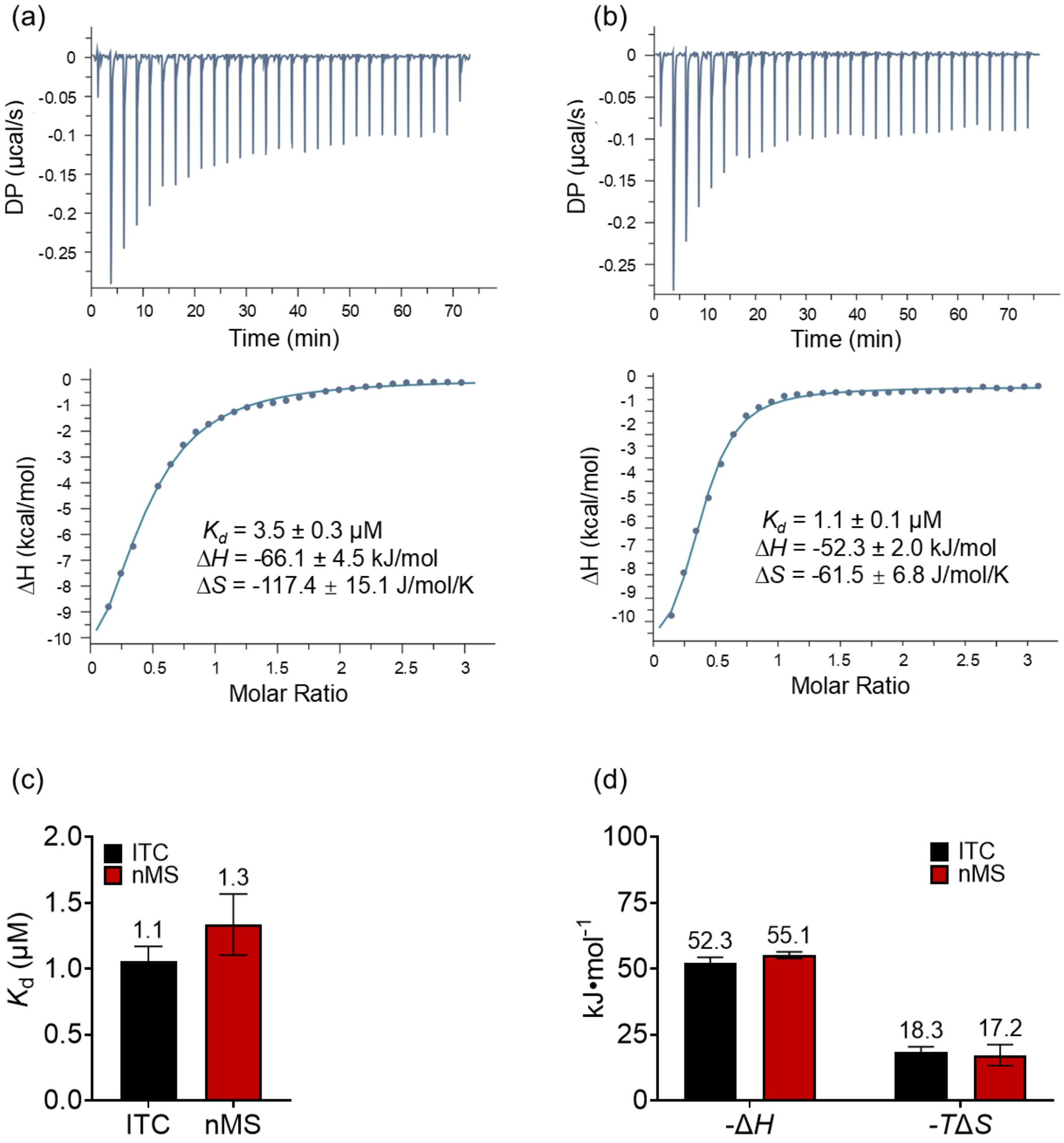


**Figure S5.** Comparison of warfarin-binding affinities and thermodynamic parameters determined by vT-nMS and ITC. The ITC data for warfarin binding to (a) AGP and (b) asialo-AGP were acquired at 25 °C in 200 mM ammonium acetate (pH 7.6) using protein and warfarin concentrations of 15 µM and 300 µM, respectively. Comparison of (c) dissociation constants and (d) binding thermodynamic parameters for asialo-AGP, using population-weighted mean values from vT-nMS and apparent values from ITC.

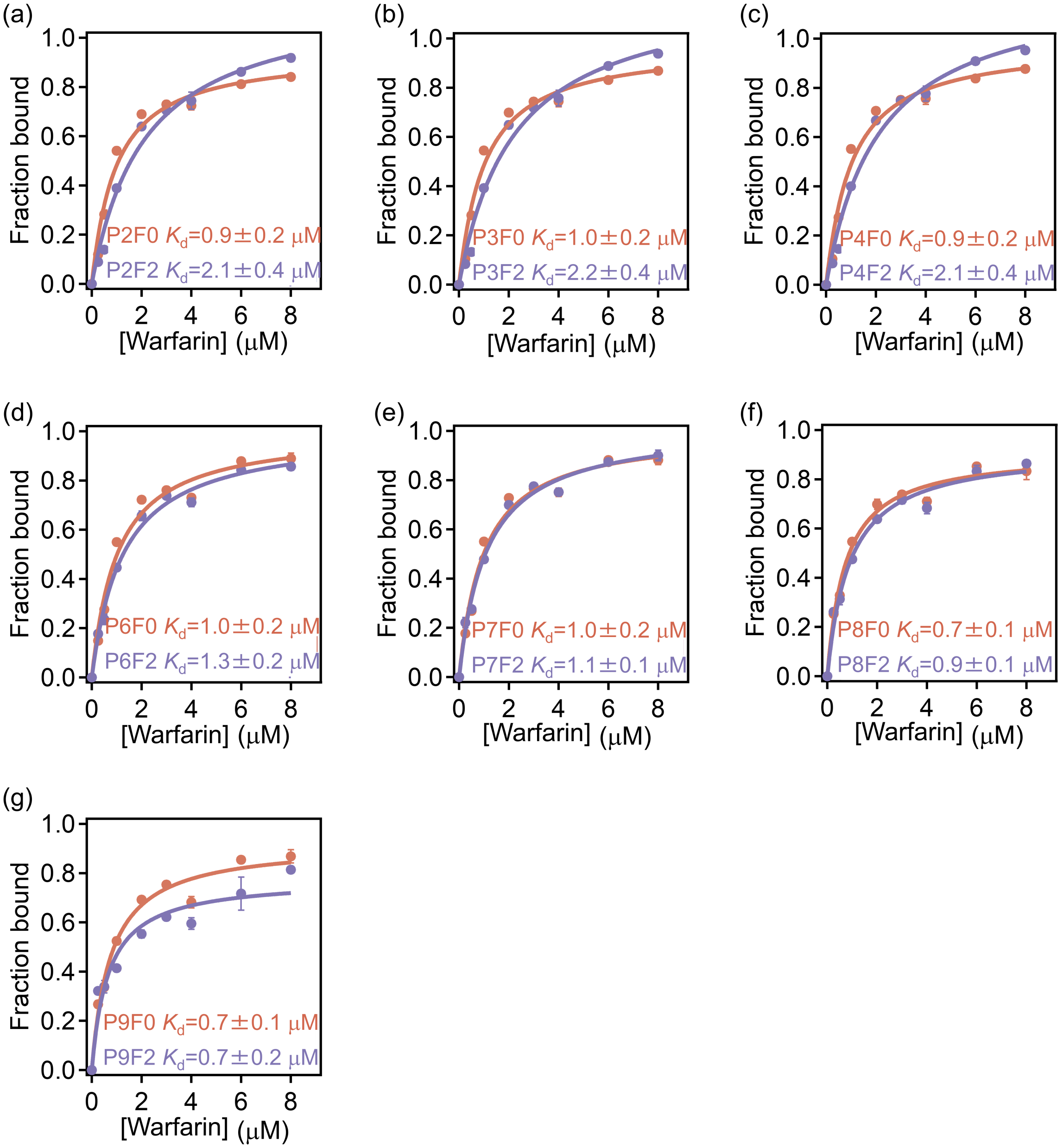


**Figure S6.** Warfarin-binding titration curves for unfucosylated (F0, orange) and difucosylated (F2, purple) glycan compositions of the asialo-AGP F1 variant: (a) P2, (b) P3, (c) P4, (d) P6, (e) P7, (f) P8, and (g) P9. Symbols show experimentally observed bound fractions, and solid curves show fits to eq S8, accounting for in-source dissociation. Fitted dissociation constants ($K_d$) are indicated.

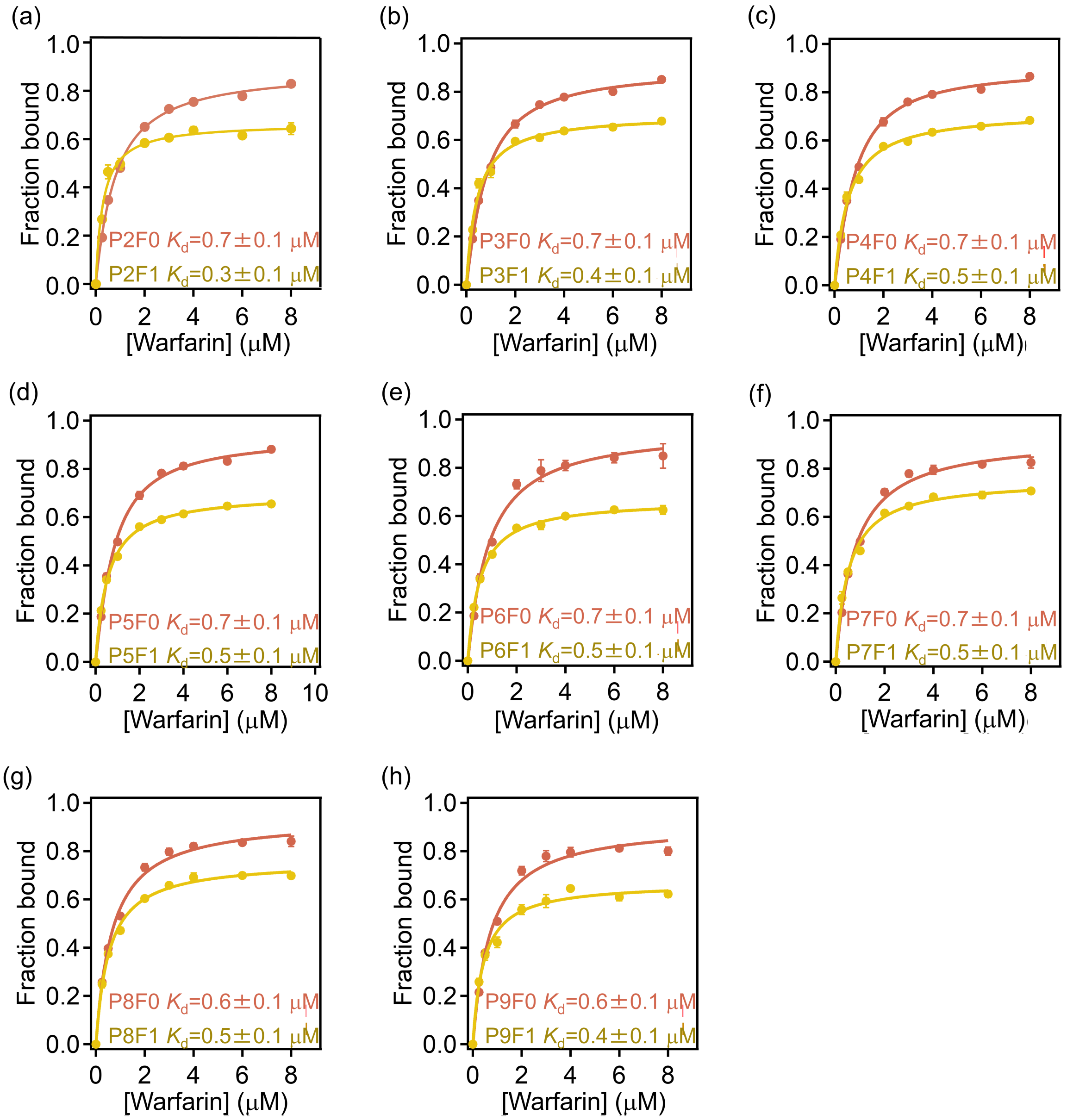


**Figure S7.** Warfarin-binding titration curves for unfucosylated (F0, orange) and core-monofucosylated (F1, yellow) glycan compositions of the AD-asialo-AGP F1 variant: (a) P2, (b) P3, (c) P4, (d) P5, (e) P6, (f) P7, (g) P8, and (h) P9. Symbols show experimentally observed bound fractions, and solid curves show fits to eq S8, accounting for in-source dissociation. Fitted dissociation constants ($K_d$) are indicated.

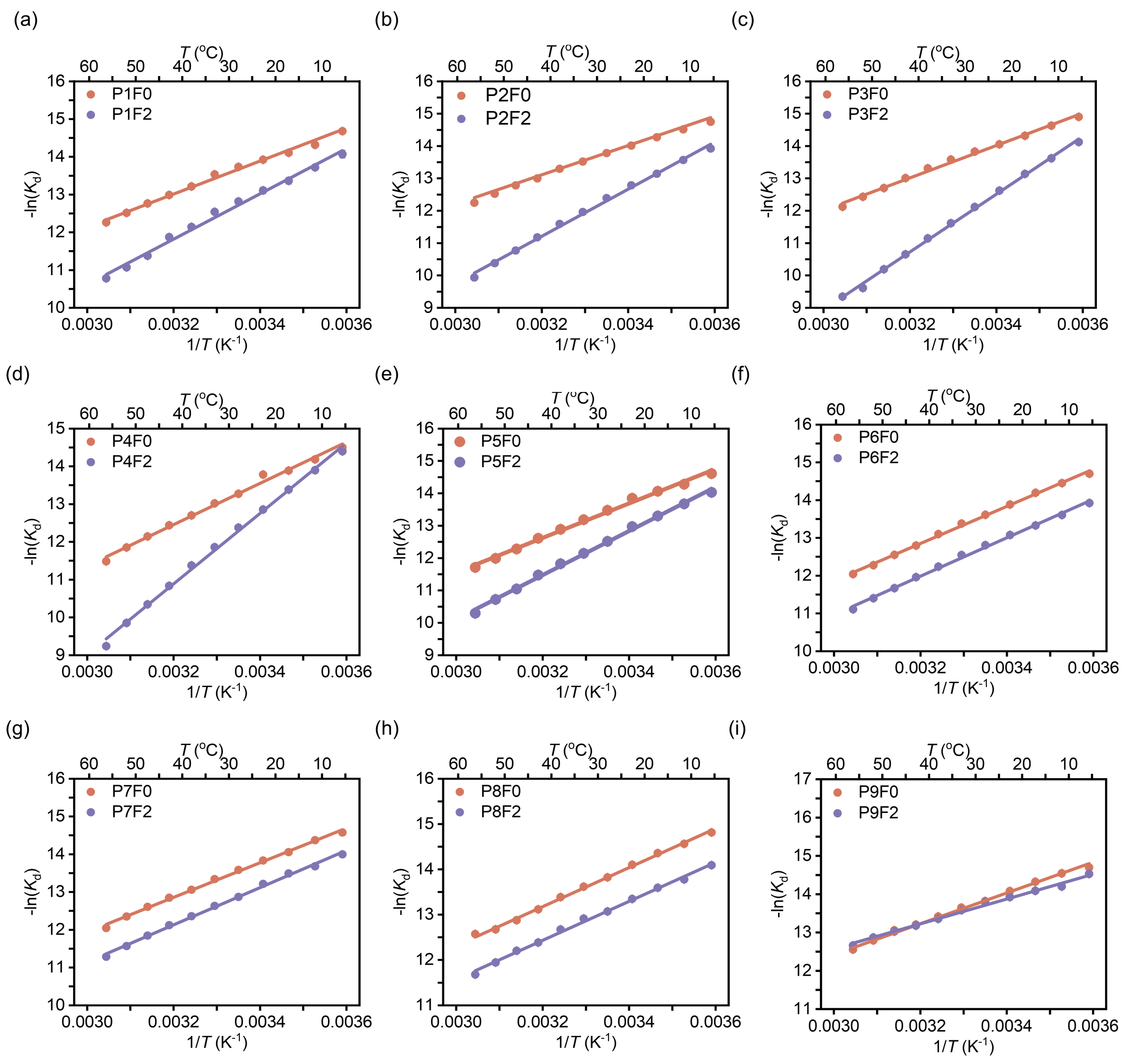


**Figure S8.** Van't Hoff plots for warfarin binding to unfucosylated and difucosylated glycan compositions of the asialo-AGP F1 variant, determined by vT-nMS.

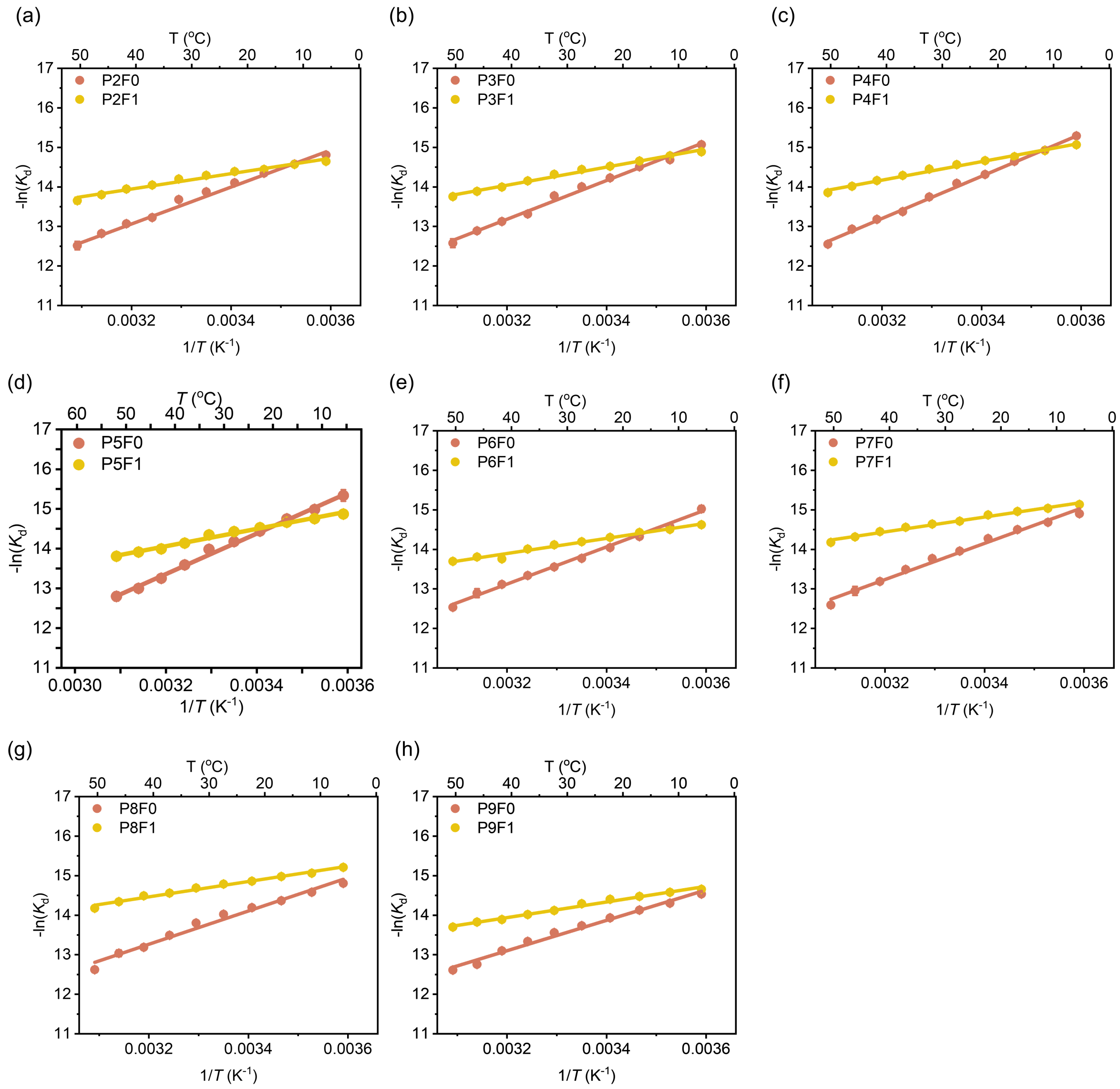


**Figure S9.** Van't Hoff plots for warfarin binding to unfucosylated and core-monofucosylated glycan compositions of the AD-asialo-AGP F1 variant, determined by vT-nMS.

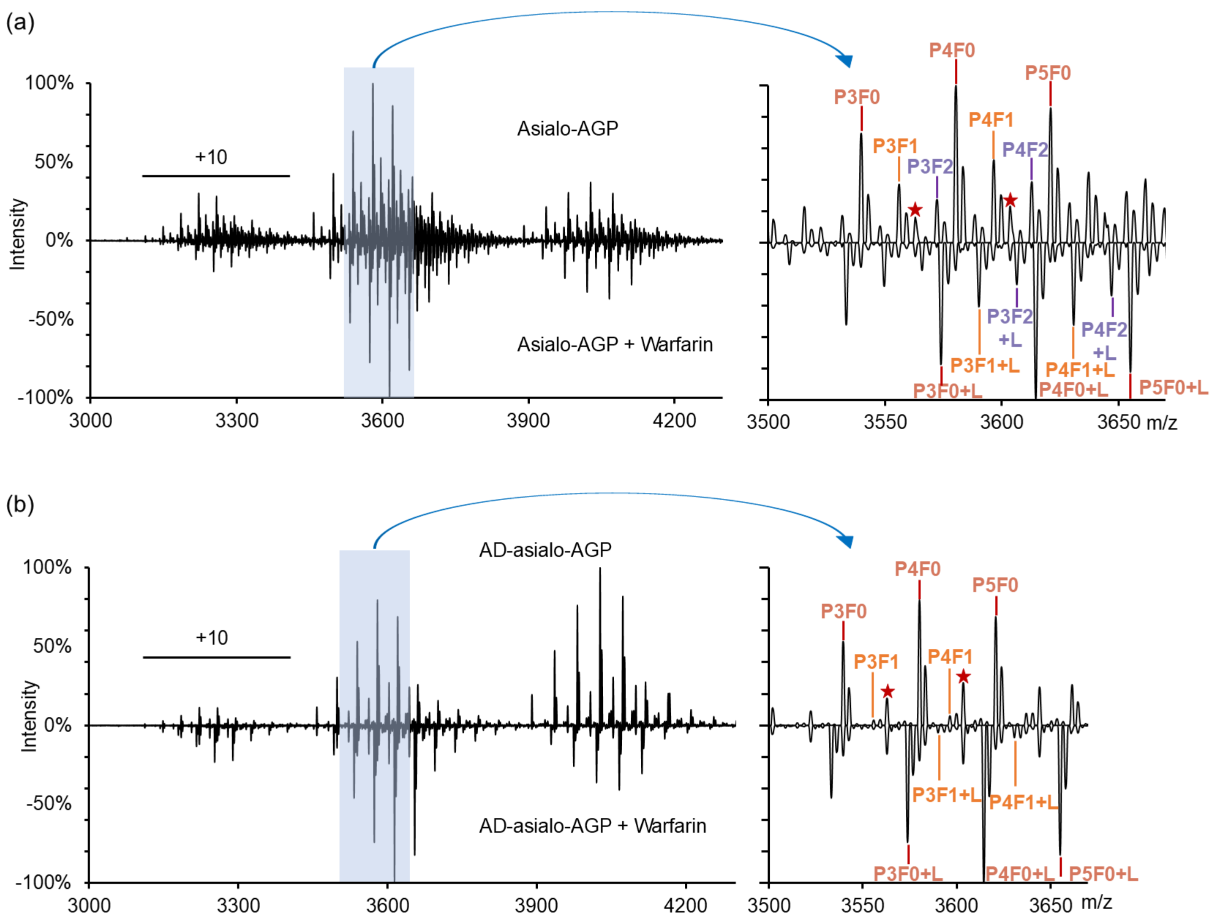


**Figure S10.** Representative native mass spectra of (a) asialo-AGP and (b) AD-asialo-AGP in the absence (upper traces) and presence (inverted lower traces) of warfarin. Expanded views of the shaded regions are shown on the right. Red stars indicate peaks assigned to compositions containing a bisecting GlcNAc residue; corresponding warfarin-bound complexes were not detected. Warfarin-bound compositions are labelled "+L."

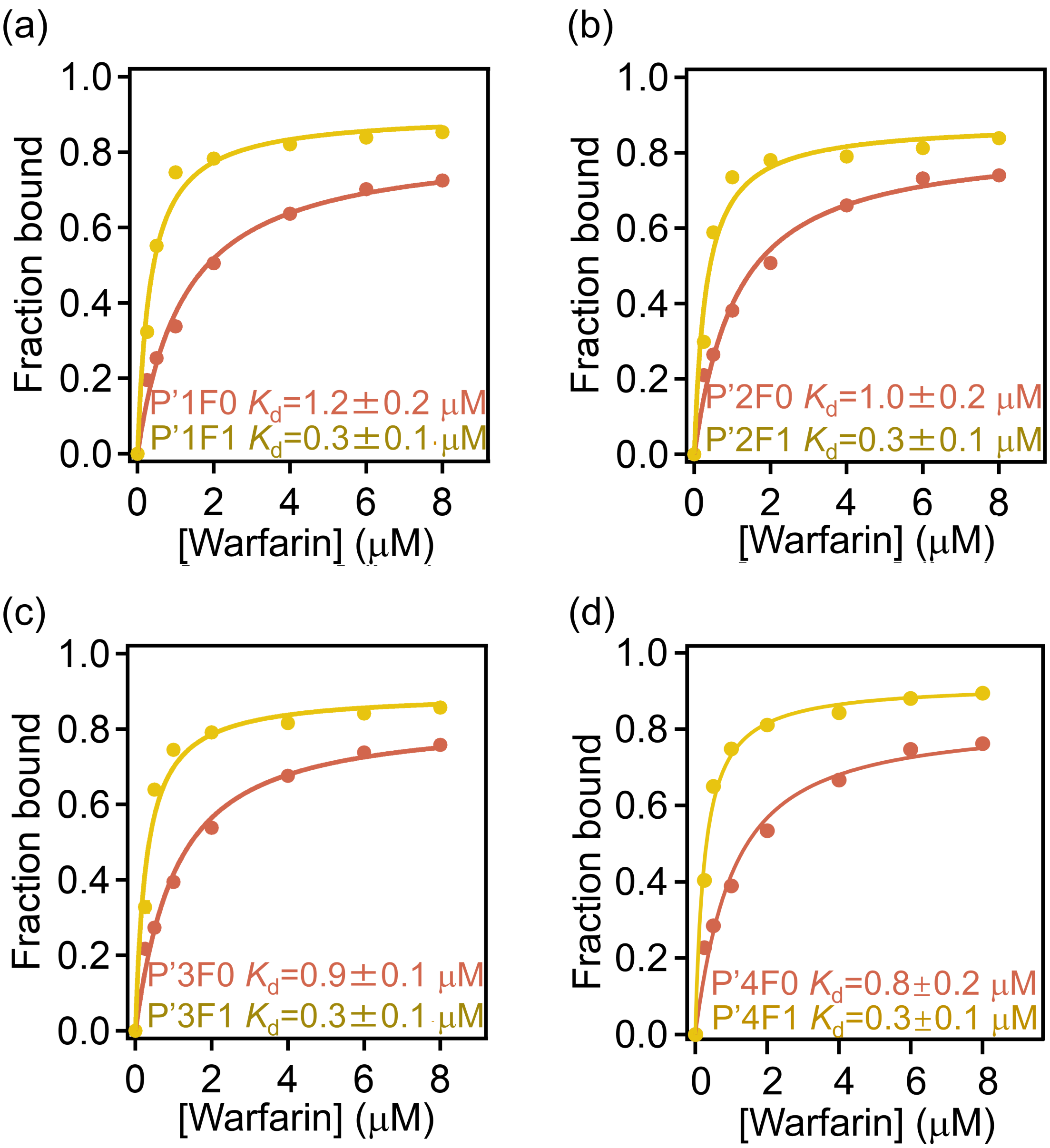


**Figure S11.** Warfarin-binding titration curves for unfucosylated (F0, orange) and core-monofucosylated (F1, yellow) glycan compositions of the AG-AD-asialo-AGP F1 variant: (a) P′1, (b) P′2, (c) P′3, and (d) P′4. Symbols show experimentally observed bound fractions, and solid curves show fits to eq S8, accounting for in-source dissociation. Fitted dissociation constants ($K_d$) are indicated.

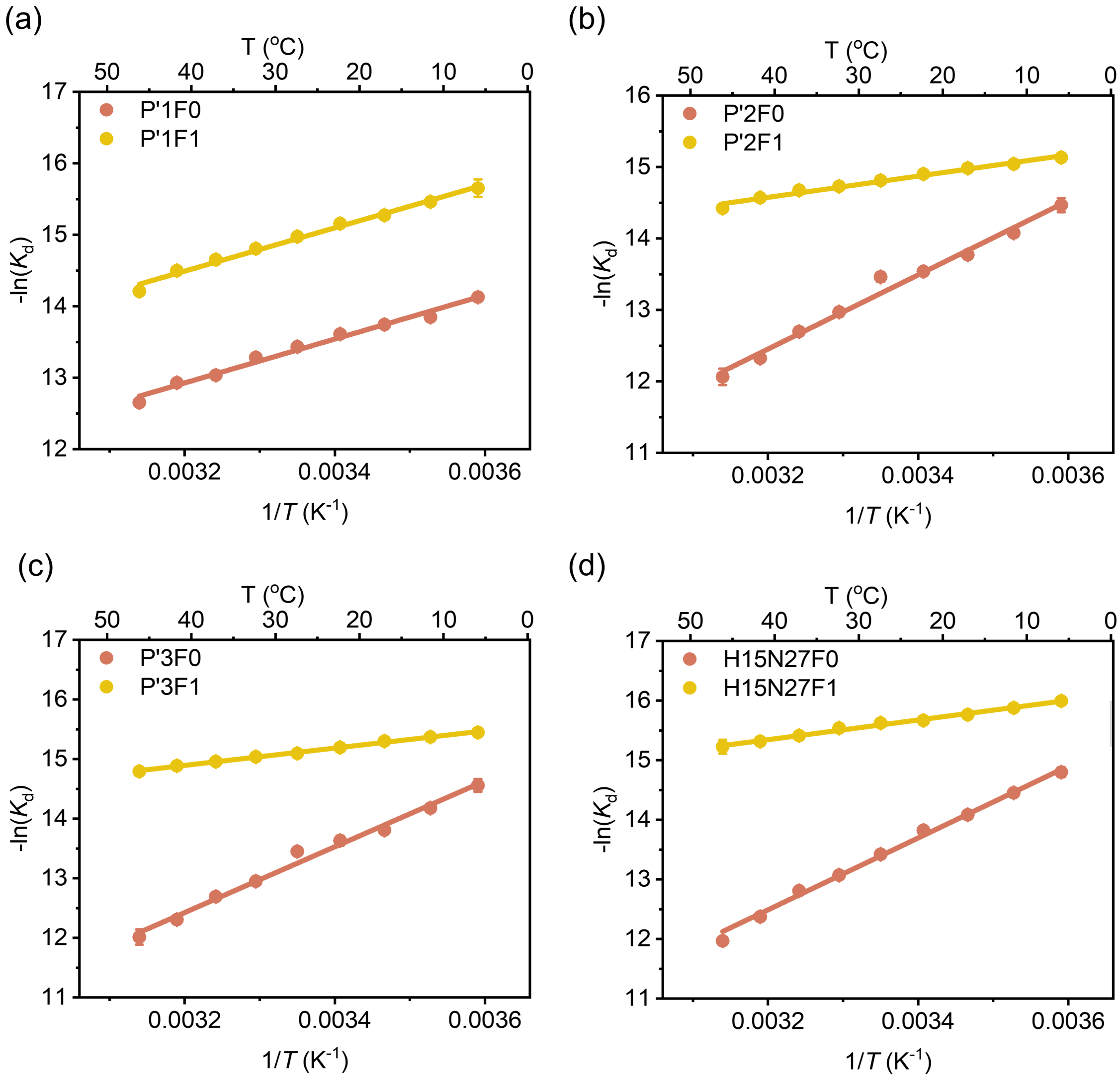


**Figure S12.** Van't Hoff plots for warfarin binding to unfucosylated and core-monofucosylated glycan compositions of the AG-AD-asialo-AGP F1 variant, determined by vT-nMS.

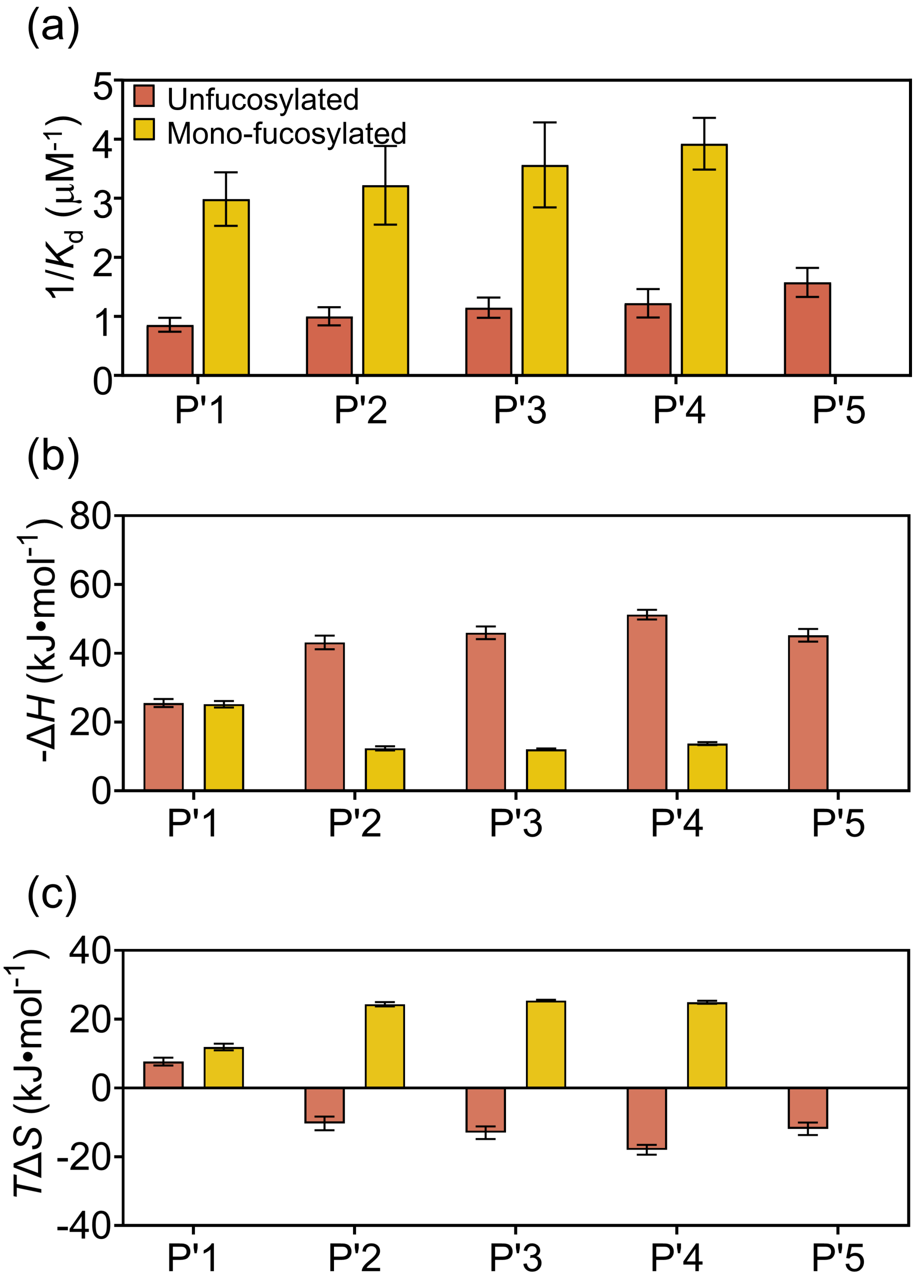


**Figure S13.** Composition-dependent affinity and thermodynamics of warfarin binding to the AG-AD-asialo-AGP F1 variant. (a) Association constants ($1/K_d$), (b) negative binding enthalpies ($-\Delta H$), and (c) entropic contributions ($T\Delta S$) at 298.15 K for unfucosylated and core-monofucosylated compositions, determined by vT-nMS. Error bars indicate ±1 standard deviation from three replicate measurements.

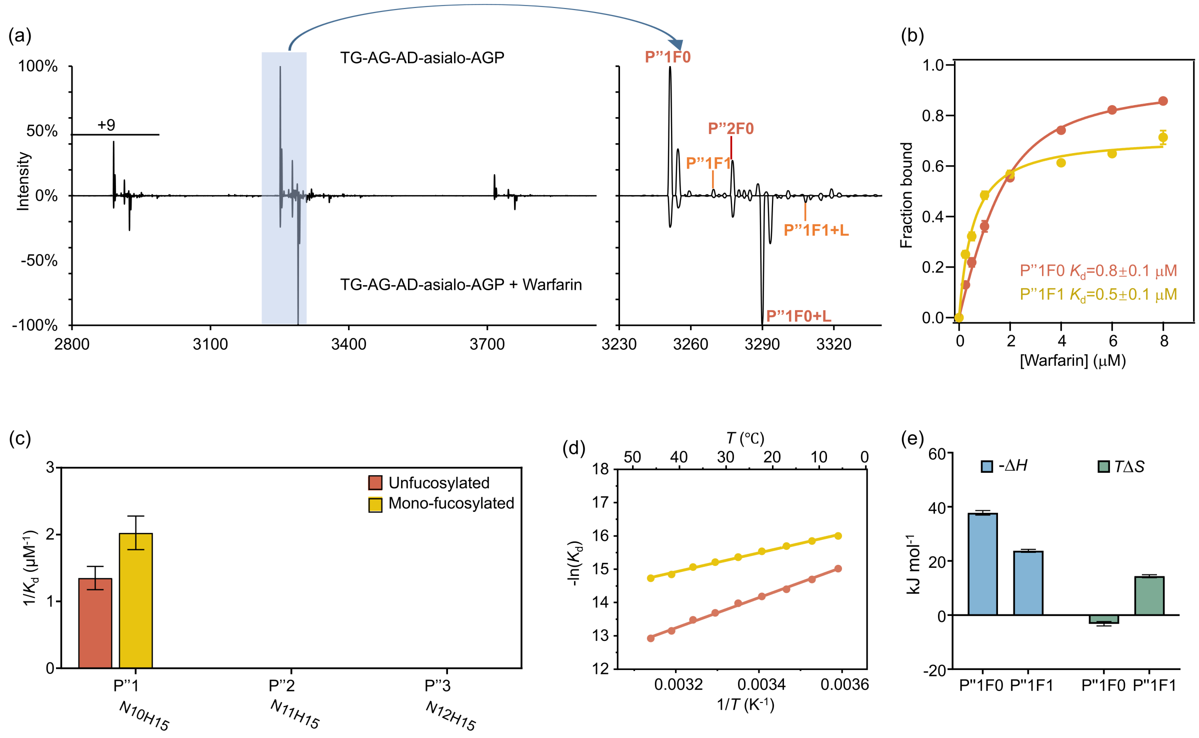


**Figure S14.** Composition-dependent warfarin binding and thermodynamics of the TG-AG-AD-asialo-AGP F1 variant. (a) Representative native mass spectra of TG-AG-AD-asialo-AGP (3 μM) in 200 mM ammonium acetate (pH 7.6) containing 1% DMSO, in the absence (upper trace) and presence (inverted lower trace) of warfarin (6 μM). An expanded view of the shaded region is shown on the right; warfarin-bound compositions are labelled "+L." (b) Warfarin-binding titration curves for unfucosylated and core-monofucosylated compositions. (c) Association constants ($1/K_d$) at 298.15 K. (d) Van't Hoff plots determined by vT-nMS. (e) Negative binding enthalpies ($-\Delta H$) and entropic contributions ($T\Delta S$, 298.15 K) for the compositions shown in (b). Error bars indicate ±1 standard deviation from three replicate measurements.

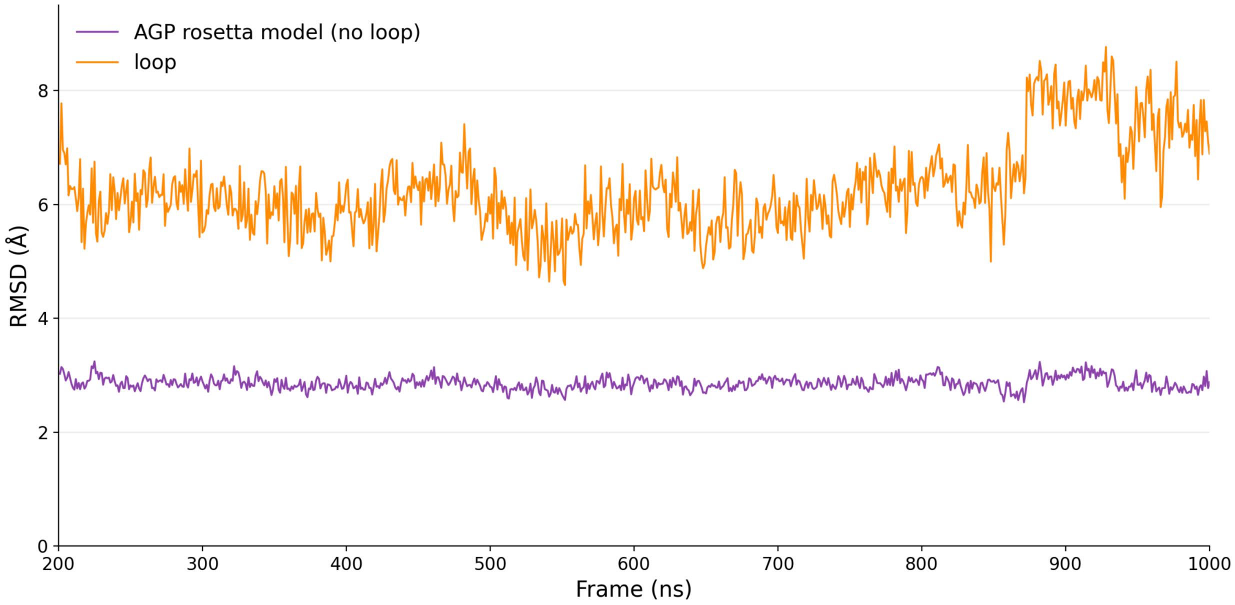


**Figure S15.** Backbone RMSD along the GaMD trajectory of the RosettaFold-modeled AGP system (first 200 ns excluded), after superposition on the protein backbone (excluding the loop). Purple: RMSD of the protein (loop excluded, resid 36 to 42). Orange: RMSD of the RosettaFold-modeled loop, measured after the same alignment.

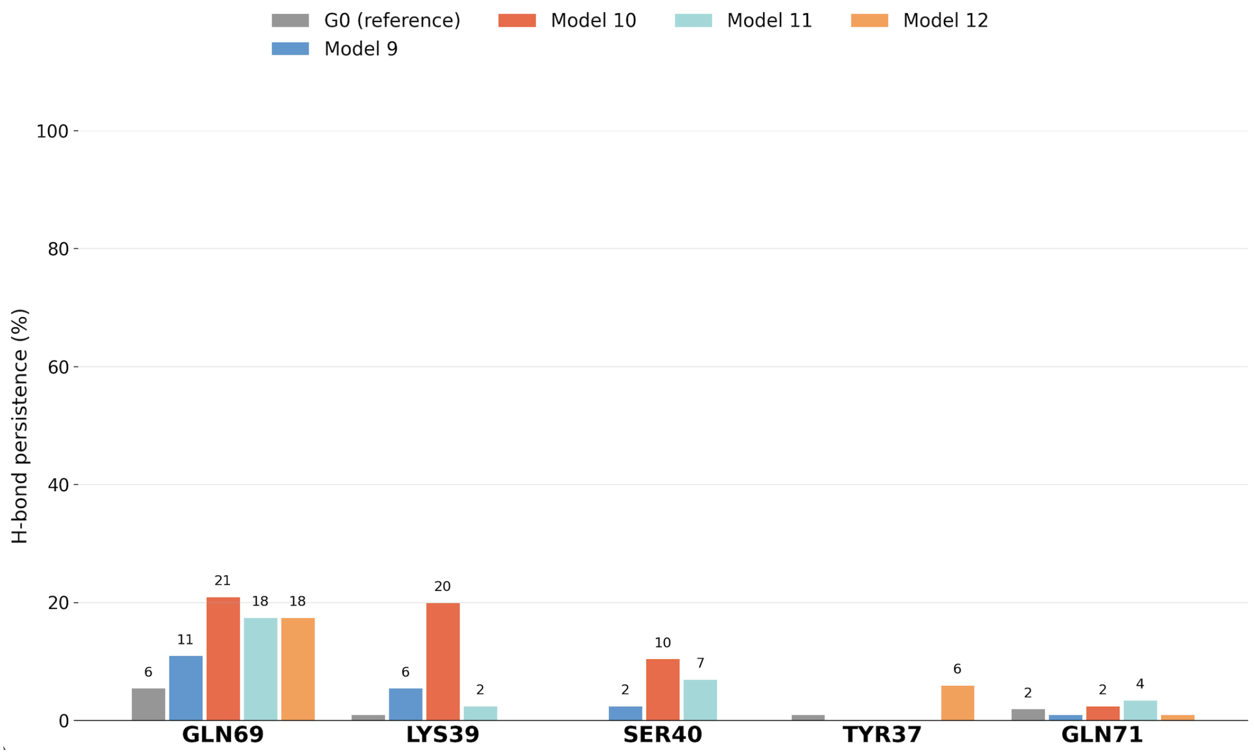


**Figure S16.** Hydrogen-bond persistence between the Asn38 *N*-glycan and residues near the funnel entrance, for the S-variant unfucosylated reference (G0) and antennary-fucosylated glycoforms differing in glycan branching and fucosylated arm at Asn38 (Models 9–12; α1,3 or α1,6 arm, bi- or tri-antennary; see Table S1). Persistence is expressed as the percentage of Re-Glyco Ensemble conformations (200 per glycoform) in which a hydrogen bond was detected between any atom of the Asn38 glycan and the indicated protein residue.

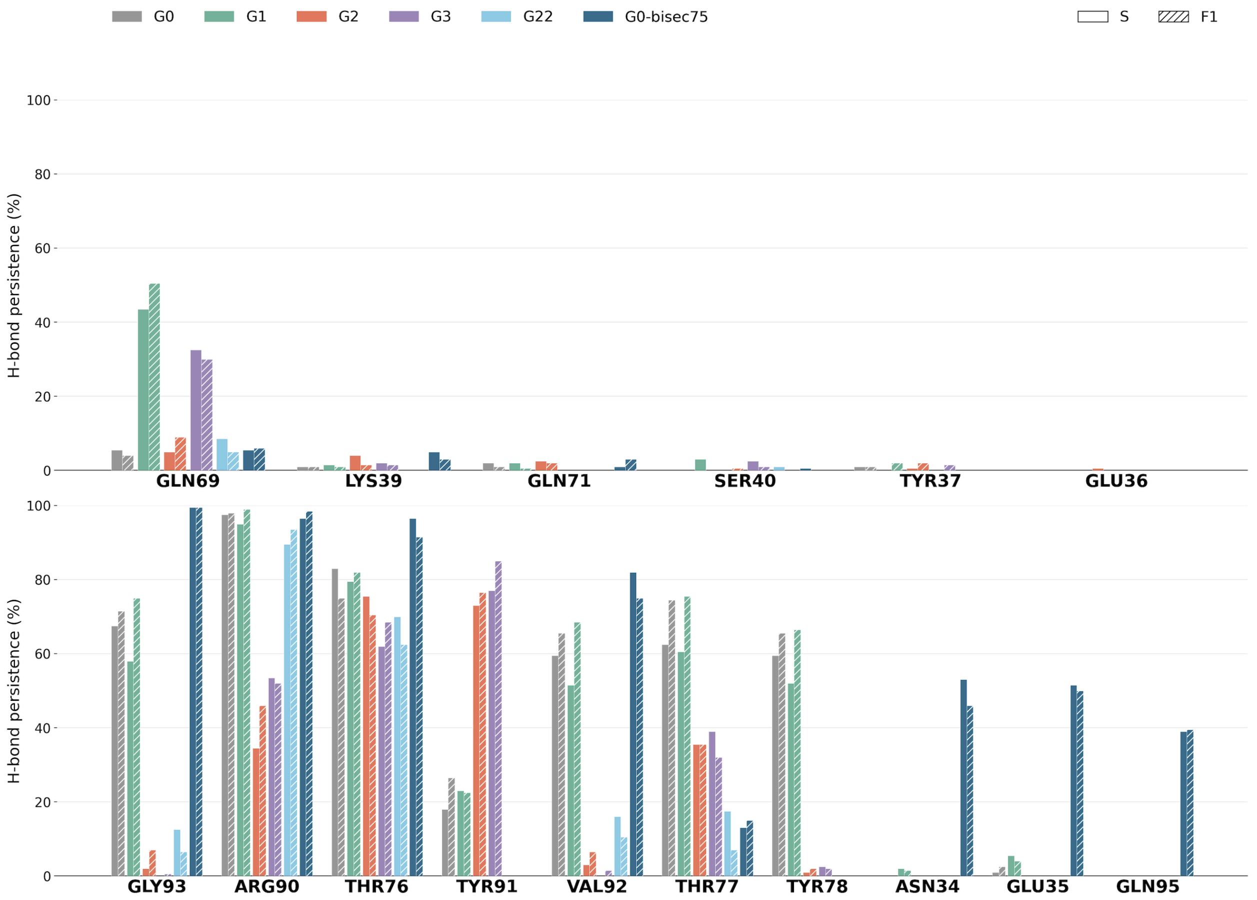


**Figure S17.** Comparison of hydrogen-bond persistence between the S and F1 AGP variants (structural models differing only by the R20Q substitution), for the reference and core-fucosylated/bisecting glycoforms (G0, G1, G2, G3, G22, G0-bisec75). Top panel: interactions between the Asn38 glycan and residues near the funnel entrance; bottom panel: interactions between the Asn75 glycan and residues lining the binding pocket. Persistence is expressed as the percentage of Re-Glyco Ensemble conformations (200 per glycoform) showing a hydrogen bond between the glycan and the indicated protein residue. Solid bars, S variant; hatched bars, F1 variant; colors denote glycoform.
